# Discovery of stereoselective targeted covalent inhibitors of the RAB27-effector protein-protein interaction

**DOI:** 10.64898/2026.05.22.727177

**Authors:** Elena De Vita, Adam M. Thomas, Delia Brustur, Montse Tersa, Rita Petracca, Shradda Vadodaria, Deborah A Briggs, Jack W. Houghton, Thomas Lanyon-Hogg, Gregory B. Craven, Rhodri M. Morgan, Alan Armstrong, David J. Mann, Katharine Lodge, Alistair N. Hume, Ernesto Cota, Edward W. Tate

## Abstract

RAB27A and RAB27B are homologous small GTPases that regulate intracellular vesicle trafficking, orchestrating endocytic and exocytic processes that affect cellular communication, immune responses, and dynamics of the cellular microenvironment. Through their interactions with effector proteins, RAB27A/B play roles in tumor metastasis and chronic inflammation. However, pharmacological modulation of their activity faces challenges typical of small GTPases, including a lack of well-defined pockets outside the conserved GTP binding site, and large RAB27-effector protein-protein interaction (PPI) surfaces. Here, we present the discovery and development of the first cell-active, rationally designed covalent inhibitors of the RAB27-effector interaction, targeting a non-conserved cysteine residue flanking the PPI interface. An electrophile-first biochemical screen led to a novel class of acrylamide covalent inhibitors, and X-ray crystallography structure-guided design led to optimized inhibitors and probes that enantioselectively target RAB27A/B-Cys123 in cells. Potency and selectivity were confirmed through biochemical and cellular assays, including chemical proteomics and phenotype recapitulation in melanocytes alongside a matched inactive enantioprobe control. In contrast, a previously reported compound, Nexinhib-20, was found to be toxic and to exert its activity through non-selective reactivity. This work provides the first toolbox of cell-active chemical probes for RAB27 which can be used in future studies to shed light on the function of this protein and its potential as a therapeutic target.

## Introduction

RAB27A and B are small GTPases belonging to the Ras-related in brain (Rab) family, which encompasses >60 members of the Ras superfamily that control trafficking and fate of intracellular vesicles.^1^ Human RAB27A and B (hereafter referred to as RAB27A/B) share 72% homology, and regulate vesicle trafficking and docking to membranes, with non-redundant roles in vesicle secretion.^2,3^

For example, in melanocytic cells, RAB27A specifically controls the trafficking and secretion of melanosomes, which are intracellular vesicles that shuttle the pigment melanin into keratinocytic skin and hair cells.^4^

The interaction of Rab proteins with membranes is directed by C-terminal geranylgeranylation by Rab Geranyl-Geranyl Transferase (GGTase), and inhibited by Rab GDP Dissociation Inhibitor (GDI), which complexes to geranylgeranylated GDP-bound Rab proteins in the cytosol.^5^ RAB27A/B display low intrinsic GTPase activity, and cycling between active (GTP-bound) and inactive (GDP-bound) states is tightly regulated by GTPase Activating Proteins (GAPs) and Guanosine Nucleotide Exchange Factors (GEFs). When bound to GTP, RAB27A/B docks to intracellular membranes such as melanosomes or multivesicular bodies, and adopts a more rigid conformation that favors binding to specific effector proteins (Figure 1A). The eleven known RAB27A/B effectors belong to three distinct families: synaptotagmin-like proteins (Slp), containing tandem C2 Ca^2+^-binding motifs; the Slac2 family, lacking these C2 motifs; and Munc13-4.^6^ Recently, SPIRE-type actin nucleators were also identified to work as RAB27A/B effectors.^7^ RAB27A/B-effector protein-protein interactions (PPIs) feature an essential binding interface corresponding to a shallow pocket that hosts a lipophilic effector residue pair, typically tryptophan (W) and phenylalanine (F), also known as the ‘WF pocket’ (Figure 1B)^8–12^. Two cysteine residues flank the WF pocket in RAB27A/B, Cys123 and Cys188, which are non-conserved across the rest of the Rab protein family.^13^ In a previous study, we validated covalent fragments binding to both cysteines by X-ray crystallography (**A01** and **B01**, Figure 1C), demonstrating their ligandability.^13^

**Figure 1.**
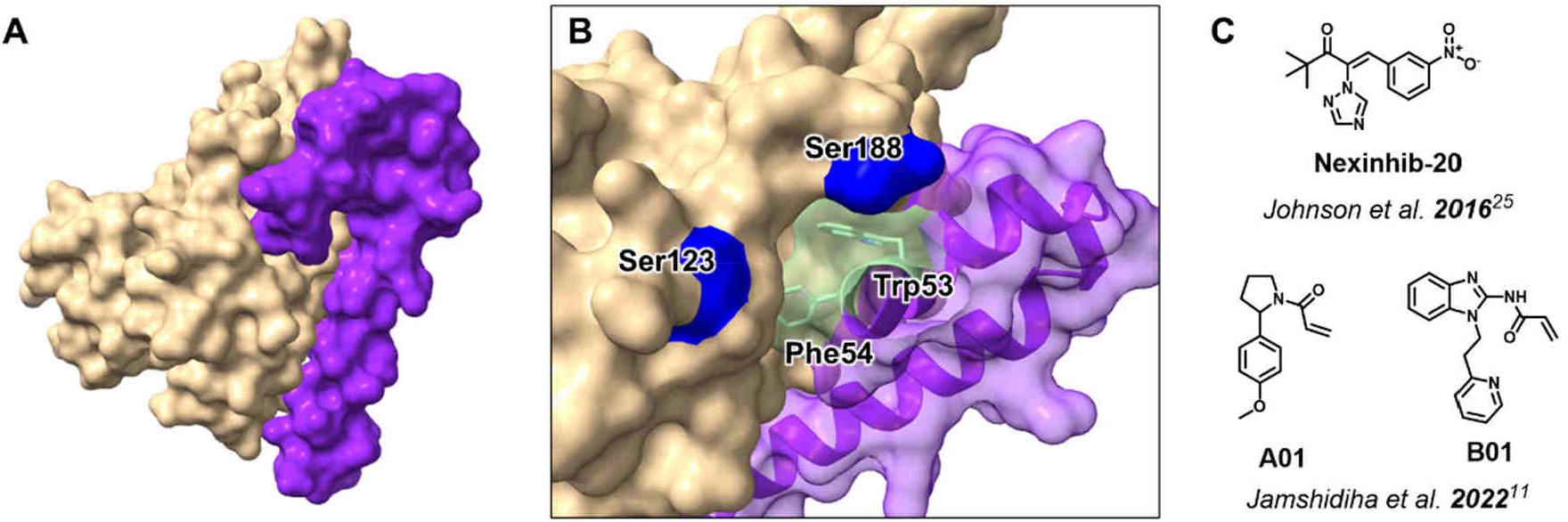
A) Mouse RAB27A (sand) in complex with truncated human effector SLP2-A (purple) from PDB:3BC1; B) WF pocket interacting residues (green) and flanking non-conserved cysteines mutated to serine (blue); C) Previously reported ligands for RAB27A/B: **Nexinhib-20**^25^, **A01**^11^ and **B01**^11^.

Due to its role in regulating exocytotic processes, RAB27A/B has been implicated in the remodeling of the tumor microenvironment across multiple types of malignancies,^14^ including breast,^15,16^ pancreatic^17,18^ and colorectal cancers.^19,20^ For example, RAB27A knock-down significantly improves anticancer efficacy of immune-checkpoint inhibitor therapies by reducing exosomal PD-L1, which otherwise impairs T-cell mediated anti-tumor surveillance and dampens antibody therapies targeting PD-L1.^21^ These findings suggest that targeting RAB27A/B could disrupt the tumor microenvironment and potentially prevent the establishment of pre-metastatic niches.^22^ Furthermore, a recent study reported that targeting RAB27A/B could be exploited to facilitate mobilization of immune cells to the brain tissue in brain tumours.^23^ RAB27A has also been found to support the budding process of virus release for example for Influenza virus, via effectors SYTL1 and SYTL4, suggesting a potential application of RAB27A/B inhibitors in antiviral therapy.^24^

Loss-of-function mutations of RAB27A are viable in humans, but result in an autosomal-recessive disorder known as Griscelli type II syndrome, which is characterized by partial albinism and haemophagocytic lymphohistocytosis (HLH).^25^ If identified early, HLH can be treated by bone marrow transplant, while knock out (KO) of RAB27A/B in adult mice is viable with a milder phenotype.^26^ Together, these observations suggest that transient chemical inhibition of RAB27A/B function in adult patients is likely to carry a low risk of mechanism-based toxicity, motivating the identification of selective inhibitors.

The current lack of high-quality chemical probes hampers efforts to validate RAB27A/B as an actionable target in disease. Although widely used to study RAB27A/B function, Nexinhib-20, originally reported as a selective, noncovalent RAB27A inhibitor,^27^ raises concerns in our eyes due to the presence of pan-assay interferers (PAINs) functional groups, such as a nitro group and a highly conjugated α,β-unsaturated ketone (Figure 1C).

Here we report the discovery and validation of a novel chemical series of covalent inhibitors that target RAB27A/B-Cys123. Through a range of orthogonal biochemical and cellular assays we demonstrate that optimized inhibitors and probes engage endogenous RAB27A/B in different cell lines and phenocopy RAB27A KO in primary melanocytes through disruption of key effector PPIs. These compounds represent the first example of validated cellular probes for this challenging PPI target, alongside matched enantiomeric negative controls.

## Results and Discussion

### Identification of a synthetically tractable acrylamide fragment with enhanced reactivity for RAB27A-Cys123

To enable structure-guided discovery and validation of novel RAB27A/B chemical probes, we recently reported a RAB27A recombinant construct termed fRAB27A which is GppNHp-bound (Q78L locked, stopping intrinsic GTP hydrolysis), lacks the hypervariable prenylated C-terminal region and is N-terminally fused to a truncated section of the RAB27A/B effector SLP2-A, allowing facile crystallization (Figure 2A,B).^13^ RAB27A and B are highly homologous at the WF pocket, hence we selected RAB27A as the primary model for biochemical studies. To study modification of a single exposed cysteine, we produced fRAB27A-Cys123 and fRAB27A-Cys188 recombinant constructs, where the number reflects which of the two cysteines is present, while the other cysteine was mutated to a serine (Figure 2A). Following a pilot screen we identified two fragments (**A01** and **B01**, Figure 1C) binding at either cysteine, which were validated by X-ray crystallography using our fusion constructs (Figure 2B).^13^ However, these scaffolds were found unsuitable for further development into specific RAB27A/B probes.

**Figure 2.**
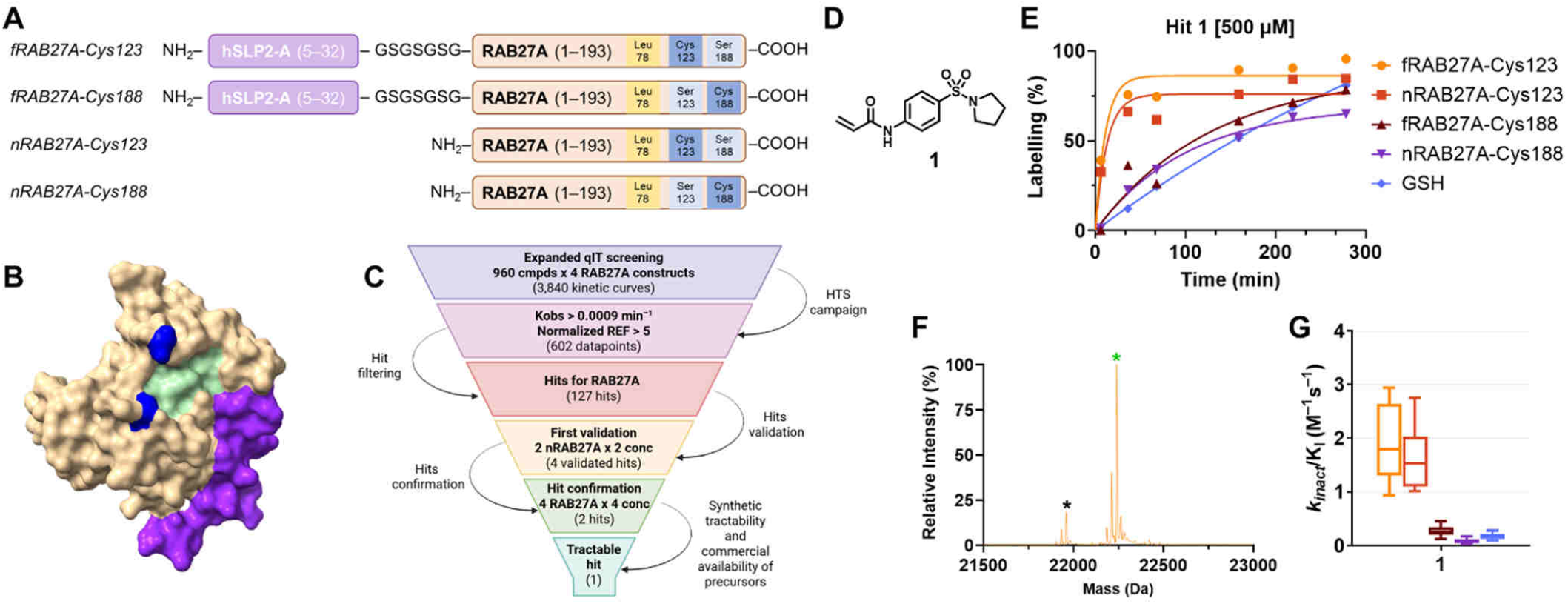
A) Main RAB27A recombinant constructs used in this work with introduced point mutations including present Cys;^11^ B) Fusion construct of RAB27A (PDB:7OPP) highlighting the available WF pocket space (green) and the flanking cysteine residues (blue), both mutated to serine in this structure;C) Covalent fragment screening cascade; D) Chemical structure of hit fragment **1**; E) Kinetic covalent modification curves for all thiols determined via the qIT assay, normalized as percentage labeling over time; F) Intact protein mass spectrometry analysis of nRAB27A-Cys123 (black asterisk, MW: 21961 Da) following incubation with **1** (500 µM, 1 h, rt) showing expected covalent adduct formation (green asterisk, MW: 22241 Da); G) k_inact_/K_i_ values calculated for **1** from at least 10 independent experiments (for K_obs_ graphs, see figure S2).

We performed an expanded screen using an in-house library of 960 acrylamide fragments with the quantitative Irreversible Tethering (qIT) assay, which measures engagement of a single exposed thiol in the target protein (Figure 2C).^28^ We screened fusion constructs fRAB27A-Cys123 and fRAB27A-Cys188, and non-fusion constructs nRAB27A-Cys123 and nRAB27A-Cys188 lacking the fused sequence of the truncated effector (Figure 2A). Furthermore, we screened the library against glutathione (GSH) as a filter for hyperreactive fragments. Our aim was to identify covalent fragments with enhanced and selective reactivity for either RAB27A-Cys123 or Cys188 across both the corresponding fusion and non-fusion constructs, minimizing the risk of false positives whilst preserving the capacity to crystalize hits in the fusion construct.

Library members were incubated at 500 µM with each thiol-bearing protein construct or GSH (5 µM) for 24 h, and samples were taken for qIT analysis at regular time points, leading to 3,840 kinetic curves across four protein constructs. These data were processed to estimate *k*_*obs*_ for each construct and filtered by various criteria, including >5-fold enhanced *k*_*obs*_ over GSH (Figure 2C, see SI methods for details). Four compounds were selected for validation in dose-response analyses against all thiols (fRAB27A-Cys123, fRAB27A-Cys188, nRAB27A-Cys123, nRAB27A-Cys188, and GSH). Following assessment of synthetic tractability, we selected hit compound **1** (**IMP-1704**, Figure 2D) which showed enhanced *k*_obs_ for RAB27A-Cys123 (Figure 2E) and single site modification by intact protein mass spectrometry (Figure 2F). *k*_inact_/*K*_i_ analysis showed an 18-fold enhanced rate of covalent modification for nRAB27A-Cys123 over nRAB27A-Cys188, and 10-fold over GSH (Figure 2G). Binding to nRAB27B-C123 was also confirmed by intact mass spectrometry (Figure S1).

### X-ray crystallography reveals novel IMP-1704/RAB27A interactions

In order to gain insight into the binding mode of **1** (**IMP-1704**) we solved the X-ray crystallographic structure of ligand-modified fRAB27A-Cys123 to a resolution of 2.1 Å (Figure 3A, PDB: 8P3G, Table S1), revealing that the compound occupies the WF pocket (Figure 3A, Figure S3). The phenyl ring is oriented towards the hydrophobic patch generated by Met93, displacing the Tyr122 outwards as previously observed for covalent fragments,^13^ alongside forming a key hydrogen bond between one of the sulfonamide oxygens and Lys11 (Figure 3A). We next sought to visualize the non-covalent complex formed prior to covalent modification by soaking **1** into crystals of fRAB27A-Ala123 (fRAB27A[C123A/C188S]) revealing a similar non-covalent interaction with Lys11, which orients the acrylamide group towards RAB27A Cys123 (Figure 3B, Figure S4, PDB: 8P3J, 2.1 Å). Interestingly, an additional non-covalent binding pocket was observed with a second molecule in the vicinity of the WF pocket, but this site is distal to Cys123 and therefore unlikely to be involved in covalent modification (Figure S5). These structures highlighted a viable vector for fragment growth from the pyrrolidine ring with the potential to extend further into the effector binding site and thereby displace effector-RAB27A PPIs.

**Figure 3.**
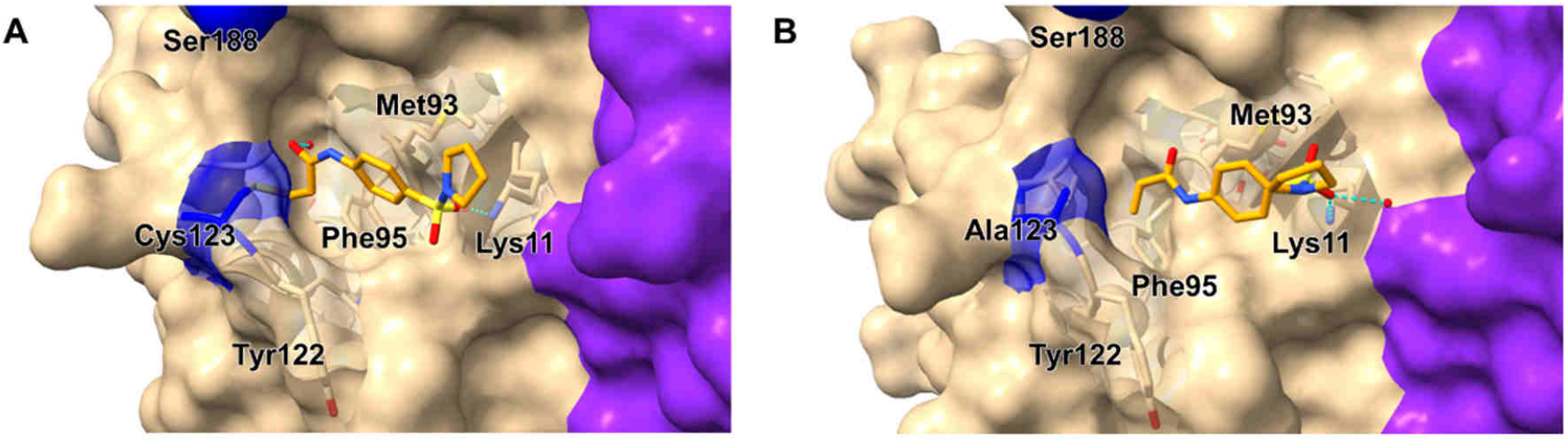
A) Structure of fRAB27A-Cys123 (PDB: 8P3G) covalently bound to hit **1** (orange) showing key interactions with Lys11; B) Non-covalent complex between fRAB27A-Ala123 and hit **1** (PDB: 8P3J), confirming that the affinity-based orientation favors covalent reaction with Cys123.

### Fragment growth identifies ligands with enhanced selective reactivity for RAB27A-Cys123

We next established a fluorescence polarization (FP) assay for rapid analysis of PPI inhibitors at the effector binding site using a novel high-affinity non-covalent cyclic peptide probe for the WF pocket of nRAB27A recently discovered in our labs (**IMP-2660**, Figure S6, *manuscript in preparation*), conjugated to a TAMRA fluorophore. This assay enabled continuous kinetic analysis of **IMP-2660-T** displacement from nRAB27A-Cys123 (*K*_*d*_*= 221 nM*, Figure S6) in 384-well plates, and high-throughput determination of *k*_inact_/*K*_i_.

We used this setup to explore the structure–activity relationships (SAR) for the pharmacophore of hit **1** (Figure 4A,B). Consistent with X-ray crystallography, the *para*-arylsulfonamide moiety was found to be critical for binding, likely contributing to both affinity and reactivity of the series, with analogous amide **2** and *meta*-sulfonamide **3** being inactive (Figure 4B). Functionalization of the pyrrolidine with a 2-(*S*)-methyl ester (**4, IMP-1712**) was well tolerated and led to 2-fold enhancement in modification kinetics, consistent with a conformation leading to enhanced hydrogen bonding interactions with Lys11 and the backbone of Arg90, as observed by X-ray crystallography for this analogue (Figure 4C). Inversion of the stereochemistry of **4** drastically reduced activity both in FP and intact mass spectrometry (MS) assays (**5**, Figure 4B), demonstrating a stereoselective interaction between **4** and RAB27A (Figure 4C). We then proceeded to explore SAR for the corresponding amide analogues of this ester derivative and identified compound **7** with significantly enhanced potency for nRAB27A-Cys123 (>5-fold increased *k*_inact_/*K*_i_ over **1**, Figure 4A,B). Matched enantiomer **8** displayed significantly reduced binding in this assay, consistent with an enantioselective interaction. As an orthogonal method to confirm covalent engagement and Cys123 selectivity, we employed intact protein MS to measure covalent binding against nRAB27A-Cys123 or nRAB27A-Cys188 (Figure 4B). Modification of RAB27A (500 nM) in the presence of excess compound (10 µM) showed clearly favored binding of this series for the Cys123-containing protein, with a single modification event at the expected mass, confirming covalent engagement and Cys123 selectivity (Figure 4B). These results were consistent with the FP data and revealed that selectivity for Cys123 was completely retained over Cys188, demonstrating that the improvement in potency was not driven by intrinsic reactivity of the inhibitors. **Nexinhib-20** (Figure 1C) also exhibited covalent behavior, but in stark contrast displayed no selectivity for Cys123 nor Cys188 (Figure 4B).

**Figure 4.**
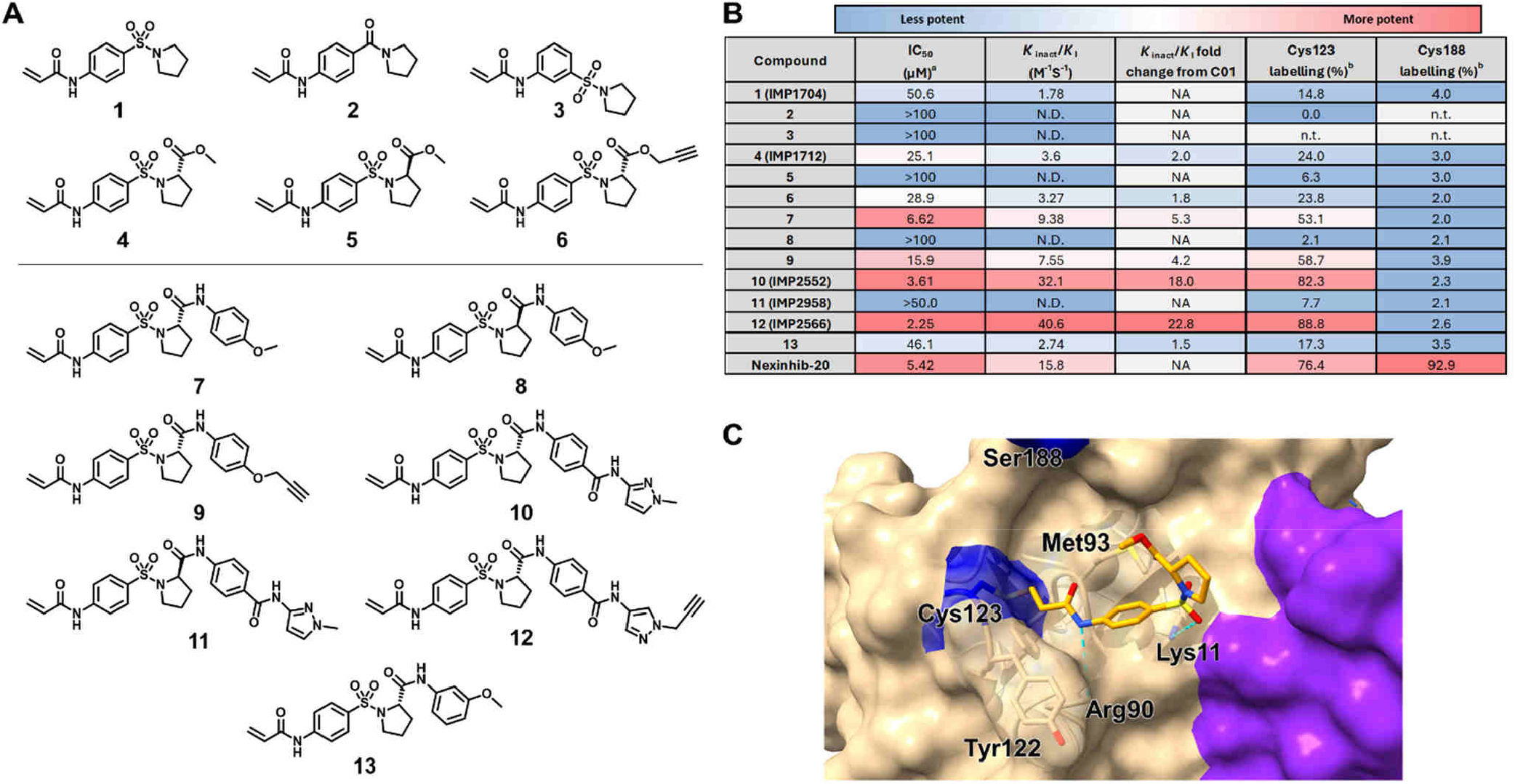
A) Chemical structure of compounds synthesized for pharmacophore analysis of **1** (top) and fragment growth campaign analogues (bottom); B) Activity values determined by fluorescence polarization (FP) and intact protein MS assays. Color scale highlights derivatives from least potent (blue) to most potent (red). a = IC_50_ determined by the FP assay (200 nM nRAB27A-Cys123, 35 nM **IMP-2660-T**) at 2 h; b = percentage labelling determined by intact protein MS following 1 h incubation of protein (500 nM) with compound (10 µM) at rt; N.D.= not determined; n.t. = not tested; C) Structure of fRAB27A-Cys123 (PDB: 8P3H) covalently bound to **4** (orange) showing key interactions with Lys11 and the backbone of Arg90.

### In-cell target engagement of endogenous RAB27A/B by clickable covalent probes

To explore whether improved binding in biochemical assays correlated with engagement of endogenous RAB27A/B in cells, we synthesized corresponding alkyne-functionalized analogues of **4** and **7, 6** and **9** respectively, and confirmed that these small modifications distal to the acrylamide warhead did not significantly affect covalent modification of recombinant nRAB27A-Cys123 (Figure 4B). We evaluated RAB27A expression in different cancer cell lines, and identified triple negative breast cancer cell line MDA-MB-231 as a suitable model system with readily detectable RAB27A expression (Figure S7). Incubation of MDA-MB-231 cells with alkyne-tagged probe followed by lysis, ligation to biotin-azide, and analysis by streptavidin mass-shift assay revealed good target engagement by **9**, with significantly higher modification of endogenous RAB27A compared to the less potent probe **6** (Figure 5A), and in a dose-dependent manner as confirmed by immobilized streptavidin pull-down experiments (Figure 5B). Encouraged by the significant improvement in cellular engagement of RAB27A, we explored the broader *in cellulo* selectivity of alkyne **9** and corresponding inhibitor **7** by pull-down and competitive chemical proteomics. Incubation with **9** (10 µM) for 2 h showed that RAB27A was significantly enriched in probe treated samples, and was found to be within the top 8.1% of engaged proteins (Figure S11). Consistent with our hypothesis, RAB27B was also found to be significantly enriched in samples treated with probe **9**, albeit with lower engagement, likely due to its lower expression in this cellular system (Figure S8). Competition experiments with 2 h pre-incubation with **7** (20 µM) or negative control **8** (20 µM) showed that only the active enantiomer inhibitor **7** competed for engagement of RAB27A/B, while inactive control **8** maintained a similar off-target reactivity profile but did not block RAB27A/B binding by **9** (Figure 5C, Figure S12). These results represent the first example of covalent in-cell target engagement for any Rab protein, providing a benchmark for further optimization of activity and selectivity.

**Figure 5.**
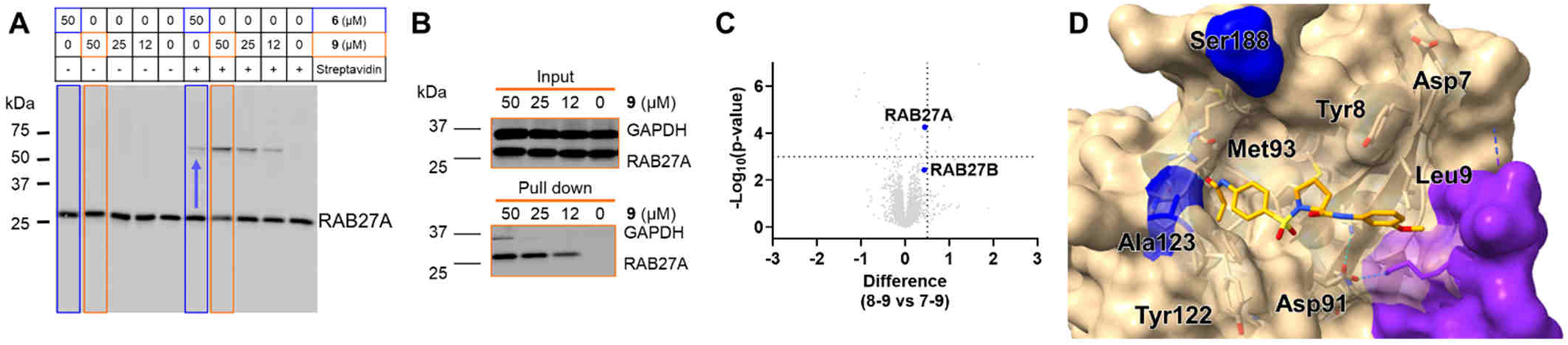
A) Western-blot analysis of streptavidin-shift assay using alkyne-tagged probes **6** and **9** (2 h incubation); B) Western blot analysis (α-GAPDH and α-RAB27A) of dose-dependent pull-down of RAB27A by probe **9** using CuAAC chemistry and streptavidin beads; C) Volcano plot for chemical proteomics competition experiments showing the difference between targets competed by **7** and inactive control **8** against probe **9**.Treatment conditions: **7** (20 µM) or inactive control **8** (20 µM) for 2 h followed by **9** (10 µM) for 2 h; D) Non-covalent complex between fRAB27A-Ala123 (sand) and **13** (orange, PDB: 8P3K), showing orientation of the second phenyl ring toward the fused effector region.

### Structure-guided optimization delivers potent stereoselective RAB27A/B probe IMP-2552 and inactive enantiomer control IMP-2958

With the first cellular RAB27A probes in hand, we set out to improve potency and selectivity by further extending into the effector binding pocket to effectively disrupt RAB27/effector PPIs, based on elucidation of the non-covalent binding mode by X-ray crystallography of **13** (Figure 5D). *Para*-substituted analogue **7**, despite being more potent, did not result in defined crystal structures likely due to clashes with the fused effector. Elaboration through an amide linker yielded compound **10** (**IMP2552**, Figure 4A) which exhibited enhanced binding kinetics for nRAB27A-Cys123 with a *k*_inact_/*K*_i_ of 32.1 M^-1^s^-1^ in the competition FP assay (Figure 4B). Selectivity for Cys123 over Cys188 was further corroborated for **10** by intact protein MS analyses, showing a single labeling event for nRAB27A containing both Cys123 and Cys188 (Figure 6A), and for nRAB27B-Cys123, consistent with the high homology between RAB27A and B in the WF pocket. **IMP-2552** was found to engage both GTP-(active) and GDP-bound (inactive) forms of nRAB27A-Cys123 with equal potency by intact protein MS, suggesting that binding for these compounds at the WF pocket is nucleotide-independent (Figure S9). Consistent with the expectation that these optimized analogues would interfere with effector binding, reactivity with effector fusion construct fRAB27A-Cys123 was significantly reduced relative to nRAB27A-Cys123, and we were unable to determine interactions introduced by the methylated pyrazole ring through X-ray crystallography. Based on the bound trajectory observed for compound **13** (Figure 5D), we hypothesize that additional interactions may be made towards RAB27A, potentially weakening or disrupting existing effector interactions such as between RAB27A Asp91 with SLP2-A Lys14, which support crystal packing in our X-ray models. Importantly, stereoselectivity was maintained in these analogues, with matched enantiomer **11** (**IMP-2958**) showing significantly reduced RAB27A binding in both the FP and intact MS assays (Figure 4B), providing an ideal negative control probe for off-target binding in phenotypic assays. In order to assess *in cellulo* engagement of **10**, a corresponding alkyne-bearing analogue, **12** (**IMP-2566**), was synthesized. It should be noted that the pyrazole regioisomers between **10** and **12** differ slightly, owing to the synthetic challenges of synthesizing the propargylated 1,3-isomer. Probe **12** significantly engaged endogenous RAB27A at low micromolar concentrations in MDA-MB-231 cells, reaching ∼50% engagement after 24 h incubation at 10 µM, as quantified by streptavidin-shift assays (Figure 6B,C) and pull-down experiments (Figure 6D). Evidence from streptavidin-shift analysis of transiently overexpressed FLAG-tagged RAB27A-WT, RAB27A-Cys123[C188S] and RAB27A-Cys188[C123S] mutants further supported specific Cys123 modification in cells (Figure 6E). We observed that the α-FLAG antibody was less sensitive towards the streptavidin-RAB27A complex, but this could be better visualized with α-RAB27A antibody, which also showed engagement of endogenous RAB27A across all treatment conditions with **12** (see Figure S10). Chemical proteomics revealed significantly improved selectivity for RAB27A, observing an improved engagement in the top 1.7% of enriched targets across the proteome (Figure S11). Competition experiments once again confirmed stereoselective inhibition of RAB27A/B engagement by **10** but not by **11** (Figure 6F, Figure S12). No other significantly competed protein belonged to the Rab or Rab effector families, minimizing the risk of off-target impacts on vesicle trafficking. No cytotoxicity was observed at concentrations that delivered significant RAB27A/B target engagement in chemoproteomic analyses for these compounds (Figure S13). Overall, the cellular profile of positive and negative control compounds with good off-target overlap (see supporting tables for proteomics) supports the use of these probes in cell-based systems at acceptable concentrations for the study of RAB27A/B-related phenotypes.

**Figure 6.**
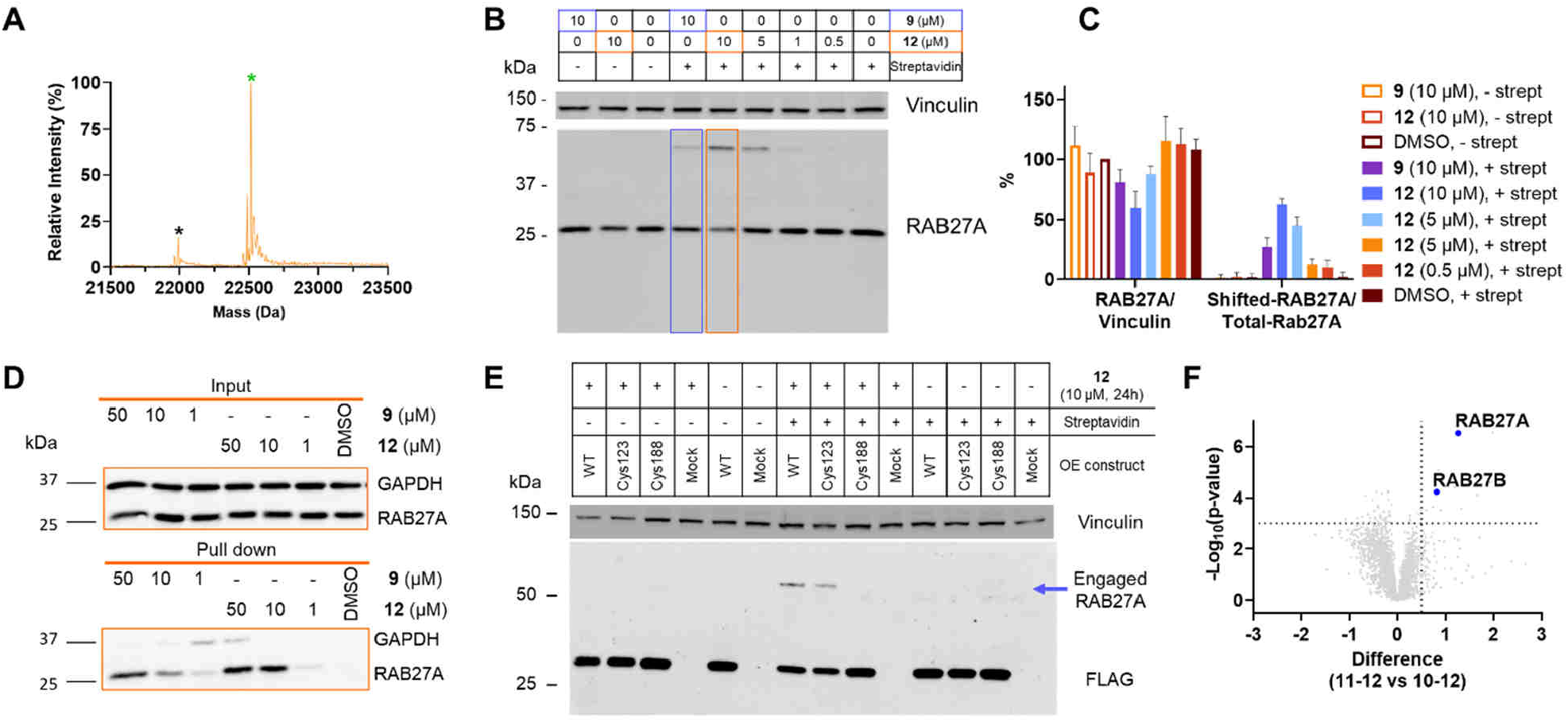
A) Intact protein MS analysis of nRAB27A-Cys123,Cys188 (1–193, black asterisk, MW: 21991 Da) following incubation with **10** (10 µM, 1 h, rt) showing expected single covalent adduct formation (green asterisk, MW: 22514 Da); B) Streptavidin-shift assay comparing alkyne probes **9** and **12** (24 h incubation); C) Quantification of streptavidin-shift assay shown in B (n= 3 independent replicates); D) Western blot analysis (α-GAPDH and α-RAB27A) of dose-dependent pull-down of RAB27A by probe **12** using CuAAC chemistry and streptavidin beads; E) Western blot analysis of streptavidin-shift assay using α-FLAG antibody. Importantly, only mutants containing Cys123 are engaged by **12**, hence the shifted band is only present in WT and Cys123 lanes. WT= FLAG-RAB27A (1–221, P51159); Cys123= FLAG-RAB27A (1–221, C188A); Cys188= FLAG-RAB27A (1–221, C123A); Mock= no DNA control; F) Volcano plot for chemical proteomics competition experiments showing the difference between targets competed by **10** and inactive control **11** against probe **12.**Treatment conditions: **10** (10 µM) or inactive control **11** (10 µM) for 2 h followed by **12** (5 µM) for 2 h.

### Targeting the WF pocket with IMP-2552 disrupts RAB27-effector interactions and phenocopies RAB27A knockout in melan-a melanocytes

In order to test our hypothesis that a targeted covalent WF pocket ligand would inhibit RAB27A/B activity by displacing effectors, we developed a co-immunoprecipitation (Co-IP) assay for transiently expressed GFP-tagged full length hSLP2-A to evaluate interaction with native RAB27A in cells, employing GFP-trap beads for enrichment. Whilst unperturbed PPIs between hSLP2-A and RAB27A resulted in the expected Co-IP of RAB27A, treatment of cells with either lead inhibitor **10** or alkyne probe **12** significantly blocked RAB27A-SLP2-A interaction at 10 and 20 µM, while inactive enantiomer control **11** showed no effect up to 20 µM (Figure 7A,B). Taken together with in-cell target engagement data, these results further support the use of these compounds as the first in-cell targeted covalent RAB27A/B PPI inhibitors.

**Figure 7.**
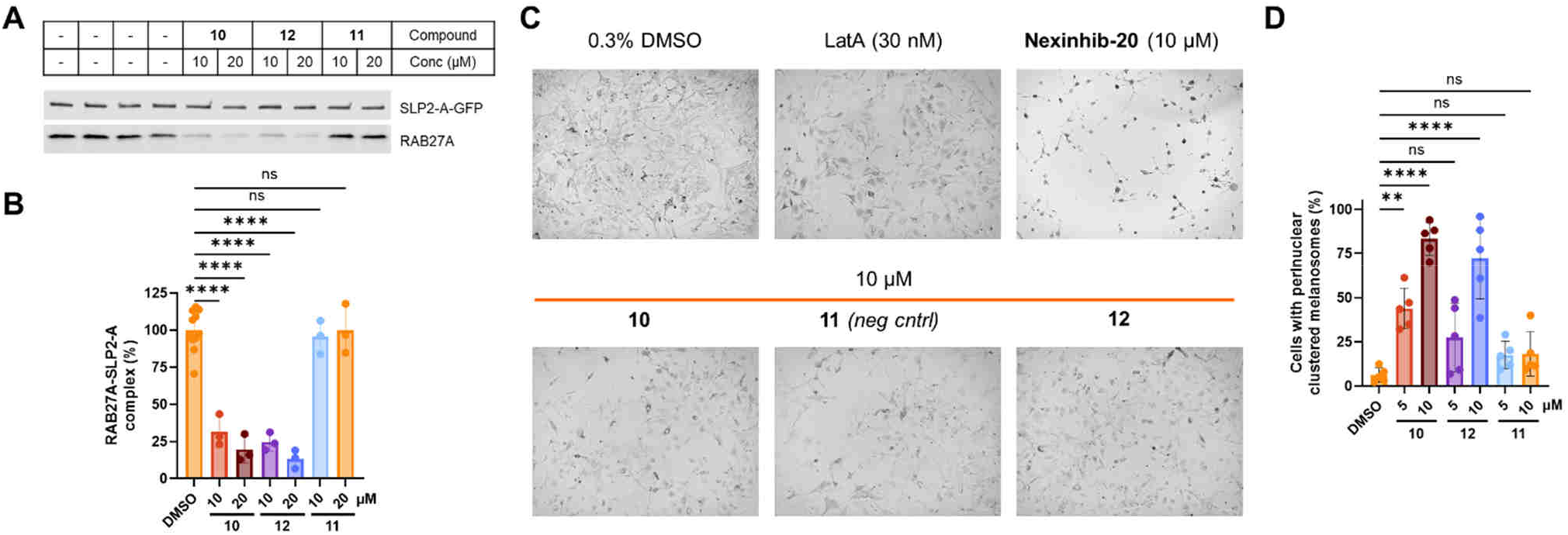
A) Western blot analysis of GFP-trap Co-IP assays of MDA-MB-231 cells transiently expressing GFP-hSLP2-A. Treatment conditions: **10, 11**, and **12** (10 and 20 μM, 14 h); B) Quantification of Co-IP experiments shown in A (n= 3 independent replicates). ****p<0.0001, ns= not significant; C) Representative brightfield images of melan-a mouse melanocytes treated with Latrunculin A (30 nM), **Nexinhib-20** (10 µM), **10** (10 µM), **11** (10 µM) or **12** (10 µM) for 24 h, white scale bar is 100 µm; D) Quantification of melanocytes displaying perinuclear melanosomes from assays shown in A (n= 5 independent replicates, see montage in Figure S14).

The role of RAB27A in the regulation of skin and hair pigmentation is well-established, and loss of function is responsible for the hypopigmentation observed in Griscelli type II patients. RAB27A knockout (KO) in melanocytes or mutations leading to loss of binding to effector protein SLAC2-A (melanophilin) result in perinuclear accumulation of melanosomes, the specialized transport vesicles that shuttle melanin outside the cell to the epidermis, and causes hypopigmentation *in vivo*.^29^ This phenotype is fully dependent on RAB27A activity, and can be readily assayed by brightfield microscopy in melanocytes. We treated melan-a mouse melanocytes at increasing probe concentrations for 24 h and quantified cells exhibiting perinuclear accumulation of melanosomes (Figure 7C,D, Figure S14). In agreement with our previous cellular results, both **10** and **12** dose-dependently inhibited melanosome distribution leading to perinuclear clustering, while enantiomeric inactive compound **11** showed similar distribution of melanosomes to untreated melanocytes (Figure 7C). Consistent with selective RAB27A inhibition, no significant cytotoxicity was observed for compounds **10, 11** or **12** at the tested concentrations over the course of the experiment (Figure S15).

**Nexinhib-20** has been widely used in cellular and *in vivo* studies of RAB27A,^27^ despite a lack of direct evidence for on-target activity. In this assay, **Nexinhib-20** showed profound cytotoxicity and cell death at 10 µM (Figure 7C, Figure S15), a standard concentration used for this compound in cellular assays.^30^ These data, together with evidence for promiscuous reactivity (Figure 4) and the presence of PAIN motifs in the structure, suggest that Nexinhib-related compounds should not be applied in studies of RAB27A/B. Prior studies, which found Nexinhib compounds unable to recapitulate RAB27A KO, may merit revisiting with bona fide RAB27A/B probes.^31^

## Conclusions

Our work demonstrates the first successful application of a targeted covalent approach to tackle a challenging small Rab GTPase, expanding the scope of covalent inhibition in the Ras superfamily.^32^ Optimized probes engage native RAB27A/B at a non-conserved cysteine residue (Cys123) and consequently inhibit protein-protein interactions that drive RAB27A/B cellular activity, producing the first validated chemical probes to study this target, alongside matched enantiomeric negative controls.Starting from a suitable covalent fragment and a carefully selected vector for elaboration, we were able to identify and develop the first small molecule inhibitors of the RAB27A/effector PPI through structure-guided fragment growth towards the effector interface, supported by a suite of biochemical and cellular assays and X-ray crystallography. Early introduction of a stereocenter of defined configuration essential for specific binding to RAB27A/B allowed the development of matched enantiomer inactive probes as negative controls, enabling comparative analysis in complex cellular models by normalizing for potential off-target effects, as shown in melanocyte cells. Chemical proteomics studies confirmed that our lead probes, inhibitor **10** (**IMP-2552**) and alkyne probe **12** (**IMP-2566**), but not matched negative control **11** (**IMP-2598**), engage RAB27A and RAB27B in cells, and enantioselectively disrupt RAB27 PPIs.Our results also invalidate **Nexinhib-20** as a putative selective probe for RAB27A/B, inviting great caution in its use in any biological system, in agreement with several previously published studies.^31^ In summary, our work brings together a novel toolbox of chemical probes to advance the study of RAB27A/B biology, and its future exploration as an actionable target in diseases from cancer to chronic inflammation. We will be pleased to share these probes with interested labs to advance this field. We anticipate that some of the concepts applied here, for example the generation of a potent non-covalent peptide probe for high-throughput analysis of covalent modification leading to inhibition of a PPI, may prove useful for other targets. More broadly, we demonstrate a structure-guided “covalent-first” approach to produce cell-active orthosteric small molecule inhibitors for a specific PPI in the context of a very challenging and to date largely intractable target class.

## Acknowledgements

We thank Diamond Light Source for access to beamline I02 (proposal No. mx17221 and mx23620), which contributed to the results presented here. The X-ray facility at Imperial College was funded by BBSRC (BB/D524840/1) and the Wellcome Trust (202926/Z/16/Z). The authors would like to acknowledge the services available within the Agilent Measurement Suite at Imperial College which enabled intact protein mass spectrometry experiments and HPLC purification. We thank Dr Jianan Lu (Imperial), Mass Spectrometry Technical Specialist for running the proteomics samples discussed in this study. The authors would like to acknowledge the scientific insight provided by Prof Wendy Barclay (Imperial) and Dr Joelle Mettier (Imperial), Dr Pauline Franz for the MDA-MB-231 whole proteome data, as well as Dr Mostafa Jamshidiha for their contribution to the fusion Rab construct design, published in 2022.^13^ The research, MT and EDV were supported by Cancer Research UK (Drug Discovery Committee grant C29637/A20781 and C29637/ A26892 to EC and EWT). The research, RP, EDV and AMT were also supported by Worldwide Cancer Research (Grant 22-0170 to EDV, JCN and EWT). EDV was also supported by a H2020 (EC) MSCA-IF, project RabTarget4Metastasis (EU project 890900). RP was supported by a H2020 (EC) MSCA-IF, project ProCenDecl (EU project 797995). KL was supported by a Wellcome Trust Fellowship Ref: 314556/Z/24/Z. Further support for this project was granted by the Imperial Confidence in Concept (ICiC) scheme (PS3291 and PS3305).

## Notes

A GB Patent application for the compounds disclosed in this work has been filed by the authors (EDV, DB, MT, AMT, SV, EC, and EWT) with Application n° 2416727.2, filed November 13, 2024.

## Supporting Information

X-ray crystallography structure solved for this work have been deposited on the PDB database (PDBs: 8P3G, 8P3H, 8P3J, 8P3K, 8P3l). Proteomics data have been deposited on PRIDE (whole proteome data: PXD077793; chemical proteomics data: PXD077741)

Supplementary figures and additional information about organic synthesis procedures, biochemical assays and biological assays is provided in the Supporting Information file.

## Graphical Abstract

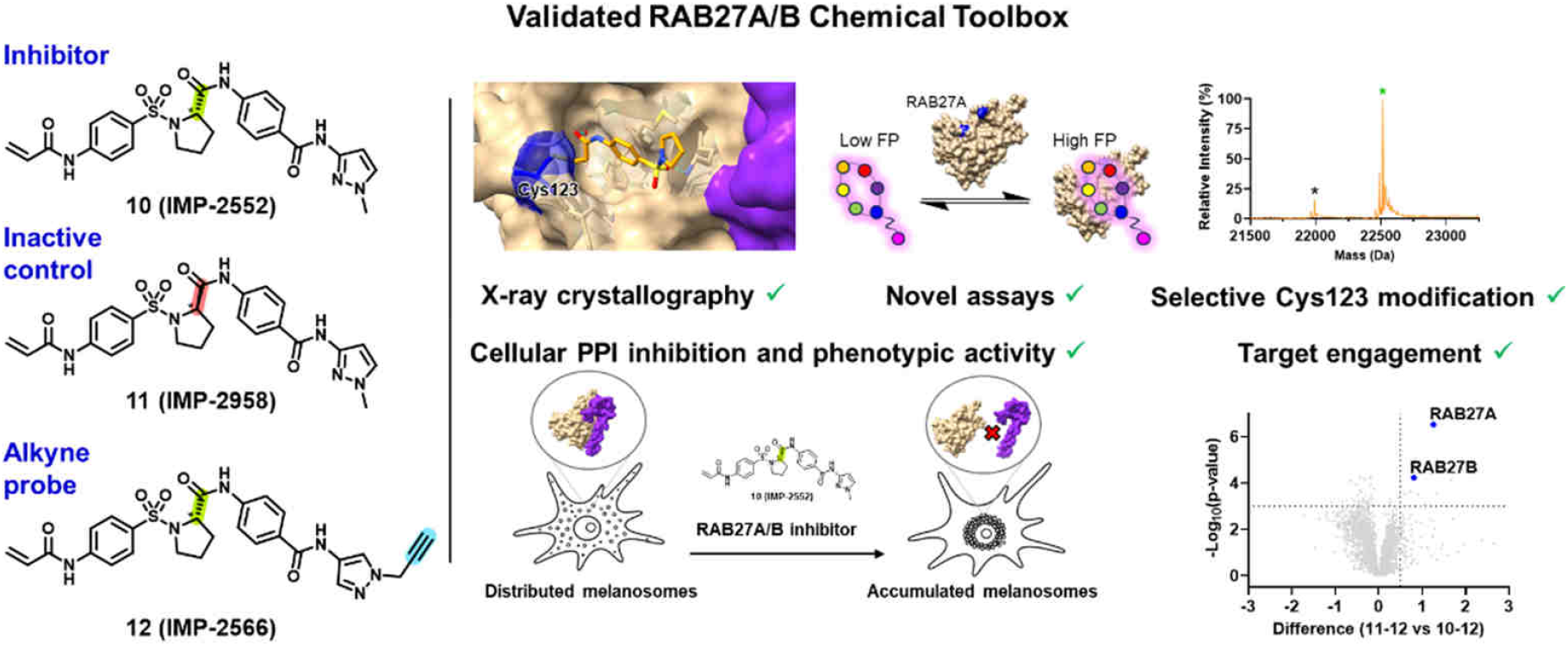

