## Supporting Information File for "Discovery of stereoselective targeted covalent inhibitors of the RAB27-effector protein-protein interaction"

**for**

**Table of contents**


- **Supplementary Figures ........................................................................S3**
- **Supplementary Tables ........................................................................S13**
- **List of abbreviations ........................................................................S17**
- **Biological Procedures ........................................................................S21**
- **Organic Synthesis ........................................................................S29**
- **NMRs ........................................................................S45**
- **HPLC Traces ........................................................................S58**
- **References ........................................................................S71**

### Supplementary Figures

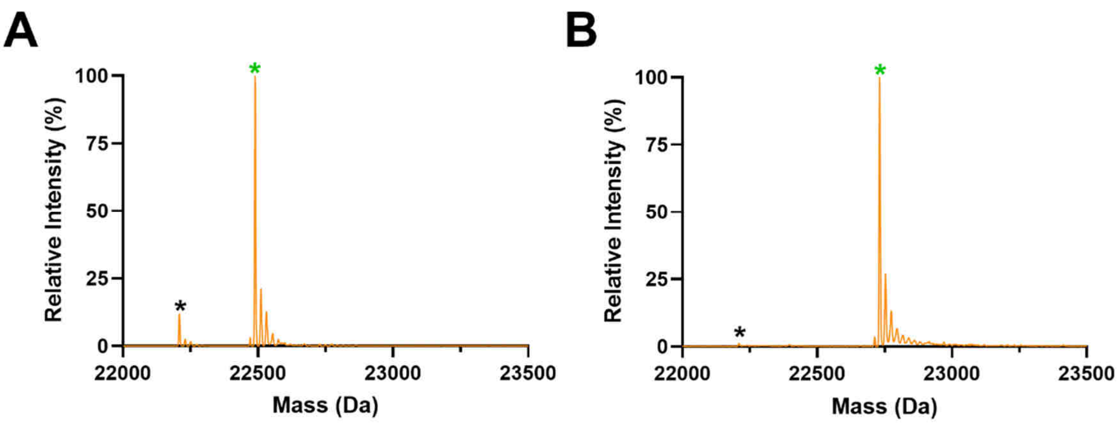


**Figure S1.** Intact protein MS analysis of 500 µM compound **1** (A) and 10 µM compound **10** (B) incubated for 1 h at rt with nRAB27B-Cys123 (black asterisk, MW: 22209 Da) showing expected single covalent adduct formation (green asterisk, A → MW: 22489 Da; B → MW: 22731 Da).

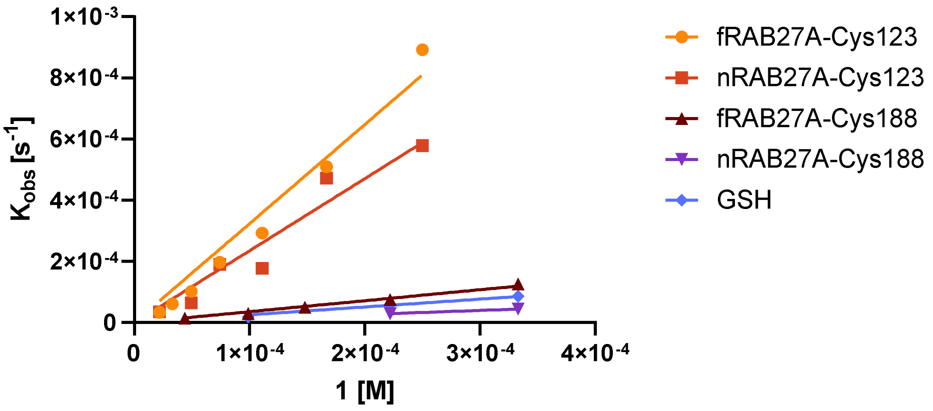


**Figure S2.** Calculated $K_{obs}$ from dose-response qIT assay for compound **1**.

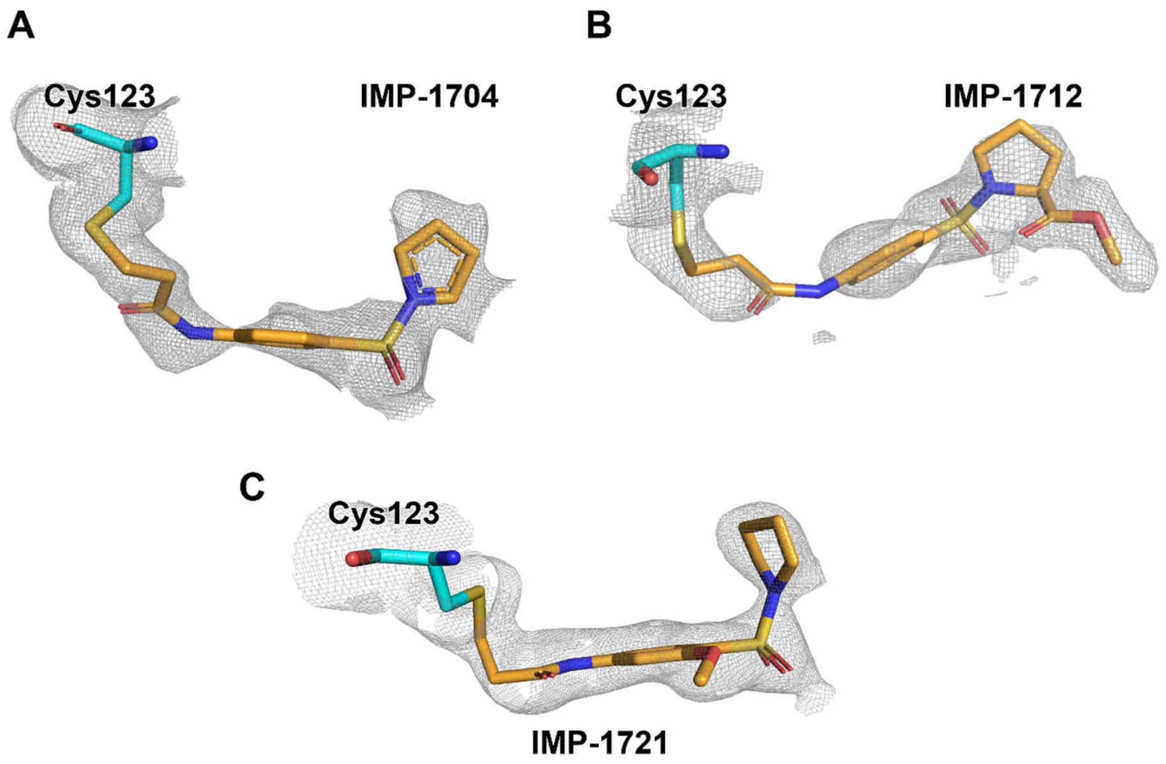


**Figure S3. Electron density maps for covalent complexes of fRAB27A-Cys123 solved by X-ray crystallography.** 2FO-FC electron density maps (grey) contoured at 1.0 σ for covalently bound ligands (orange) **IMP-1704** (**1**, A), **IMP-1712** (**2**, B) and **IMP-1721** (**14**, C).

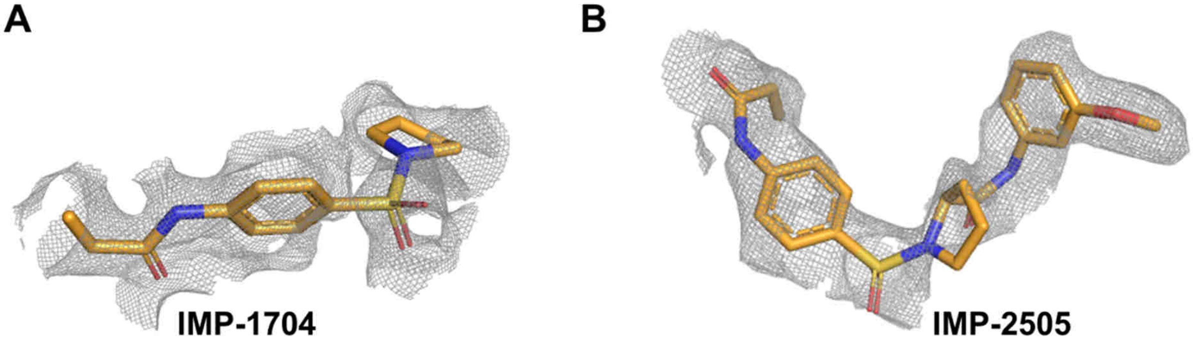


**Figure S4. Electron density maps for non-covalent complexes of fRAB27A-Ala123 solved by X-ray crystallography.** 2FO-FC electron density maps (grey) contoured at 1.0 σ for non-covalently soaked ligands (orange) **IMP-1704** (**1**, A), and **IMP-2505** (**13**, B).

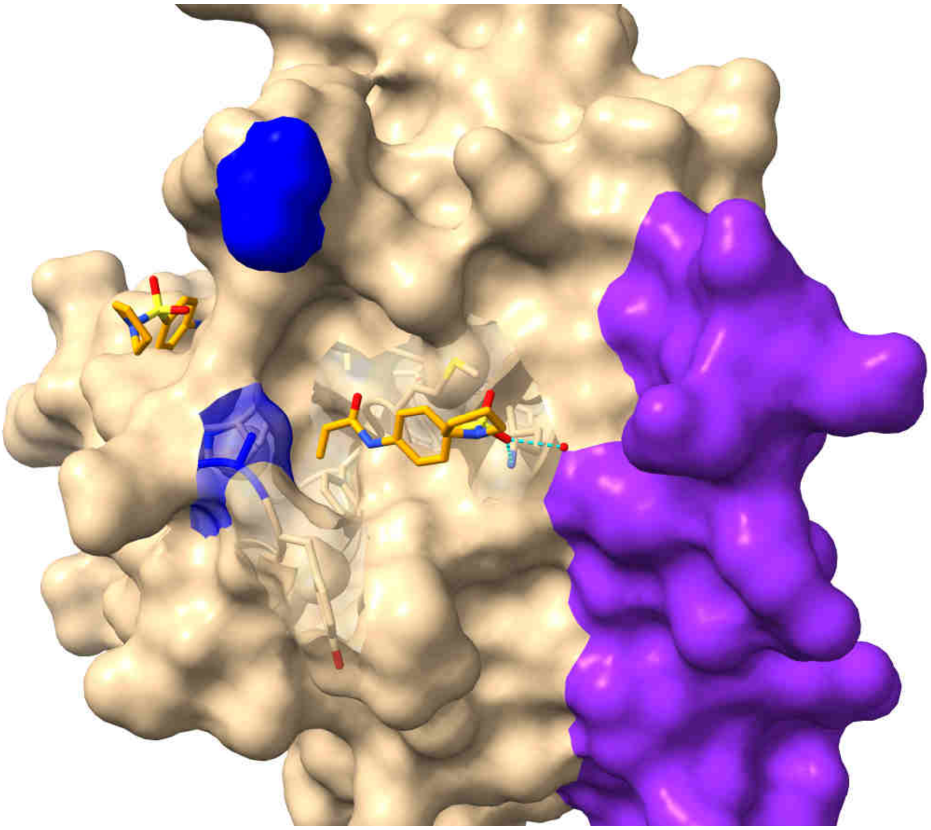

**Figure S5.** Non-covalent complex between fRAB27A-Ala123 and hit **1** (PDB: 8P3J) showing a second ligand-binding pocket.

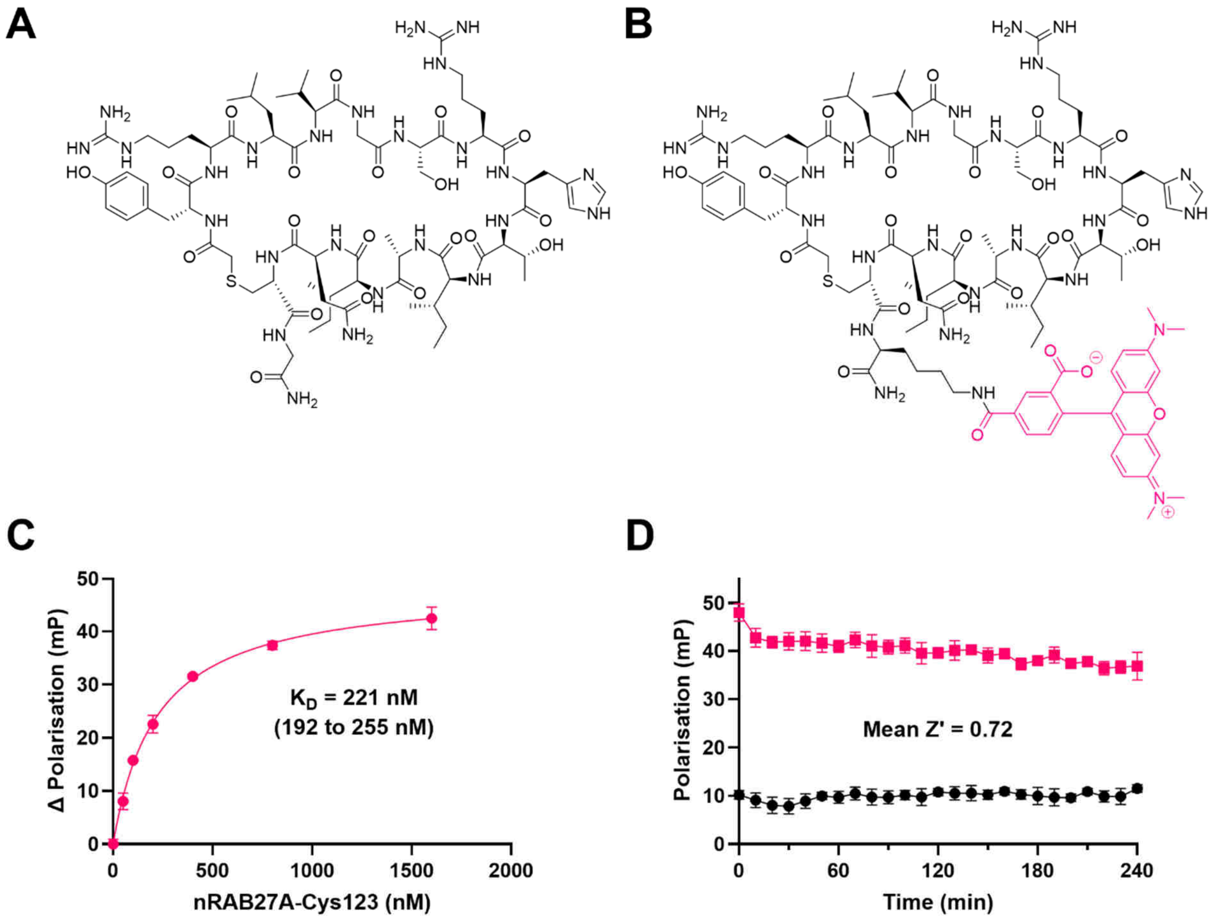


**Figure S6.** A) Chemical structure of **IMP2660**; B) Chemical structure of **IMP-2660-T**; C) $K_D$ measurement for **IMP-2660-T** via fluorescence polarization, including 95% CI; D) Fluorescence polarization assay window under standard conditions (200 nM nRAB27A-Cys123, 35 nM **IMP-2660-T**).

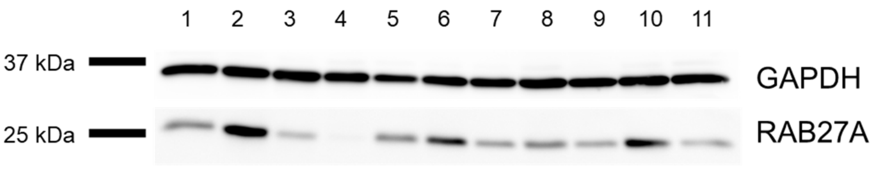


**Figure S7.** Analysis of RAB27A expression levels in a panel of breast and prostate cell lines by western blotting. *Lane 1* = MCF7; Lane 2 = MDA-MDB-231; Lane 3 = SKBR3; Lane 4 = BT474; Lane 5 = T47D; Lane 6 = DU-145; Lane 7 = PC3; Lane 8 = LnCap; Lane 9 = Vcap; Lane 10 = BxPC3; Lane 11 = Capan2.

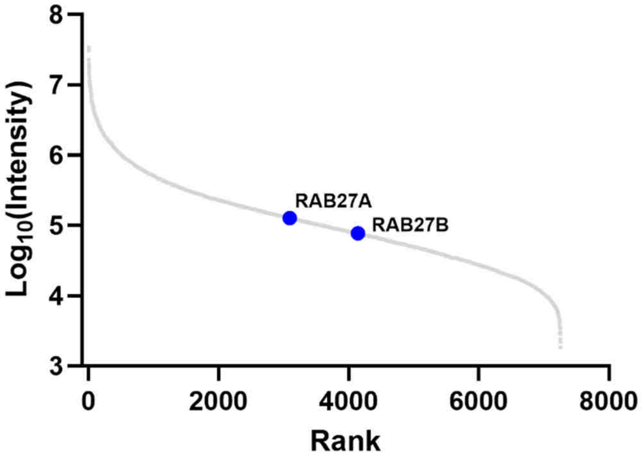


**Figure S8.** Analysis of RAB27A and RAB27B expression levels by whole proteome analysis in MDA-MDB-231 showing moderate expression.

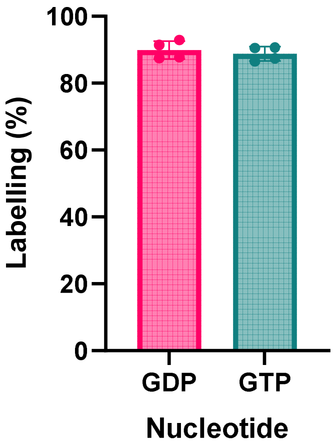


**Figure S9.** Labelling efficiency for compound **10** against GDP- or GTP-bound nRAB27A-Cys123(L78) determined by intact protein MS.

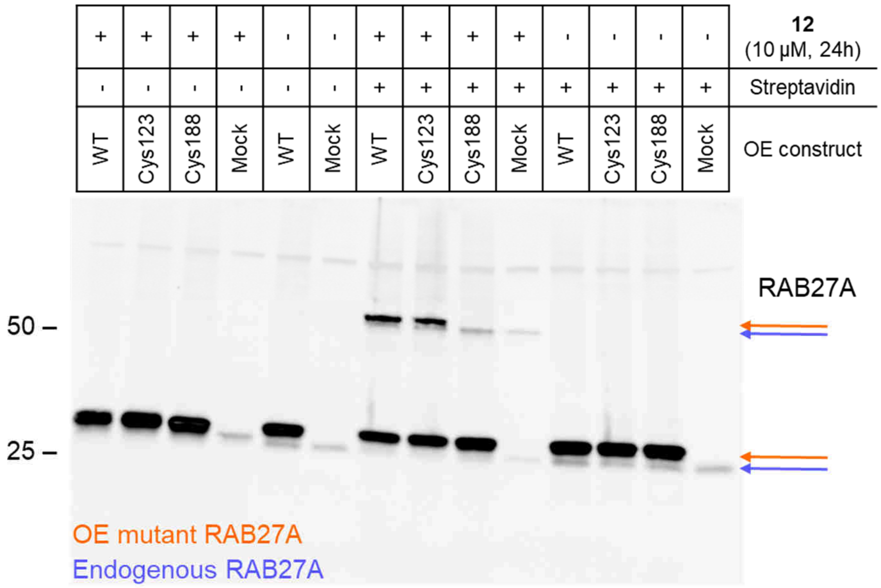


**Figure S10.** Western blot analysis of streptavidin-shift assay using α-RAB27A antibody. Two close bands corresponding to endogenous RAB27A (lower) and the overexpressed (OE) FLAG-tagged mutant specified (upper band) are observed. Only mutants containing Cys123 are engaged by **12**, hence the OE mutant shifted band is only appearing in WT and Cys123 lanes, while engaged endogenous RAB27A is present in all treated samples. This is the same blot as in Figure 6E after stripping. WT= FLAG-RAB27A (1–221, **P51159**); Cys123= FLAG-RAB27A (1–221, **C188A**); Cys188= FLAG-RAB27A (1–221, **C123A**); Mock= no DNA control.

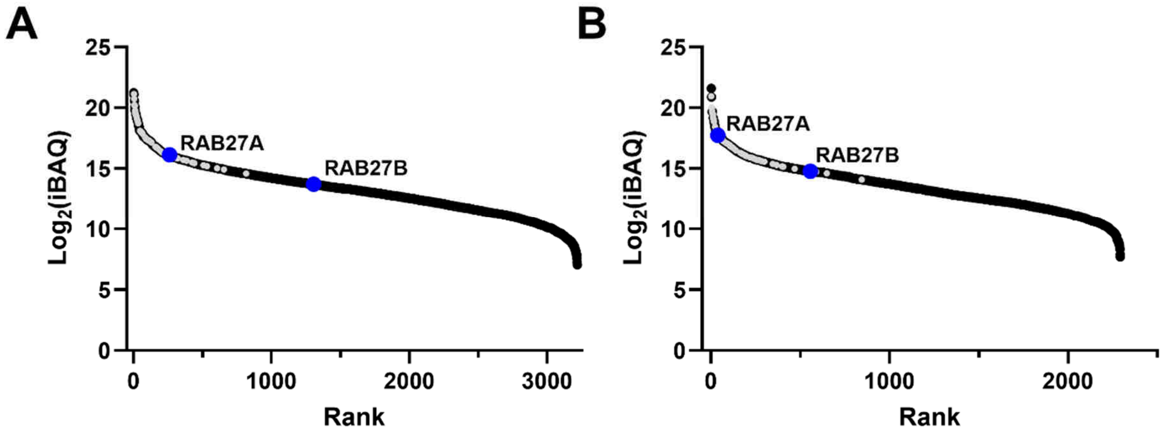


**Figure S11.** iBAQ-normalized S-plots showing the ranked enrichment of protein targets engaged by probes **9** (A) and **12** (B) in chemical proteomics experiments. Proteins shown in grey were also detected in DMSO-treated controls, likely reflecting non-specific, sticky, or highly abundant proteins rather than true targets.

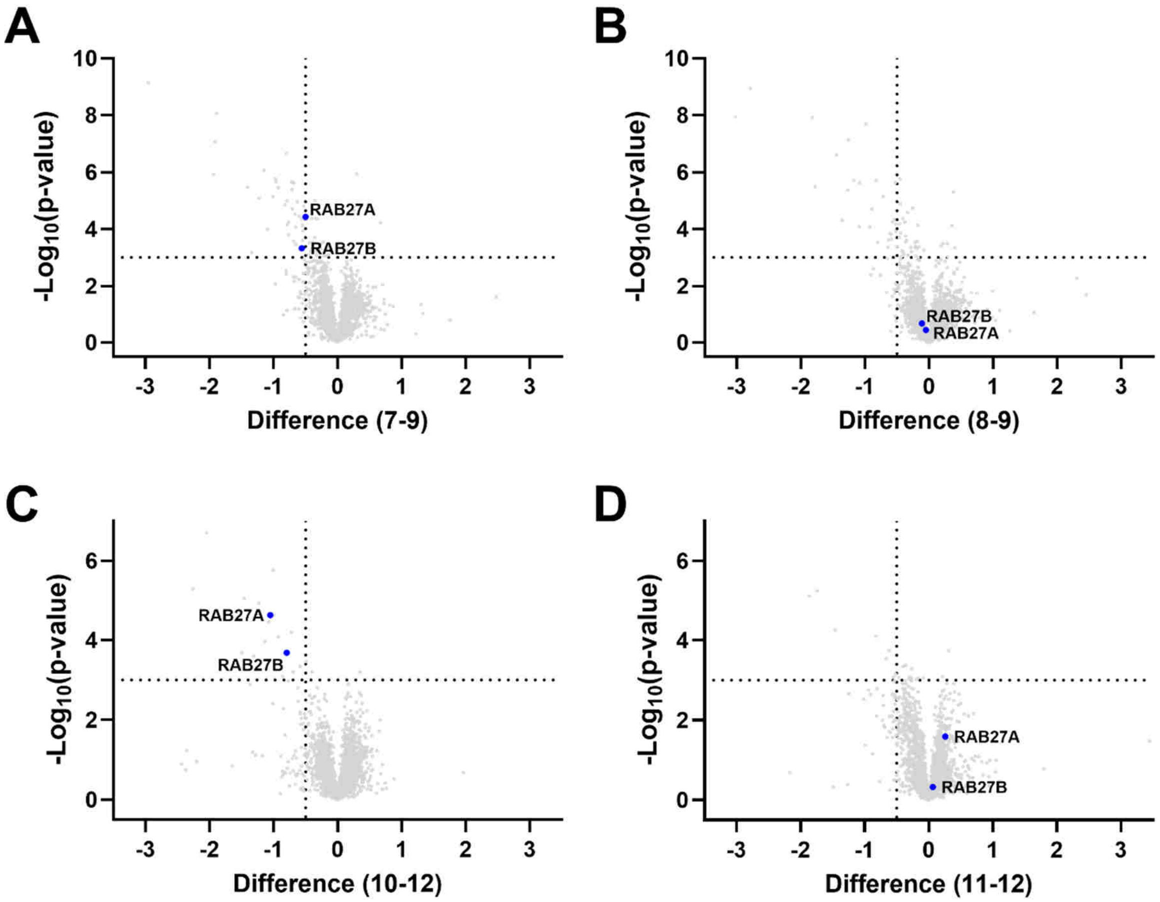


**Figure S12.** Volcano plots for chemical proteomics competition experiments showing the difference between targets competed by 20 µM **7** (A) or inactive control **8** (B) against 10 µM probe **9**, and targets competed by 10 µM **10** (C) or inactive control **11** (D) against 5 µM probe **12**. Treatment conditions: Parent or inactive control incubated for 2 h followed by probe incubation for 2 h.

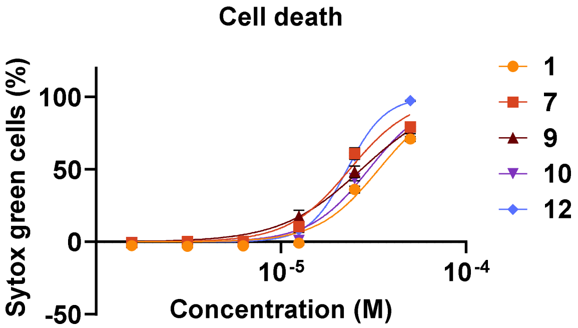


**Figure S13.** Cell death analysis following 72 h treatment in MDA-MB-231 cells with compounds obtained via Incucyte® live-cell imaging using sytox green to count death cells.

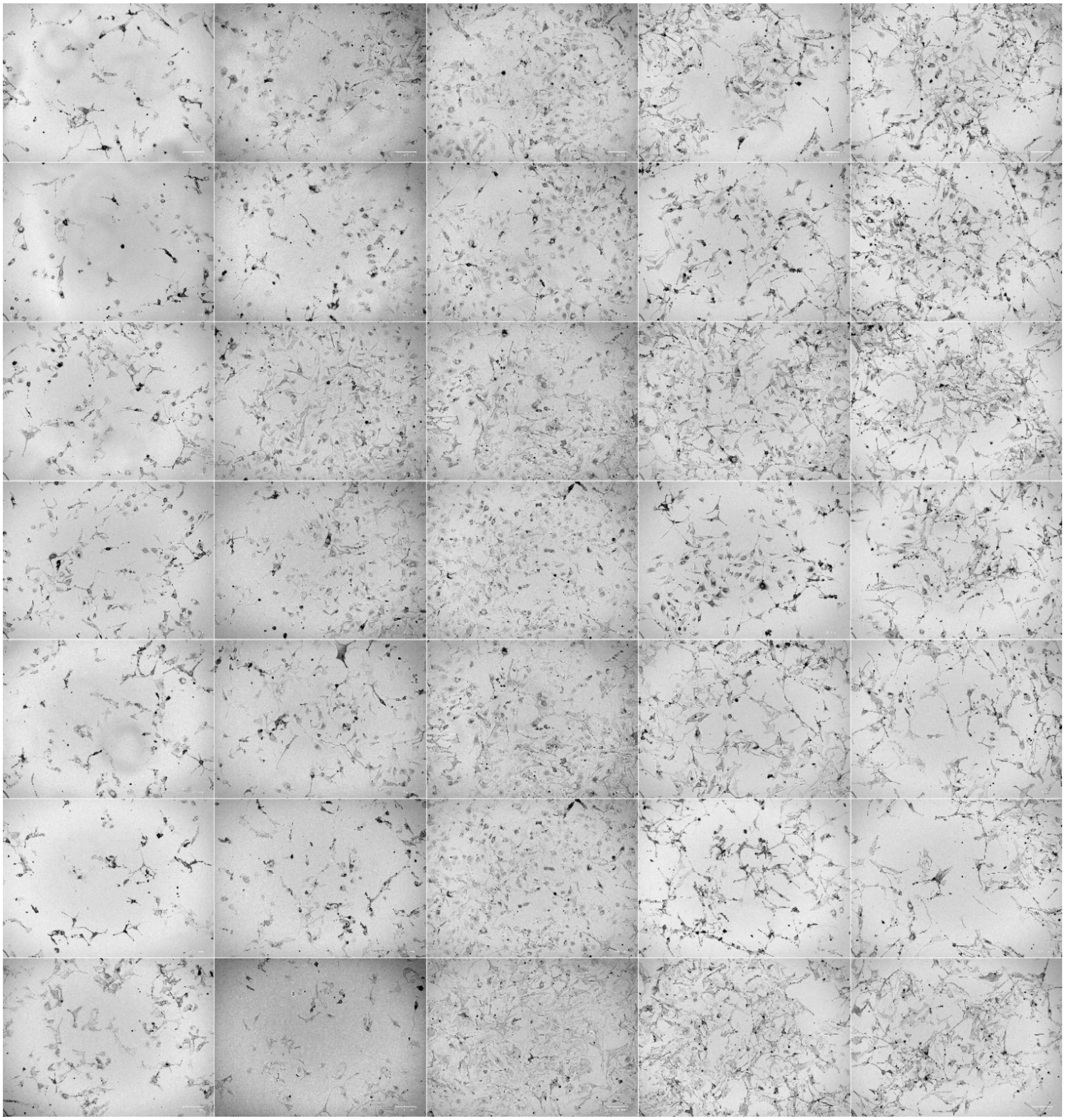

**Figure S14.** Montage of mouse melanocyte images used for quantification analyses shown in Figure 7B. Conditions array is displayed in the table below for the 5 biological replicates:

| | | | | | |
|---|---|---|---|---|---|
| **Cmpd 10, 5 µM** | **R #1** | **R #2** | **R #3** | **R #4** | **R #5** |
| **Cmpd 10, 10 µM** | **R #1** | **R #2** | **R #3** | **R #4** | **R #5** |
| **Cmpd 12, 5 µM** | **R #1** | **R #2** | **R #3** | **R #4** | **R #5** |
| **Cmpd 12, 10 µM** | **R #1** | **R #2** | **R #3** | **R #4** | **R #5** |
| **Cmpd 11, 5 µM** | **R #1** | **R #2** | **R #3** | **R #4** | **R #5** |
| **Cmpd 11, 10 µM** | **R #1** | **R #2** | **R #3** | **R #4** | **R #5** |
| **DMSO** | **R #1** | **R #2** | **R #3** | **R #4** | **R #5** |

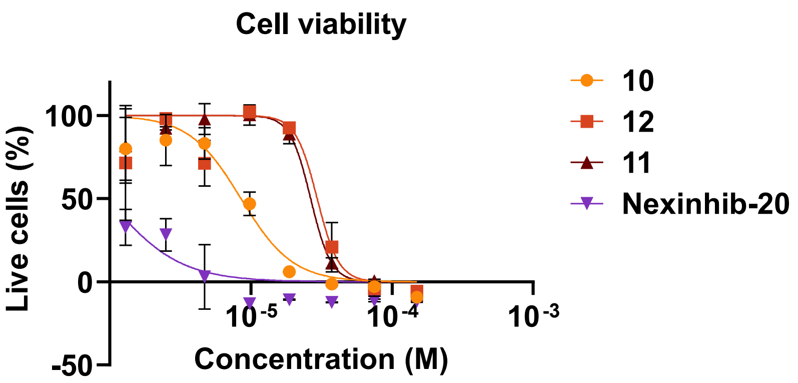


**Figure S15.** Melan-a cell viability following 24 h treatment with specified compound, as determined by Prestoblue assay (ThermoFisher A13262) according to the manufacturer protocol.

### Supplementary Tables

**Table S1.** Data Processing and Refinement Statistics for fusion hSlp2-a_RAB27A-C123 bound to **IMP1704** (**1**) (PDB: 8P3G).

| | **8P3G** |
|---|---|
| **Data collection** | |
| Wavelength (Å) | 0.9795 |
| Space group | $P2_12_12_1$ |
| Unit cell (Å, °) | a = 62.30, b = 76.03, c = 118.85; α = β = γ = 90 |
| Resolution (Å) | 55.18–2.13 (2.21–2.13) |
| Total reflections | 64,540 |
| Unique reflections | 32,270 (3,162) |
| Multiplicity | 2.0 (2.0) |
| Completeness (%) | 99.72 (99.06) |
| ⟨I/σ(I)⟩ | 4.2 |
| CC1/2 | 0.981 (0.344) |
| Wilson B (Å$^2$) | 26.37 |
| **Refinement** | |
| Reflections (work/test) | 32,207 / 1,614 |
| Rwork / Rfree | 0.193 / 0.243 (0.261 / 0.314) |
| No. atoms | 3,833 |
| Protein residues | 418 |
| RMS bonds (Å) | 0.007 |
| RMS angles (°) | 0.86 |
| Ramachandran favored (%) | 97.26 |
| Ramachandran allowed (%) | 2.74 |
| Ramachandran outliers (%) | 0.00 |
| Rotamer outliers (%) | 0.28 |
| Clashscore | 3.75 |
| **Average B-factors (Å$^2$)** | |
| Macromolecules | 31.91 |
| Ligands | 33.98 |
| Solvent | 35.90 |
| Overall | 32.36 |

*Values in parentheses correspond to the highest-resolution shell.*

**Table S2.** Data Processing and Refinement Statistics for fusion hSlp2-a_RAB27A-C123 bound to **IMP1712** (**2**) (PDB: 8P3H)

| | **8P3H** |
|---|---|
| **Data collection** | |
| Wavelength (Å) | 0.9795 (Diamond I04) |
| Space group | $P2_12_12_1$ |
| Unit cell (Å, °) | a = 60.1723, b = 77.6959, c = 116.15; α = β = γ = 90 |
| Resolution (Å) | 58.08 – 2.76 (2.859 – 2.76) |
| Total reflections | 14,556 |
| Unique reflections | 14,556 (1,419) |
| Multiplicity | 2.0 (2.0) |
| Completeness (%) | 99.68 (98.95) |
| ⟨I/σ(I)⟩ | 5.21 |
| CC1/2 | 0.972 (0.227) |
| Wilson B (Å$^2$) | 56.54 |
| **Refinement** | |
| Reflections (work/test) | 14,514 / 723 (1,408 / 71) |
| Rwork / Rfree | 0.227 / 0.313 (0.321 / 0.353) |
| Non-H atoms | 3,459 |
| — Macromolecules | 3,264 |
| — Ligands | 148 |
| — Solvent | 47 |
| Protein residues | 409 |
| RMSD bonds (Å) | 0.009 |
| RMSD angles (°) | 1.06 |
| Ramachandran favored (%) | 92.07 |
| Ramachandran allowed (%) | 7.16 |
| Ramachandran outliers (%) | 0.77 |
| Rotamer outliers (%) | 0.00 |
| Clashscore | 17.79 |
| **Average B-factors (Å$^2$)** | |
| Macromolecules | 64.90 |
| Ligands | 62.39 |
| Solvent | 58.24 |
| Overall | 64.70 |

*Values in parentheses correspond to the highest-resolution shell.*

**Table S3.** Data Processing and Refinement Statistics for fusion hSlp2-a_RAB27A-C123 bound to **IMP1721** (**14**) (PDB: 8P3I)

| | **8P3I** |
|---|---|
| **Data collection** | |
| Wavelength (Å) | 0.9763 (Diamond I03) |
| Space group | $P2_12_12_1$ |
| Unit cell (Å, °) | a = 61.3673, b = 76.6009, c = 118.674; α = β = γ = 90 |
| Resolution (Å) | 64.36 – 2.00 (2.071 – 2.00) |
| Total reflections | ~73,200* (calculated from redundancy × unique) |
| Unique reflections | 38,565 (3,801) |
| Multiplicity | 1.9 (1.9) |
| Completeness (%) | 87.98 (21.07) |
| ⟨I/σ(I)⟩ | 3.39 |
| CC1/2 | 0.997 (0.331) |
| Wilson B (Å$^2$) | 47.15 |
| **Refinement** | |
| Reflections (work/test) | 33,930 / 1,757 (801 / 48) |
| Rwork / Rfree | 0.2325 / 0.2796 (0.4565 / 0.5171) |
| Non-H atoms | 3,586 |
| — Macromolecules | 3,337 |
| — Ligands | 192 |
| — Solvent | 57 |
| Protein residues | 426 |
| RMSD bonds (Å) | 0.008 |
| RMSD angles (°) | 1.05 |
| Ramachandran favored (%) | 96.36 |
| Ramachandran allowed (%) | 3.64 |
| Ramachandran outliers (%) | 0.00 |
| Rotamer outliers (%) | 0.29 |
| Clashscore | 7.09 |
| **Average B-factors (Å$^2$)** | |
| Macromolecules | 58.03 |
| Ligands | 57.36 |
| Solvent | 55.10 |
| Overall | 57.95 |

*Values in parentheses correspond to the highest-resolution shell.*

**Table S4.** Data Processing and Refinement Statistics for fusion hSlp2-a_RAB27A-A123 bound to **IMP1704** (**1**) (PDB: 8P3I)

| | **8P3J** |
|---|---|
| **Data collection** | |
| Wavelength (Å) | 0.9795 (Diamond I04) |
| Space group | $P2_12_12_1$ |
| Unit cell (Å, °) | a = 62.4364, b = 75.8281, c = 118.53; α = β = γ = 90 |
| Resolution (Å) | 63.88 – 2.16 (2.237 – 2.16) |
| Total reflections | 30,872 |
| Unique reflections | 30,871 (3,056) |
| Multiplicity | 1.9 (1.9) |
| Completeness (%) | 96.39 (76.33) |
| ⟨I/σ(I)⟩ | 2.8 |
| CC1/2 (overall) | 0.972 (0.325) |
| Wilson B (Å$^2$) | 15.21 |
| **Refinement** | |
| Reflections (work/test) | 29,767 / 1,522 (2,335 / 120) |
| Rwork / Rfree | 0.2847 / 0.3508 (0.6497 / 0.7102) |
| Non-H atoms | 3,686 |
| — Macromolecules | 3,429 |
| — Ligands | 160 |
| — Solvent | 97 |
| Protein residues | 426 |
| RMSD bonds (Å) | 0.008 |
| RMSD angles (°) | 1.02 |
| Ramachandran favored (%) | 95.45 |
| Ramachandran allowed (%) | 3.59 |
| Ramachandran outliers (%) | 0.96 |
| Rotamer outliers (%) | 0.00 |
| Clashscore | 10.10 |
| **Average B-factors (Å$^2$)** | |
| Macromolecules | 28.63 |
| Ligands | 34.28 |
| Solvent | 23.07 |
| Overall | 28.73 |

*Values in parentheses correspond to the highest-resolution shell.*

**Table S5.** Data Processing and Refinement Statistics for fusion hSlp2-a_RAB27A-A123 bound to **IMP2505** (**13**) (PDB: 8P3K)

| | **8P3K** |
|---|---|
| **Data collection** | |
| Wavelength (Å) | 0.9795 (Diamond I04) |
| Space group | $P2_12_12_1$ |
| Unit cell (Å, °) | a = 60.8957, b = 76.5361, c = 117.795; α = β = γ = 90 |
| Resolution (Å) | 47.65 – 2.58 (2.672 – 2.58) |
| Total reflections | 17,938 |
| Unique reflections | 17,938 (1,736) |
| Multiplicity | 6.6 (6.6) |
| Completeness (%) | 98.75 (94.77) |
| ⟨I/σ(I)⟩ | 4.17 |
| CC1/2 (overall) | 0.959 (0.216) |
| Wilson B (Å²) | 42.54 |
| **Refinement** | |
| Reflections (work/test) | 17,722 / 859 (1,650 / 74) |
| Rwork / Rfree | 0.2410 / 0.3143 (0.3642 / 0.4219) |
| Non-H atoms | 3,565 |
| — Macromolecules | 3,340 |
| — Ligands | 186 |
| — Solvent | 39 |
| Protein residues | 421 |
| RMSD bonds (Å) | 0.009 |
| RMSD angles (°) | 1.06 |
| Ramachandran favored (%) | 91.04 |
| Ramachandran allowed (%) | 8.23 |
| Ramachandran outliers (%) | 0.73 |
| Rotamer outliers (%) | 0.00 |
| Clashscore | 17.51 |
| **Average B-factors (Å²)** | |
| Macromolecules | 46.97 |
| Ligands | 50.13 |
| Solvent | 37.91 |
| Overall | 47.03 |

*Values in parentheses correspond to the highest-resolution shell.*

**Abbreviations**

AzB: azidobiotin

BME: β-mercaptoethanol

DIPEA: diisopropylethylenediamine

DMEM: Dulbecco's modified Eagle medium

DMF: dimethylformamide

DMSO: dimethylsulfoxide

DTT: dithiothreol

EDCI: 1-ethyl-3-(3-dimethylaminopropyl)carbodiimide

FP: fluorescence polarization

GAPDH: glyceraldehyde 3-phosphate dehydrogenase

GDP: guanosine diphosphate

GTP: guanosine triphosphate

GppNHp: guanosine 5′-(β,γ-imino)triphosphate

HATU: hexafluorophosphate Azabenzotriazole Tetramethyl Uronium

HBTU: hexafluorophosphate Benzotriazole Tetramethyl Uronium

HEPES: 4-(2-hydroxyethyl)-1-piperazineethanesulfonic acid

HOBt: hydroxybenzotriazole

HRMS: high resolution mass spectrometry

IPTG: isopropyl β-D-1-thiogalactopyranoside

LB: lysogeny broth

LC/Q-TOF: liquid chromatography/quadrupole time-of-flight

LRMS: low resolution mass spectrometry

NTA: nitrilotriacetic acid

MES: 2-(N-morpholino)ethanesulfonic acid

PBS: phosphate-buffered saline

PCR: polymerase chain reaction

qIT: quantitative irreversible tethering

Rab: Ras-related protein in brain

SDS-PAGE: sodium dodecyl sulfate-polyacrylamide gel electrophoresis

TAMRA: carboxytetramethylrhodamine

TCEP: tris(2-carboxyethyl)phosphine

TEV: Tobacco Etch Virus

TIC: total ion chromatograms

THF: tetrahydrofuran

Tris: tris(hydroxymethyl)aminomethane

### Biological Procedures

#### Protein expression and purification

As previously reported,[1] all RAB27A constructs contain the sequence for human RAB27A (P51159, residues 1–192, mutations: Q78L and/or C123S/A or C188S or both as specified), which was cloned into a pET15b vector (Invitrogen) including a N-terminal His-tag followed by a Tobacco Etch Virus (TEV) recognition site (ENLYFQ¦G). Fusion constructs also contain the C-terminus of Slp2a SHD1 (SFLTEEEQEAIMKVLQRDAALKRAEEER (residues 5–32 in the fusion)) linked to the N-terminus of RAB27A via a flexible poly glycine-serine linker (GSGSGSG). nRAB27B-C123 constructs contain the sequence for human RAB27B (O00194, residues 1–193, mutation: C188A), cloned in the same vector.

For protein expression, plasmids were transformed to E. coli BL21 cells and spread on LB agar plates containing 100 mg/L Ampicillin for selection. Single colonies were picked for amplification and incubated overnight into LB media containing 100 mg/L Ampicillin at 37 °C, shaking. Big scale cultures were inoculated using these overnight cultures at 1% v/v and grown at 37 °C until absorbance at 600 nm reached 0.6–0.8. Protein expression was induced using 0.5–1 mM isopropyl β-D-1-thiogalactopyranoside (IPTG) for 3 hours at 37 °C. Subsequently cells were pelleted at 5,000 ×g for 15 min, then re-suspended in lysis buffer containing 500 mM NaCl, 10 mM imidazole, 5 mM $MgCl_2$ and 50 mM Tris at pH 8. Cells were lysed with a cell disruptor at 25K psi and centrifuged at 17,000 ×g for 45 min. The supernatant was loaded on $Ni^{2+}$-NTA resin equilibrated with lysis buffer, washed extensively and eluted using buffer containing 500 mM NaCl, 300 mM imidazole, 5 mM $MgCl_2$ and 50 mM Tris at pH 8. The protein was buffer exchanged using 2x 5 mL Hitrap columns (Cytiva) in tandem with 100 mM NaCl, 5 mM $MgCl_2$ and 50 mM Tris, pH 8. Afterwards TEV protease (obtained in-house as previously described[2]) was added to the protein solution at a molar ratio of 1/20 in the presence of 1 mM DTT, and the solution was incubated overnight at 4 °C, shaking. The solution was clarified by centrifugation (10 min at 10,000 ×g at 4 °C) and buffer exchanged using 2x 5 mL Hitrap columns (Cytiva) in tandem with 100 mM NaCl, 5 mM MgCl2 and 50 mM Tris, pH 8. The solution was then loaded on $Ni^{2+}$-NTA resin and the flowthrough was collected, concentrated to 5.5 mg/mL in 150 mM NaCl, 5 mM $MgCl_2$, 20 mM Tris pH 8 buffer. If GppNHp loading was required, a 10X buffer containing 10 mM $ZnCl_2$ and 2 M $(NH_4)_2SO_4$, 4 molar excess of GppNHp and 25 units of Antarctic phosphatase (New England Biolabs) were added to the solution and incubated overnight at 4 °C. Finally, the sample was loaded on a Superdex S-75 gel filtration column at a flow rate of 1 mL/min. The column was pre-equilibrated with 150 mM NaCl, 5 mM $MgCl_2$, and 20 mM Tris at pH 8 for crystallography or 20 mM HEPES at pH 8 for biochemical assays. The peaks corresponding to RAB27A constructs were analysed by SDS-PAGE, pooled, concentrated and flash frozen using liquid nitrogen.

#### Protein labelling and purification

To a 15 mL falcon tube were added 600 µL of desired construct (100 µM stock), 100 µL of 50% w/v TCEP-agarose beads (ThermoFisher), and 1.8 mL of 100 mM NaCl, 5 mM $MgCl_2$, and 20 mM HEPES pH 8.0. 50 µL of ligand (50 mM stock) were diluted with 2.45 mL of 100 mM NaCl, 5 mM $MgCl_2$, and 20 mM HEPES pH 8.0, followed by centrifugation (2,400 rpm, 5 min). The supernatant was added to the protein mixture and incubated at 4 °C. The labelled protein solution was concentrated to 0.5 mL by using a Vivaspin 20 filter (5,000 MWCO). The protein was diluted with 4.5 mL of 100 mM NaCl, 5 mM $MgCl_2$, and 20 mM HEPES pH 8.0 and concentrated again to 0.5 mL (5x), to remove excess compound, then purified by superdex S-75 gel filtration at a flow rate of 1 mL/min in 150 mM NaCl, 5 mM MgCl2, and 20 mM Tris at pH 8 for crystallography.

### Protein crystallisation

*Covalent complexes:* Pure samples of labelled protein were concentrated between 4–12 mg/mL, in buffer containing 150 mM NaCl, 5 mM $MgCl_2$, and 20 mM Tris at pH 8.0. Crystals were grown at 4 °C using the sitting-drop vapor-diffusion method with a mother liquor containing 150 mM $NH_4SO_4$, 100 mM MES, and 15% PEG 4K, pH 6.0. Crystals were harvested in 30% glycerol.

*Noncovalent soaking (fRAB27A, C123A mutant):* Pre-formed fRAB27A-A123 crystals using conditions above were incubated for 2 h at 4 °C with 1 µL of a solution of the covalent ligand at 6 mM in 12% DMSO.

### Crystal diffraction, data collection and data processing

Data collections were carried out at Diamond Light Source in Oxford using the i03 or i04 beamline at 100K, using Pilatus detector. Data were collected at 0.4°-0.5° oscillations per image with a total oscillation up to 200°. Data processing was carried out using Dials. Initial phases were calculated using the molecular replacement program phaser. The coordinates of RAB27A from chain A of the RAB27a-Slp2a complex (PDB:3BC1) without the nucleotide and the magnesium ion were used as the search model. Subsequently, the initial model generated by phaser was refined through an iterative cycle using COOT and REFMAC5. Final model structures were validated using the Molprobity server at http://molprobity.biochem.duke.edu. All structure images were prepared using Pymol. X-ray data collection, processing and refinement statistics are given in Table S1–S5.

### qIT assay for screening and hit validation

960 electrophilic acrylamides were screened in the qIT assay adapted from Craven et al[3]. Briefly, the reaction buffer (20 mM HEPES pH 8.0, 100 mM NaCl, 5 mM $MgCl_2$) and quench buffer (20 mM HEPES pH 7.4, 100 mM NaCl, 5 mM $MgCl_2$) were prepared, filtered, de-gassed, and re-gassed with Ar for 15 min on ice. Reaction setup: To each well of a 96-well PCR plate (reaction plate), 8 µL of 50% w/v TCEP-agarose beads in reaction buffer was added, followed by the addition of 92 µL of 10.87 µM protein or glutathione (GSH). In a separate 96-well PCR plate (ligand plate), 3 µL of DMSO or 50 mM ligand in DMSO was added to 147 µL reaction buffer and centrifuged (1k rpm, 5 min, 4 °C). 100 µL of ligand solution or DMSO control from the ligand plate was added to the reaction plate (final concentration: 5 µM protein/GSH and 500 µM ligand). After mixing, the TCEP-agarose beads were pelleted by centrifugation (1k rpm, 5 min, 4 °C) and the plate was kept at 4 °C. At a series of time points (t = 0.25. 1, 2, 4, 7, 24, 48, 72, and 96 h), a 3 µL aliquot in duplicate from the reaction plate was quenched in a black 384-well plate, in which each well was pre-filled with 27 µL of 7-Diethylamino-3-(4'-Maleimidylphenyl)-4-Methylcoumarin (CPM) solution (1.4 µM in quench buffer). The fluorescence plate was spun down (1k rpm, 1 min) and incubated for 60 min at room temperature and then fluorescence intensity (excitation/emission: 384/470 nm) was measured on an EnVision™ plate reader. Data analysis: All analyses were conducted using Prism software (Graphpad). Each fluorescence readout was normalized to the average of the DMSO controls. The normalized fluorescence was plotted against time. A one phase exponential decay was fitted to each plot (constraints: $Y(0) > 0.8$; $0 < plateau < 0.3$; $k > 0$).

### Hit selection

Following the screen, we analyzed the obtained 3,840 kinetic curves to estimate $k_{obs}$ values and used these to determine rate enhancement factors (REFs) by dividing the $k_{obs}$ of each construct against the $k_{obs}$ for GSH. To further filter for any thiol intrinsic reactivity, REF values for each construct were normalized against the REF value obtained from the geometric mean of $k_{obs}$ values for the specific construct. Compounds with a normalized REF>5 were considered to have significantly enhanced labelling rates for the specific construct. We further filtered the hitlist by excluding compounds that had showed REF>5 for other targets screened in previous campaigns. Overall, 127 acrylamides with enhanced labelling for a specific cysteine mutant RAB27A construct were selected for a validation screening in which we repeated the qIT assay against nRAB27A-Cys123 and nRAB27A-Cys188 at 500 µM (Figure 2A). Four compounds were selected for validation in dose-response analyses (500 µM, 250 µM, 125 µM, 62.5µM, 0 µM) against all thiols (fRAB27A-Cys123, fRAB27A-Cys188, nRAB27A-Cys123, nRAB27A-Cys188, and GSH). Due to best synthetic tractability, we selected hit compound **IMP-1704**.

### Intact protein MS analyses

Assays were performed in Agilent InfinityLab 384-well plates where 15 µL of freshly prepared and filtered assay buffer (20 mM HEPES, 100 mM NaCl, 5 mM $MgCl_2$, pH 8.0) with 1 µM protein was mixed with 15 µL of the desired compound (20 µM) in filtered assay buffer + 4% DMSO, to give final assay concentrations of 500 nM protein, 10 µM compound, and 2% DMSO. The reaction plate was shaken (~300 rpm) at rt for 60 min, after which the reaction was quenched by the addition of 7.5 µL of 2% formic acid in MilliQ water. Samples were then submitted to LC-MS analysis on an AdvanceBio 6545XT LC/Q-TOF system (Agilent) fitted with an ACQUITY UPLC Protein BEH C4 Column, 300 Å, 1.7 µm, 2.1 mm x 50 mm (Waters) heated to 45 °C. Data was analysed and processed using BioConfirm (Agilent). The total ion chromatograms (TIC) were extracted (region containing protein) and the summed scans were deconvoluted (using MaxEnt). Protein labelling (%) was calculated by dividing the labelled protein intensity by the sum of the unlabelled and labelled protein intensities. Figures were made using GraphPad Prism 10.0.

**Table 1.** Intact protein LC-MS method.

| Time (min) | A – $H_2O$+0.1% FA (%) | B – MeCN+0.1% FA (%) | Flow (mL/min) |
|---|---|---|---|
| 0.00 | 80 | 20 | 0.6 |
| 0.20 | 80 | 20 | 0.6 |
| 1.40 | 35 | 65 | 0.6 |
| 1.70 | 5 | 95 | 0.6 |
| 1.90 | 5 | 95 | 0.6 |
| 2.10 | 80 | 20 | 0.6 |
| 2.30 | 80 | 20 | 0.6 |

### Fluorescence polarization assay

The assay buffer (20 mM HEPES, 100 mM NaCl, 5 mM $MgCl_2$, pH 8.0) was freshly prepared and filtered (0.2 µm), after which 0.005% Tween was added. 2× Peptide-only buffer was prepared by adding 70 nM A10-TAMRA to the assay buffer. 2× Protein and peptide buffer was prepared by the addition of 70 nM A10-TAMRA and 400 nM nRAB27a-Cys123 to the assay buffer.

Compound plate setup: 2× Compound dilutions (seven-fold 1:3 dilution series in triplicate) in filtered assay buffer +4% DMSO were prepared in a 384-well microplate (Corning CLS4514).
Reaction plate setup: 5 µL of 2× peptide-only buffer or 2× protein and peptide buffer were added to a clear-bottom 384-well microplate (Greiner 788096—black, small volume, lobase, MED binding, µCLEAR). The reaction was started by the addition of 5 µL of 2× compound or 4% DMSO in assay buffer from the compound plate to the reaction plate using an Apricot S2 liquid handler. Final assay concentrations: 35 nM A10-TAMRA, 200 nM nRAB27a-Cys123, desired compound concentrations (seven-fold 1:3 dilution series in triplicate), and 2% DMSO. Polarisation (mP) was measured every 10 min for 2 h using a Clariostar plate reader (filter set 1-TAMRA FP, ex. 540-20, em. 590-20, 20 flashes per well).
Data analysis was performed using Graphpad Prism software (Version 10.2.3). Raw mP values were normalised by subtracting the mean mP values of the peptide-only wells, followed by normalisation to the mean mP values of the DMSO-containing peptide and protein wells. $IC_{50}$ values were calculated for 30 and 120 mins by plotting normalised data against concentration and fitting to the “[Inhibitor] vs. normalised response -- Variable slope” model using Prism software. The pseudo-first order rate constants ($k_{obs}$) values were calculated by plotting normalised data against time and fitting to the “one phase exponential association” model (Y=0, Plateau >50) using Prism software. The $k_{obs}$ values with $R^2$ values > 0.6 were then plotted against concentration and fitted to a linear regression model (Y=0) to generate $k_{inact}/K_I$ values.

#### Cell culture

MDA-MB-231 cells were cultivated in Dulbecco's Modified Eagle Medium (DMEM) [high glucose, pyruvate] supplemented with 10% (v/v) fetal bovine serum and grown at 37 °C under a humidified atmosphere supplied with 5% CO2. For all experiments, cells cultivated for ≤ 20 passages were used.

#### Cell death assay in MDA-MB-231

MDA-MB-231 cells were plated in a 96-well plate (Greiner; clear) at $5*10^3$ cells/well in 100 µL of 10% FBS in DMEM. After 16 h, 100 µL of medium containing a 2x final concentration of treatment were added to each well. Conditions tested were 6x 1:2 dilutions from 50 µM in biological triplicate (final DMSO 0.1%). Sytox green was added to medium to a final concentration of 250 nM. 2 µg/mL of puromycin were used to define 100% cell death. Quantification of green fluorescence area (488) over brightfield area was performed using the Incucyte® software. Data were blank corrected, normalized and plotted using Graphpad Prism.

#### SDS–PAGE, WB and antibodies.

12% SDS–PAGE gels were freshly prepared prior to use. Gels were run for 15–20 min at 80 V, then for 120 min at 120 V (running buffer: 35 mM Tris, 285 mM glycine, 0.6% (w/v) SDS in Milli-Qwater). The SDS-gel was blotted on a nitrocellulose membrane for 60 min at 100 V (blotting buffer: 25 mM Tris, 192 mM glycine, 20% methanol) using a Bio-Rad Mini Trans-Blot® Cell or Trans-Blot SD semidry transfer cell setup. The membrane was blocked in 5% milk powder with 0.1% Tween-20 TBS (50 mM Tris, 150 mM NaCl, pH 7.4), washed once with 0.1% Tween-20

TBS, and incubated with the specified antibody overnight at 4 °C. The blot was washed 4 × 5 min with 0.1% Tween-20 TBS, incubated with secondary HRP antibody (rabbit or mouse, Advansta) for 1 h at rt, and washed 4 × 5 min with 0.1% Tween-20 TBS prior to HRP substrate addition and chemiluminescence detection.

| Primary antibody | Target | Dilution | Buffer |
|---|---|---|---|
| RAB27A (D7Z9Q) Rabbit mAb #69295 (Cell Signalling) | RAB27A | 1:1,000 | 3% BSA in 1% Tween-20 PBS |
| Anti-FLAG® (H7425) Rabbit (Aldrich) | FLAG-tag | 1:500 | 1% Tween-20 PBS |
| Anti-Vinculin antibody [EPR8185] (Abcam) | Vinculin | 1:10,000 | 5% milk in 1% Tween-20 PBS |
| Anti-GAPDH antibody [ab9485] (abcam) | GAPDH | 1:5,000 | 3% BSA in 1% Tween-20 PBS |
| Anti-GFP antibody [ab1218] (abcam) | GFP-tag | 1:3,000 | 3% BSA in 1% Tween-20 PBS |

**Click chemistry protocol**

To every 100 µL of the protein sample (0.5–2 mg/mL) was added 6 µL of a mixture containing 1 mM $CuSO_4$, 1 mM tris(2-carboxymethylphosphine) (TCEP), 100 µM tris(1-benzyl-1H-1,2,3-triazol-4-yl)methylamine (TBTA), and 100 µM azide (containing TAMRA (AzT), biotin (AzB) or both (AzTB)), and then the mixture was incubated for 1 h at rt with shaking. The reaction was stopped by protein precipitation with 4 volumes of MeCN. The samples were centrifugated (5 min at 16,000 g) and the supernatant discarded. The obtained pellet was washed with 80% EtOH (3–5x) and sonication, followed by centrifugation (5 min at 16,000 g) and careful removal of the supernatant. Finally, the pellet was dried in a fume hood for 5 min and redissolved stepwise in 0.2% SDS in PBS (starting from 2% SDS in PBS) for further assays (e.g., pull-downs) or directly in 20% β-mercaptoethanol in Laemmli buffer, denatured for 5 min at 95 °C, and subjected to SDS−PAGE analysis.

**Streptavidin shift assay**

MDA-MB-231 cells were plated in a 12-well plate at 5 × $10^5$ cells/well in 1 mL of 10% FBS in DMEM. After 24 h, the cells were treated with the specified compound concentration or DMSO for the specified time at 37 °C. After incubation, the medium was removed and the cells were washed with 2 × 1 mL PBS, lysed with 100 µL ice-cold lysis buffer (1% NP-40 in PBS, containing protease inhibitors [cOmplete™ Protease Inhibitor Cocktail] and benzonase), clarified by centrifugation and protein concentration adjusted to 0.8–1 mg/mL. Cell lysates (20–50 µg protein) were clicked according to the Click Chemistry protocol with AzTB or AzB including precipitation and washing. The dried samples were resuspended in 18–46 µL of 10% BME in Laemmli buffer 1x in PBS. Streptavidin stock solution (100 µM, Jackson Immunoresearch, 016-000-113) was thawed on ice. Each sample was added 2–4 µL Streptavidin (+ Streptavidin samples) or $H_2O$ (– Streptavidin samples) and incubated 10 min at rt before running SDS-PAGE and western blotting according to standard protocol.

### Pull-down experiments

MDA-MB-231 cells were plated in a 12-well plate at 5 × $10^5$ cells/well in 1 mL of 10% FBS in DMEM. After 24 h, the cells were treated with the specified compound concentration or DMSO for 2 h at 37 °C. After incubation, the medium was removed and the cells were washed with 2 × 1 mL PBS, lysed with 100 µL ice-cold lysis buffer (1% NP-40 in PBS, containing protease inhibitors [cOmplete™ Protease Inhibitor Cocktail] and benzonase), clarified by centrifugation and protein concentration adjusted to 0.8–1 mg/mL. Cell lysates (50–150 µg protein) were clicked according to the Click Chemistry protocol with AzB including precipitation and washing. The clicked insoluble samples were redissolved in 0.2% SDS in PBS and dincubated with magnetic streptavidin beads (NEB) in 1:3–10 ratios (beads to protein) for 2 h at room temperature. The beads were washed 5 × 0.2% SDS in PBS, then redissolved in 20% β-mercaptoethanol in Laemmli buffer and incubated for 10 min at 95 °C. The supernatant was collected and subjected to SDS–PAGE analysis followed by western blotting for RAB27A.

### PPI inhibition

MDA-MB-231 cells were seeded in 6-well plates (6 × $10^5$ cells/well) using standard cell culture conditions (2 mL of medium per well). After 24 h each well was transfected with 1.5 µg of purified GFP-Slp2-a plasmid (Addgene #40047) using Lipofectamine 3000 (Invitrogen) in opti-MEM™ medium, according to the manufacturer's instructions. After 6 h, the cells were treated with the specified compound concentration or 0.5% DMSO for 16 h at 37 °C. After incubation, the medium was removed and the cells were washed with 2 × 1 mL ice-cold PBS, lysed with 200 µL ice-cold lysis buffer (20 mM Tris-HCl, 50 mM NaCl, 1 mM EGTA, 1% CHAPS, 1 mM GppNHp, pH 7.5, containing protease inhibitors [cOmplete™ Protease Inhibitor Cocktail]), clarified by centrifugation and protein concentration adjusted to 1 mg/mL.
In lo-bind Eppendorf tubes, magnetic GFP-trap beads (10 µL / well) were washed with 3 × 1 mL ice-cold wash buffer (20 mM Tris-HCl, 50 mM NaCl, 1 mM EGTA, 1% CHAPS, pH 7.5) before the addition of 150 µL of normalized lysate. The samples were rotated for 1 h at 4 °C. Flowthrough was removed, and the magnetic GFP-trap beads were washed with 2 × 1 mL ice-cold wash buffer (20 mM Tris-HCl, 50 mM NaCl, 1 mM EGTA, 1% CHAPS, pH 7.5). Samples were eluted from the beads by the addition of 1× Laemmli buffer with 5% β-mercaptoethanol, and heating at 95 °C for 10 min, followed by centrifugation and collection of the supernatant. Samples were then subject to SDS–PAGE and WB analysis.

### Overexpression experiments

MDA-MB-231 cells were seeded in a 12-well plate (5 × $10^5$ cells/well) using standard cell culture conditions (1 mL of medium per well) and each well was transfected by addition of 50 µL of a preincubated solution containing 1 µg of the specified plasmid (Figure 6E and S10) and 3 µL of Fugene® HD in opti-MEM™ medium. After 18 h, the cells were treated with the specified probe concentration or DMSO for 24 h at 37 °C. Cells were lysed with 1% NP-40 in PBS containing protease inhibitors (cOmplete™ Protease Inhibitor Cocktail), clarified by centrifugation and protein concentration adjusted to 0.8–1 mg/mL. Samples were processed according to the specified protocol.

### Whole proteome proteomics

MDA-MB-231 cells were seeded in a 12-well plate ($3 \times 10^5$ cells/well) using standard cell culture conditions (1 mL of medium per well). After 24 h, the medium was removed and the cells were washed with 2 × 0.5 mL ice-cold PBS, lysed with 100 µL ice-cold lysis buffer (1% NP-40 in PBS containing protease inhibitors [cOmplete™ Protease Inhibitor Cocktail]), clarified by centrifugation and protein concentration adjusted to 1 mg/mL.

Reduction and alkylation of the proteins was carried out by treating with 10 mM tris(2-carboxyethyl)phosphine (TCEP, Sigma-Aldrich) and 40 mM chloroacetamide (CAA, Sigma-Aldrich) for 10 min at rt. Protein was then precipitated by the addition of 4 volumes of acetonitrile (VWR, HPLC grade). The sample was centrifuged, and the supernatant discarded. The pellet was then washed with 3 × 80% ethanol (VWR, HPLC grade). The protein pellet was then treated with trypsin (6.25 ng/µL, Pierce) in a 25 mM ammonium bicarbonate (SigmaAldrich) pH 8.0 buffer and agitated at 37 °C overnight. The samples were dried using a SpeedVac concentrator before being rehydrated in 2% acetonitrile and 0.5% TFA in water (Optima, LC-MS grade).

### Chemical proteomics

MDA-MB-231 cells were seeded in a 12-well plate ($3 \times 10^5$ cells/well) using standard cell culture conditions (1 mL of medium per well). After 24 h, the cells were treated with the specified compound concentration or 0.5% DMSO for the specified incubation time at 37 °C. After incubation, the medium was removed and the cells were washed with 2 × 0.5 mL ice-cold PBS, lysed with 100 µL ice-cold lysis buffer (1% NP-40 in PBS containing protease inhibitors [cOmplete™ Protease Inhibitor Cocktail]), clarified by centrifugation and protein concentration adjusted to 0.8 mg/mL.

In a 96-deep well plate (KingFisher), 100 µL/well of normalised lysate was added, along with 6 µL/well of a mixture containing 1 mM $CuSO_4$, 1 mM tris(2-carboxymethylphosphine) (TCEP), 100 µM tris(1-benzyl-1H-1,2,3-triazol-4-yl)methylamine (TBTA), and 100 µM azide-PEG$_3$-biotin (AzB). The mixture incubated for 1 h at rt with shaking, before being quenched by the addition of 1 µL EDTA (500 mM) to a final concentration of 5 mM.

To each well, 30 µL of a freshly prepared 1:1 mixture of Cytiva magnetic E3 and E7 SpeedBeads were added, followed by the addition of 591 µL of EtOH and incubation at rt for 15 min with shaking. The plate was then transferred to a Thermo KingFisher system where beads were washed twice with 90% EtOH (500 µL) before the protein being eluted in 0.2% SDS in 50 mM HEPES, pH 8.0 (150 µL). Pierce Streptavidin magnetic beads (10 µL/well) were added and incubated for 1 h at rt with shaking. The plate was then transferred to a Thermo KingFisher system where beads were washed with RIPA buffer (500 µL), and 1% SDS in 50 mM HEPES pH 8.0 (500 µL). Proteins were then reduced and alkylated using 10 mM TCEP and 40 mM chloroacetamide in 0.2% SDS and 50 mM HEPES pH 8.0 (500 µL), after which the beads were washed twice with 50 mM HEPES pH 8.0 (500 µL), and eluted in 50 mM HEPES pH 8.0 (200 µL). To each well, 0.2 µg of trypsin (Pierce) was added and samples were agitated at 37 °C overnight. Samples were then filtered, acidified to below pH 3 using formic acid, and frozen at -20 °C.

### LC-MS/MS analysis

Peptides for proteomics analysis were loaded onto Evotips (Evosep) as per the manufacturer's instructions. For whole proteome analysis, 200 ng peptide was loaded, while for pulldown sample, half of the sample collected was loaded for analysis.

*For whole proteome experiments:*

Peptides were analysed by nanoLC-MS/MS using an Evosep One (Evosep) coupled with a timsTOF HT (Bruker) equipped with an 8 cm × 150 µm, 1.5 µm analytical column (Evosep). 200 ng peptides were separated by the Evosep 60SPD workflow (Analytical solvents A: 0.1% FA and B: acetonitrile 17 plus 0.1% FA). Column was held at 40 °C. Data were acquired in data-independent acquisition (DIA) PASEF mode with the following settings: *m/z* range from 100 *m/z* to 1700 *m/z*, ion mobility range from 1/K0 = 1.30 to 0.85 Vs/cm$^2$ using equal ion accumulation and ramp times in the dual TIMS analyser of 100 ms each. Each cycle consisted of 8 PASEF ramps covering 21 mass steps each with 25 Da windows each with 2/3 non-overlapping ion mobility windows covering the 475 to 1000 *m/z* range and 0.85 and 1.26 Vs/cm2 ion mobility range. The collision energy was lowered as a function of increasing ion mobility from 59 eV at 1/K0 = 1.6 Vs/cm$^2$ to 20 eV at 1/K0 = 0.6 Vs/cm$^2$.

*For pulldown experiments:*

Peptides were analysed by nanoLC-MS/MS using an Evosep One (Evosep) coupled with a timsTOF HT (Bruker) equipped with an 8 cm × 150 µm, 1.5 µm analytical column (Evosep). 200 ng peptides were separated by the Evosep 60SPD workflow (Analytical solvents A: 0.1% FA and B: acetonitrile plus 0.1% FA). Column was held at 40 °C. Data were acquired in data-dependent acquisition (DDA) PASEF mode with the following settings: *m/z* range from 100 *m/z* to 1700 *m/z*, ion mobility range from 1/K0 = 1.30 to 0.85 Vs/cm$^2$ using equal ion accumulation and ramp times in the dual TIMS analyzer of 100 ms each. The collision energy was lowered as a function of increasing ion mobility from 59 eV at 1/K0 = 1.6 Vs/cm$^2$ to 20 eV at 1/K0 = 0.6 Vs/cm$^2$. Isolation width was lowered as a function of decreasing m/s from 3 *m/z* at 800 *m/z* to 2 *m/z* at 700 *m/z*. Active exclusion was applied for 0.4 min. Each cycle consisted of 4 PASEF ramps (total cycle time 0.53 s) with 2.75 ms measuring time allowed for each selected precursor.

### Data processing and downstream analysis

*For whole proteome experiments:*

diaPASEF Bruker.d files were processed using library-free analysis in DIA-NN (version 2.3.1) using the following parameters: Human database (Downloaded from UniProt 24 February 2026 containing 20518 proteins and 125 common contaminants); "deep learning-based spectra and RTs prediction" was enabled; trypsin with 1 missed cleavages; N-term Excision, C carbamidomethylation, Oxidation and N terminal Acetylation were enabled with maximum 2 variable modifications; MBR was enabled; quantification strategy set to "QuantUMS (precision)"; protein inference was enabled; Mass and MS1 accuracy set to 0.

Data analysis was carried out using Perseus version 2.1.5.0. Intensities were loaded for analysis. A text filter was applied to remove contaminants and data was $Log_{10}$ transformed. Rows were annotated to group replicates, and data was filtered for valid values in all replicates. Protein intensities were averaged and S-plots generated in GraphPad Prism version 11.

*For pulldown experiments:*

ddaPASEF Bruker .d files were processed using Fragpipe version 23.0 (Nesvilab). Data was searched against a human reference proteome with isoforms (Uniprot, UP000005640, accessed 23 February 2026, 20464 proteins) with 50% decoys and contaminants added. The built-in label-free quantification-match between runs (LFQ-MBR) workflow was used. Bruker .d files were searched using MSFragger (version 4.4.1) The following parameters were used: strict trypsin digestion; a maximum of 2 missed cleavages allowed; precursor ion tolerance of 20 ppm; oxidation (M) and N terminal acetylation as variable modifications; carbamidomethylation (C) as a fixed modification. MaxLFQ minimum ions was set to 1 and the retention time tolerance for match between runs was set to 1 min. All other default parameters were used for processing. MSFragger search results were processed using Percolator (version 3.7.1) for peptide-spectrum match validation, followed by Philosopher (version 5.1.1) for protein and FDR 18 filtering. Label-free quantification values were calculated using the MaxLFQ algorithm using IonQuant (version 1.11.20) with match between runs enabled and min ions set as 1.

Data analysis was carried out using Perseus version 2.1.5.0. MaxLFQ intensities (competition volcano plots) or iBAQ intensities (target enrichment S-plots) were loaded for analysis. For competition volcano plots, rows were filtered based on combined total peptide >1, a text filter was applied to remove contaminants and data was $Log_2$ transformed. Rows were annotated to group replicates, and data was filtered for 4/4 valid values in both conditions. Data was normalized by subtracting the median from the columns. Volcano plots were generated by plotting the fold-changes of the protein MaxLFQ intensities against the -$Log_{10}$(p-value). These were calculated by carrying out a two-sample t-test of the intensity values for each protein in the two conditions (permutation-based FDR = 0.05, S0 = 0.1). Data were plotted in GraphPad Prism version 11. For target enrichment S-plots rows were filtered based on combined total peptide >1, a text filter was applied to remove contaminants and data was $Log_2$ transformed. Rows were annotated to group replicates, and data was filtered for 4/4 valid values at least one condition. Proteins not significantly enriched (fold change >1, -$Log_{10}$(p-value) >1.3) in probe treated samples were removed. Enriched protein iBAQ intensities were averaged and S-plots generated in GraphPad Prism version 11.

**Activity of RAB27A/B inhibitors in melan-a melanocytes**

Melan-a murine melanocytes were seeded into 96-well plates (Merck, UK) at $5x10^3$ cells/well and cultured in standard conditions for 24 h (RPMI 1640 medium supplemented with 10% fetal calf serum, 2 mM glutamine, 100 U/ml penicillin G, 100 mg/ml streptomycin, and 200 nM phorbol 12-myristate 13-acetate (all Sigma-Aldrich, Poole, UK) in a humidified incubator at 37 °C with 10% $CO_2$). Compounds, diluted in growth medium (final 0.3% DMSO), were added to cells, cultured as above for 24 h and the effects on melanosome dispersion recorded using a Zoe cell imager (Bio-rad UK 1450031). Cellular melanosome distribution was scored as either ‘clustered’ or ‘dispersed’, dependent upon whether melanosomes were distributed in the perinuclear cytoplasm or isotropically distributed throughout the cytoplasm as determined by visual inspection.

**Viability assay in melan-a melanocytes**

Melan-a mouse melanocytes were seeded into 96-well plates (Merck, UK) at 5 x $10^3$ cells/well and cultured in standard conditions for 24 h (RPMI 1640 medium supplemented with 10% fetal calf serum, 2 mM glutamine, 100 U/ml penicillin G, 100 mg/ml streptomycin, and 200 nM phorbol 12-myristate 13-acetate (all Sigma-Aldrich, Poole, UK) in a humidified incubator at 37 °C with 10% $CO_2$). Compounds, diluted in growth medium, were added to cells, cultured as above for 24 h and the effects on viability were recorded using a Prestoblue assay (ThermoFisher A13262) according to the manufacturer protocol and using an ID3 microplate reader (Spectramax 412379).

### Organic Synthesis

#### General procedure A – Sulfonamide formation

To a solution of the specified amine in $CH_2Cl_2$ or THF (1 eq., 0.1–0.5 M) cooled to 0 °C, were added DIPEA or $Et_3N$ (2–3 eq.) and the specified sulfonyl chloride (1 eq.). The reaction was allowed to warm to rt and quenched with $H_2O$ after 1–16 h. The aqueous layer was extracted with ethyl acetate (3x, 20–100 mL), washed with brine (20–100 mL), dried ($MgSO_4$), filtered and concentrated *in vacuo*. The crude product was used without further purification unless specified.

#### General procedure B – Hydrogenation

To a stirring solution of the nitrosulfonamide in EtOAc and/or MeOH (1 eq, 0.05–0.2 M) was added 10% Pd/C (10% w/w). The flask was evacuated and refilled with a $H_2$ balloon (1 atm) three times and left stirring at rt for 16–72 h. The reaction mixture was filtered through celite with MeOH and the filtrate concentrated *in vacuo*. The crude product was used without further purification unless specified.

#### General procedure C – Acryloylation

To a solution of the specified aniline in anhydrous $CH_2Cl_2$ or THF (1 eq., 0.1–0.5 M) cooled to 0 °C, were added $Et_3N$ (1.5–3 eq.) and acryloyl chloride (1.1–3 eq.). The reaction was allowed to warm to rt and quenched with $H_2O$ after 1–16 h. The aqueous layer was extracted with ethyl acetate (3x, 20–100 mL), washed with brine (20–100 mL), dried ($MgSO_4$), filtered and concentrated *in vacuo*. The crude product was purified by liquid chromatography as specified.

#### General procedure D – Saponification

To a stirring solution of the specified ester in dioxane (1 eq., 0.1–0.5 M) was added a solution of LiOH (1.1–3.0 eq) in $H_2O$ (0.1–0.5 M). The reaction was stirred at rt for 1–16 h, then diluted with $H_2O$ and acidified with 1 M HCl until pH~3. The aqueous layer was extracted with EtOAc (3x, 20–100 mL), dried ($MgSO_4$), filtered and concentrated *in vacuo*. The crude product was used without further purification.

#### General procedure E – Amide coupling

To a stirring solution of the specified acid in $CH_2Cl_2$ or DMF (1 eq., 0.1–0.5 M) was added EDCI or HATU or HBTU (1.0–1.5 eq), DIPEA (3.0–4 eq) and the specified amine (1.0–1.5 eq). The reaction was stirred at rt for 1–16 h, then diluted with water (10–100 mL) and extracted with ethyl acetate (3x, 20–100 mL), washed with brine (20–100 mL), dried ($MgSO_4$), filtered and concentrated *in vacuo*. The crude product was purified by liquid chromatography as specified. In the case of parallel syntheses, the reactions were carried out in DMF and directly purified by reverse phase chromatography (5–100% MeCN in $H_2O$ with 0.1% FA) to afford corresponding amides.

**1-((4-Nitrophenyl)sulfonyl)pyrrolidine (15)**

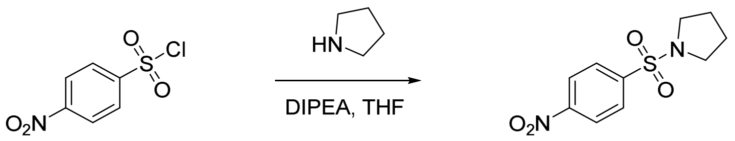

*General procedure A, reagents:* pyrrolidine (120 µL, 1.4 mmol); 4-nitrophenylsulfonyl chloride (300 mg, 1.4 mmol); DIPEA (470 µL, 2.7 mmol); THF (3.0 mL).
*Yield:* 350 mg, quantitative, yellow solid.
**$^1$H NMR** (400 MHz, DMSO-d6) δ: 8.50 – 8.36 (m, 2H), 8.19 – 7.99 (m, 2H), 3.26 – 3.12 (m, 4H), 1.70 – 1.64 (m, 4H) ppm.
**$^{13}$C NMR** (101 MHz, DMSO-*d*$_6$) δ: 150.4, 142.3, 129.3, 125.1, 48.4, 25.3 ppm.
**LRMS** (+ESI) *m/z*: $[M+H]^+$ 257.

**4-(Pyrrolidin-1-ylsulfonyl)aniline (16)**

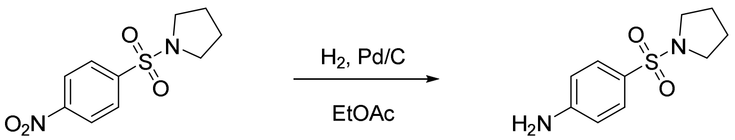

*General procedure B, reagents:* nitrosulfonamide **15** (150 mg, 0.58 mmol); $H_2$ (1 atm); 10% Pd/C (18 mg, 10% w/w); EtOAc (4.4 mL).
*Yield:* 110 mg, 85%, white solid.
**$^1$H NMR** (400 MHz, Chloroform-*d*) δ: 7.66 – 7.58 (m, 2H), 6.77 – 6.64 (m, 2H), 4.16 (s, 2H), 3.25 – 3.17 (m, 4H), 1.81 – 1.72 (m, 4H) ppm.
**$^{13}$C NMR** (101 MHz, Chloroform-*d*) δ: 150.5, 129.7, 125.1, 114.0, 47.9, 25.2 ppm.
**LRMS** (+ESI) *m/z*: $[M+H]^+$ 227.

***N*-(4-(Pyrrolidin-1-ylsulfonyl)phenyl)acrylamide (1, IMP-1704)**

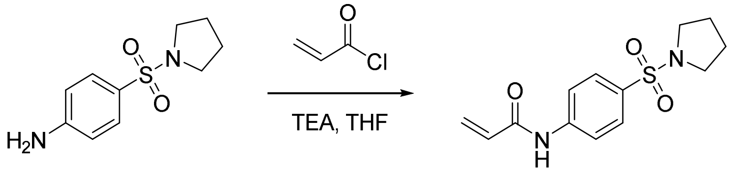

*General procedure C, reagents:* aniline **16** (100 mg, 0.44 mmol); acryloyl chloride (36 µL, 0.44 mmol); $Et_3N$ (90 µL, 0.65 mmol); THF (3.0 mL); *purification:* LC (2x, 0–5% MeOH in $CH_2Cl_2$).
*Yield:* 50 mg, 41%, white solid.
**$^1$H NMR** (400 MHz, DMSO-*d*$_6$) δ: 10.57 (s, 1H), 7.96 – 7.86 (m, 2H), 7.78 (m, 2H), 6.47 (dd, $J$ = 16.9, 10.0 Hz, 1H), 6.32 (dd, $J$ = 17.0, 2.0 Hz, 1H), 5.84 (dd, $J$ = 10.0, 2.0 Hz, 1H), 3.16 – 3.08 (m, 4H), 1.70 – 1.59 (m, 4H) ppm.
**$^{13}$C NMR** (101 MHz, DMSO-*d*$_6$) δ: 164.2, 143.5, 131.9, 130.7, 129.1, 128.6, 119.6, 48.3, 25.1 ppm.
**LRMS** (+ESI) *m/z*: $[M+H]^+$ 281.
**HRMS** (ESI) *m/z*: $[M+H]^+$ calcd for $C_{13}H_{17}N_2O_3S$: 281.0960; found: 281.0966.
**HPLC** retention time 4.0 min, >99.0%.

**(4-Nitrophenyl)(pyrrolidin-1-yl)methanone (17)**

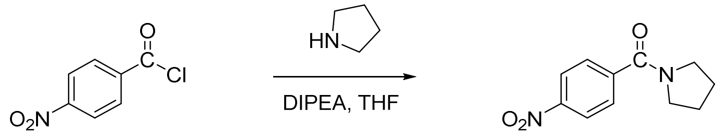

*General procedure A, reagents:* pyrrolidine (140 µL, 1.6 mmol); 4-nitrobenzoyl chloride (300 mg, 1.4 mmol); DIPEA (560 µL, 3.2 mmol); THF (3.0 mL).
*Yield:* 340 mg, 95%, orange solid.
**$^1$H NMR** (400 MHz, Chloroform-*d*) δ: 8.36 – 8.23 (m, 2H), 7.74 – 7.64 (m, 2H), 3.69 (t, *J* = 6.9 Hz, 2H), 3.40 (t, *J* = 6.6 Hz, 2H), 2.08 – 1.86 (m, 5H) ppm.
**$^{13}$C NMR** (101 MHz, Chloroform-*d*) δ 167.4, 148.4, 143.2, 128.2, 123.7, 49.5, 46.4, 26.4, 24.4 ppm.

**(4-Aminophenyl)(pyrrolidin-1-yl)methanone (18)**

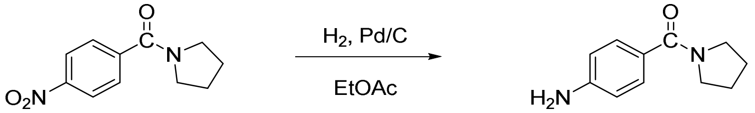

*General procedure B, reagents:* nitrobenzamide **17** (330 mg, 1.5 mmol); $H_2$ (1 atm); 10% Pd/C (30 mg, 10% w/w); EtOAc (3.0 mL).
*Yield:* 290 mg, quantitative, light orange solid.
**$^1$H NMR** (400 MHz, DMSO-*d*$_6$) δ: 7.32 – 7.20 (m, 2H), 6.59 – 6.46 (m, 2H), 5.51 (s, 2H), 3.43 (t, *J* = 6.0 Hz, 4H), 1.80 (t, *J* = 5.5 Hz, 4H) ppm.
**$^{13}$C NMR** (101 MHz, DMSO-*d*$_6$) δ: 169.0, 151.0, 129.6, 123.9, 112.9, 49.7, 46.6, 29.4, 24.4 ppm.

***N*-(4-(Pyrrolidine-1-carbonyl)phenyl)acrylamide (2)**

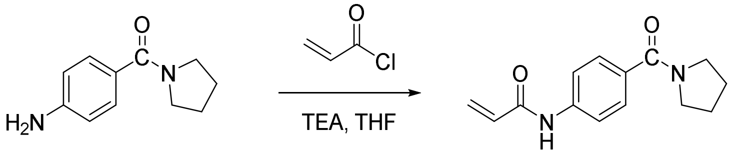

*General procedure C, reagents:* aniline **18** (100 mg, 0.53 mmol); acryloyl chloride (65 µL, 0.79 mmol); $Et_3N$ (110 µL, 0.79 mmol); THF (1.5 mL); DMF (0.5 mL); *purification:* LC (0–10% MeOH in $CH_2Cl_2$).
*Yield:* 54 mg, 42%, white solid.
**$^1$H NMR** (400 MHz, DMSO-*d*$_6$) δ: 10.33 (s, 1H), 7.77 – 7.67 (m, 2H), 7.57 – 7.46 (m, 2H), 6.45 (dd, *J* = 17.0, 10.1 Hz, 1H), 6.29 (dd, *J* = 17.0, 2.1 Hz, 1H), 5.79 (dd, *J* = 10.1, 2.0 Hz, 1H), 3.43 (dd, *J* = 10.5, 4.7 Hz, 4H), 1.94 – 1.68 (m, 4H) ppm.
**$^{13}$C NMR** (101 MHz, DMSO-*d*$_6$) δ: 168.2, 163.8, 140.8, 132.4, 132.2, 128.6, 127.8, 119.0, 49.5, 46.5, 26.5, 24.4 ppm.
**LRMS** (+ESI) *m/z*: $[M+H]^+$ 245.
**HRMS** (ESI) *m/z*: $[M+H]^+$ calcd for $C_{14}H_{17}N_2O_2$: 245.1290; found: 245.1283.
**HPLC** retention time 3.2 min, >99.0%.

**1-((3-Nitrophenyl)sulfonyl)pyrrolidine (19)**

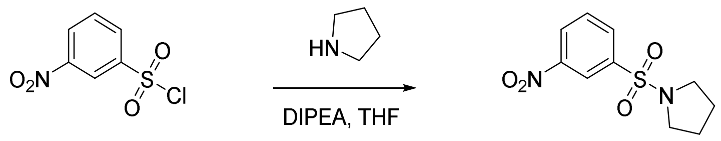

*General procedure A, reagents:* pyrrolidine (110 µL, 1.4 mmol); 3-nitrophenylsulfonyl chloride (300 mg, 1.4 mmol); DIPEA (470 µL, 2.7 mmol); THF (3.0 mL).
*Yield:* 260 mg, 75%, pale orange solid.
**$^1$H NMR** (400 MHz, DMSO-*d*$_6$) δ: 8.54 (ddd, *J* = 8.3, 2.3, 1.0 Hz, 1H), 8.42 (t, *J* = 2.0 Hz, 1H), 8.26 (ddd, *J* = 7.8, 1.8, 1.0 Hz, 1H), 7.94 (t, *J* = 8.0 Hz, 1H), 3.24 – 3.15 (m, 4H), 1.74 – 1.61 (m, 4H) ppm.
**$^{13}$C NMR** (101 MHz, DMSO-*d*$_6$) δ: 148.5, 138.4, 133.6, 132.0, 128.0, 122.4, 48.4, 25.3 ppm.
**LRMS** (+ESI) *m/z*: $[M+H]^+$ 257.

**3-(Pyrrolidin-1-ylsulfonyl)aniline (20)**

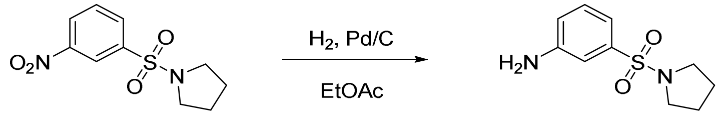

*General procedure B, reagents:* nitrosulfonamide **20** (250 mg, 0.98 mmol); $H_2$ (1 atm); 10% Pd/C (25 mg, 10% w/w); EtOAc (6 mL).
*Yield:* 220 mg, 99%, white solid.
**$^1$H NMR** (400 MHz, DMSO-*d*$_6$) δ: 7.23 (t, *J* = 7.9 Hz, 1H), 6.98 (t, *J* = 2.0 Hz, 1H), 6.85 (ddd, *J* = 7.7, 1.9, 0.9 Hz, 1H), 6.80 (ddd, *J* = 8.2, 2.3, 1.0 Hz, 1H), 3.19 – 3.05 (m, 4H), 1.70 – 1.61 (m, 4H) ppm.
**$^{13}$C NMR** (101 MHz, DMSO-*d*$_6$) δ 150.0, 136.9, 130.1, 118.2, 114.4, 112.2, 48.2, 25.2 ppm.

***N*-(3-(Pyrrolidin-1-ylsulfonyl)phenyl)acrylamide (3)**

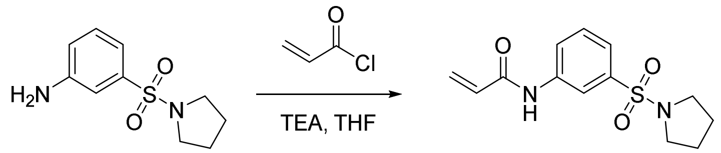

*General procedure C, reagents:* aniline **20** (100 mg, 0.44 mmol); acryloyl chloride (50 µL, 0.65 mmol); $Et_3N$ (90 µL, 0.65 mmol); THF (1.5 mL).
*Yield:* 110 mg, 91%, white solid.
**$^1$H NMR** (400 MHz, DMSO-*d*$_6$) δ: 10.51 (s, 1H), 8.22 (t, *J* = 1.9 Hz, 1H), 7.92 (ddd, *J* = 8.2, 2.1, 1.0 Hz, 1H), 7.59 (t, *J* = 8.0 Hz, 1H), 7.49 (ddd, *J* = 7.8, 1.8, 1.0 Hz, 1H), 6.43 (dd, *J* = 17.0, 9.9 Hz, 1H), 6.31 (dd, *J* = 16.9, 2.1 Hz, 1H), 5.82 (dd, *J* = 10.0, 2.1 Hz, 1H), 3.23 – 3.08 (m, 4H), 1.78 – 1.53 (m, 4H) ppm.
**$^{13}$C NMR** (101 MHz, DMSO-*d*$_6$) δ: 164.1, 140.2, 137.1, 131.9, 130.5, 128.3, 123.6, 122.4, 118.1, 48.3, 25.2 ppm.
**LRMS** (+ESI) *m/z*: $[M+H]^+$ 281.
**HRMS** (−ESI) *m/z*: $[M-H]^-$ calcd for $C_{13}H_{15}N_2O_3S$: 279.0803; found: 279.0812.
**HPLC** retention time 3.6 min, 98.2%.

**Methyl ((4-nitrophenyl)sulfonyl)-*L*-prolinate (21)**

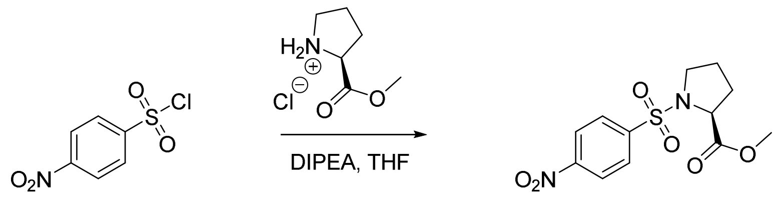


*General procedure A, reagents:* methyl L-prolinate hydrochloride (750 mg, 4.5 mmol); 4-nitrophenylsulfonyl chloride (1.0 g, 4.5 mmol); DIPEA (2.4 mL, 14 mmol); THF (5.0 mL).
*Yield:* 1.4 g, 97%, orange oil.
**$^1$H NMR** (400 MHz, Chloroform-*d*) δ: 8.42 – 8.37 (m, 2H), 8.15 – 8.07 (m, 2H), 4.48 (dd, *J* = 8.5, 3.7 Hz, 1H), 3.73 (s, 3H), 3.47 (dd, *J* = 7.4, 6.0 Hz, 2H), 2.26 – 2.13 (m, 1H), 2.12 – 1.87 (m, 3H) ppm.
**$^{13}$C NMR** (101 MHz, Chloroform-*d*) δ: 172.2, 144.7, 128.7, 124.2, 60.6, 52.6, 48.3, 31.0, 24.8 ppm.
**LRMS** (+ESI) *m/z*: $[M+Na]^+$ 337.

**Methyl ((4-aminophenyl)sulfonyl)-*L*-prolinate (22)**

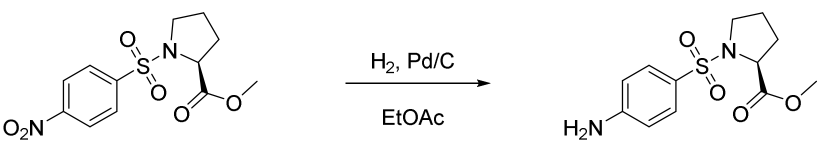


*General procedure B, reagents:* nitrosulfonamide **21** (1.4 g, 4.3 mmol); $H_2$ (1 atm); 10% Pd/C (140 mg, 10% w/w); EtOAc (6 mL).
*Yield:* 1.1 g, 85%, light pink solid.
**$^1$H NMR** (400 MHz, DMSO-*d*$_6$) δ: 7.48 – 7.40 (m, 2H), 6.69 – 6.57 (m, 2H), 6.09 (s, 2H), 4.05 (dd, *J* = 8.5, 4.4 Hz, 1H), 3.65 (s, 3H), 3.34 – 3.26 (m, 1H), 3.09 (m, 1H), 2.08 – 1.72 (m, 3H), 1.67 – 1.46 (m, 1H) ppm.
**$^{13}$C NMR** (101 MHz, DMSO-*d*$_6$) δ: 172.9, 153.7, 129.7, 122.0, 113.3, 60.6, 52.5, 48.9, 30.8, 24.7 ppm.
**LRMS** (+ESI) *m/z*: $[M+H]^+$ 285.

**Methyl ((4-acrylamidophenyl)sulfonyl)-*L*-prolinate (4, IMP1712)**

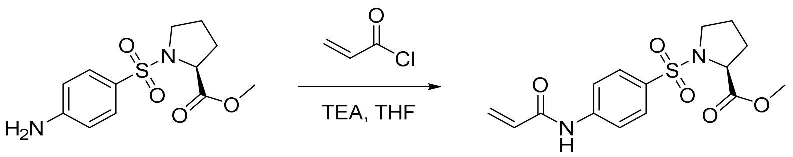


*General procedure C, reagents:* aniline **22** (710 mg, 2.5 mmol); acryloyl chloride (300 µL, 3.3 mmol); $Et_3N$ (520 µL, 3.8 mmol); THF (5.0 mL); *purification:* LC (10–80% EtOAc in hexane).
*Yield:* 510 mg, 60%, foamy transparent oil.
**$^1$H NMR** (400 MHz, Chloroform-*d*) δ: 8.12 (s, 1H), 7.92 – 7.65 (m, 4H), 6.47 (dt, *J* = 16.9, 1.1 Hz, 1H), 6.32 (dd, *J* = 16.8, 10.1 Hz, 1H), 5.82 (dt, *J* = 10.1, 1.2 Hz, 1H), 4.27 (m, 1H), 3.71 (d, *J* = 0.8 Hz, 3H), 3.48 (m, 1H), 3.28 (m, 1H), 2.16 – 1.86 (m, 3H), 1.85 – 1.66 (m, 2H) ppm.
**$^{13}$C NMR** (101 MHz, Chloroform-*d*) δ: 172.7, 164.0, 142.3, 132.7, 130.7, 129.1, 128.7, 119.7, 60.5, 52.6, 48.6, 31.0, 24.7 ppm.
**LRMS** (+ESI) *m/z*: $[M+H]^+$ 339.
**HRMS** (ESI) *m/z*: $[M+H]^+$ calcd for $C_{15}H_{19}N_2O_5S$: 339.1015; found: 339.1020.
**HPLC** retention time 4.0 min, 98.9%.

**Methyl ((4-nitrophenyl)sulfonyl)-*D*-prolinate (23)**

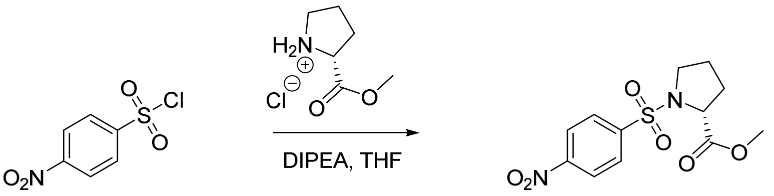


*General procedure A, reagents:* methyl D-prolinate hydrochloride (75 mg, 0.45 mmol); 4-nitrophenylsulfonyl chloride (100 mg, 0.45 mmol); $Et_3N$ (120 µL, 0.9 mmol); THF (3.0 mL).
*Yield:* 120 mg, 82%, yellow solid.
**$^1$H NMR** (400 MHz, Chloroform-*d*) δ: 8.41 – 8.35 (m, 2H), 8.12 – 8.06 (m, 2H), 4.50 – 4.44 (m, 1H), 3.73 (s, 3H), 3.50 – 3.43 (m, 2H), 2.22 – 2.14 (m, 1H), 2.09 – 1.91 (m, 3H) ppm.
**$^{13}$C NMR** (101 MHz, Chloroform-*d*) δ: 172.1, 144.6, 128.8, 124.2, 60.5, 52.5, 48.3, 30.9, 24.7 ppm.
**LRMS** (+ESI) *m/z*: $[M+Na]^+$ 337.

**Methyl ((4-aminophenyl)sulfonyl)-*D*-prolinate (24)**

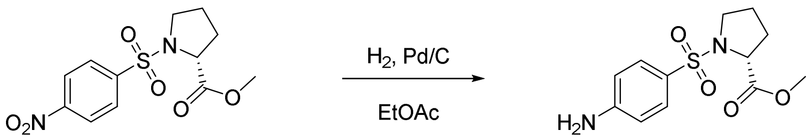


*General procedure B, reagents:* nitrosulfonamide **23** (120 mg, 0.36 mmol); $H_2$ (1 atm); 10% Pd/C (12 mg, 10% w/w); EtOAc (6 mL).
*Yield:* 110 mg, quantitative, yellow solid.
**$^1$H NMR** (400 MHz, DMSO-$d_6$) δ: 7.68 – 7.60 (m, 2H), 6.74-6.67 (m, 2H), 4.27 – 4.21 (m,1H), 3.74 (s, 3H), 3.51 – 3.44 (m, 1H), 3.31 – 3.23 (m, 1H), 2.05 – 1.89 (m, 3H), 1.79 – 1.69 (m, 1H) ppm.
**LRMS** (+ESI) *m/z*: $[M+H]^+$ 285.

**Methyl ((4-acrylamidophenyl)sulfonyl)-*D*-prolinate (5)**

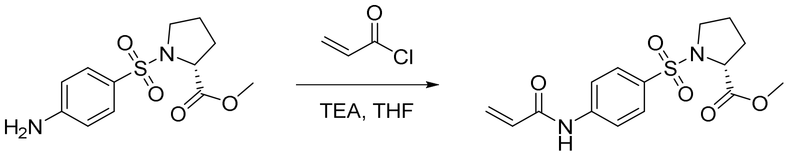


*General procedure C, reagents:* aniline **24** (110 mg, 0.36 mmol); acryloyl chloride (300 µL, 3.3 mmol); $Et_3N$ (520 µL, 3.8 mmol); THF (5.0 mL); *purification:* LC (20–50% EtOAc in hexane).
*Yield:* 95 mg, 76%, yellow oil.
**$^1$H NMR** (400 MHz, Chloroform – *d*) δ: 8.14 (br s, 1H), 7.86 – 7.78 (m, 4H), 6.53 – 6.44 (m, 1H), 6.39 – 6.29 (m, 1H), 5.87 – 5.80 (m, 1H), 4.33 – 4.25 (m, 1H), 3.74 (s, 3H), 3.55 – 3.45 (m, 1H), 3.35 – 3.25 (m, 1H), 2.06 – 1.95 (m, 3H), 1.82 – 1.72 (m, 1H) ppm.
**$^{13}$C NMR** (101 MHz, Chloroform-*d*) δ: 172.6, 163.9, 142.2, 132.8, 130.6, 129.0, 128.7, 119.7, 60.4, 52.5, 48.5, 30.9, 24.6 ppm.
**LRMS** (+ESI) *m/z*: $[M+H]^+$ 339.
**HRMS** (ESI) *m/z*: $[M+H]^+$ calcd for $C_{15}H_{19}N_2O_5S$: 339.1015; found: 339.1020.
**HPLC** retention time 3.9 min, 98.1%.

**((4-Acrylamidophenyl)sulfonyl)-*L*-proline (25)**

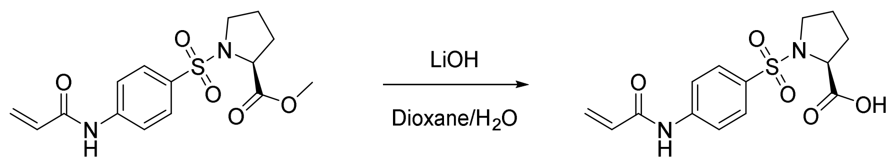


*General procedure D, reagents:* ester **4** (100 mg, 0.30 mmol); LiOH.$H_2O$ (14 mg, 0.30 mmol); dioxane (0.9 mL); $H_2O$ (0.9 mL).
*Yield:* 91 mg, 95%, white solid.
**$^1$H NMR** (400 MHz, DMSO-*d*$_6$) δ: 12.76 (s, 1H), 10.58 (s, 1H), 7.97 – 7.86 (m, 2H), 7.85 – 7.73 (m, 2H), 6.47 (dd, *J* = 16.9, 10.0 Hz, 1H), 6.32 (dd, *J* = 17.0, 2.0 Hz, 1H), 5.84 (dd, *J* = 10.0, 2.0 Hz, 1H), 4.08 (dd, *J* = 8.2, 4.4 Hz, 1H), 3.21 – 3.11 (m, 1H), 1.96 – 1.70 (m, 3H), 1.65 – 1.44 (m, 1H) ppm.
**$^{13}$C NMR** (101 MHz, DMSO-*d*$_6$) δ: 173.7, 164.2, 143.5, 132.0, 131.9, 128.9, 128.6, 119.6, 60.9, 48.9, 30.9, 24.7 ppm.
**LRMS** (+ESI) *m/z*: $[M+H]^+$ 325.

**(*S*)-1-((4-Acrylamidophenyl)sulfonyl)-*N*-(prop-2-yn-1-yl)pyrrolidine-2-carboxamide (6, C05)**

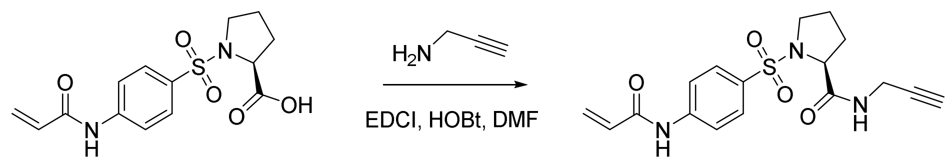


To a stirring solution of acid **25** (85 mg, 0.26 mmol) in anhydrous DMF (1.0 mL) was added HOBt (36 mg, 0.26 mmol) and EDCI.HCl (75 mg, 0.39 mmol). After 10 min, propargylamine (20 µL, 0.31 mmol) was added. After reaction was diluted with water (50 mL) and extracted with ethyl acetate (3x, 10 mL), washed with brine (10 mL), dried ($MgSO_4$), filtered and concentrated *in vacuo*. The crude product was purified by liquid chromatography (0–5% MeOH in $CH_2Cl_2$).
*Yield:* 56 mg, 59%, white solid.
**$^1$H NMR** (400 MHz, DMSO-*d*$_6$) δ: 10.58 (s, 1H), 8.43 (t, *J* = 5.6 Hz, 1H), 7.96 – 7.88 (m, 2H), 7.86 – 7.80 (m, 2H), 6.47 (dd, *J* = 17.0, 10.0 Hz, 1H), 6.33 (dd, *J* = 16.9, 2.0 Hz, 1H), 5.84 (dd, *J* = 10.0, 2.0 Hz, 1H), 4.03 (dd, *J* = 8.2, 3.5 Hz, 1H), 3.88 (qdd, *J* = 17.4, 5.6, 2.5 Hz, 2H), 3.42 (ddd, *J* = 10.1, 6.8, 4.9 Hz, 1H), 3.20 – 3.13 (m, 1H), 3.12 (t, *J* = 2.5 Hz, 1H), 1.84 – 1.62 (m, 3H), 1.58 – 1.41 (m, 1H) ppm.
**$^{13}$C NMR** (101 MHz, DMSO-*d*$_6$) δ: 171.5, 164.2, 143.7, 131.9, 131.1, 129.1, 128.6, 119.6, 81.5, 73.4, 61.9, 49.6, 31.1, 28.6, 24.5 ppm.
**LRMS** (+ESI) *m/z*: $[M+H]^+$ 362.
**HRMS** (ESI) *m/z*: $[M+H]^+$ calcd for $C_{17}H_{20}N_3O_4S$: 362.1175; found: 362.1185.
**HPLC** retention time 3.4 min, >99.0%.

***tert*-Butyl (*S*)-2-((4-methoxyphenyl)carbamoyl)pyrrolidine-1-carboxylate (26)**

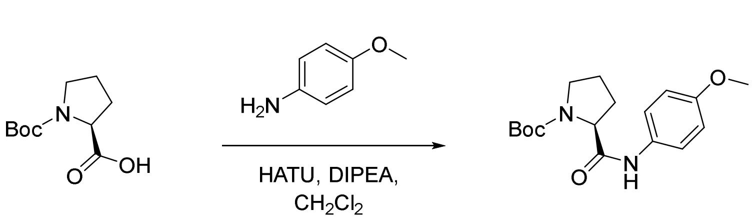

*General procedure E, reagents:* Boc-proline (2.0 g, 9.3 mmol); 4-methoxyaniline (1.7 g, 14.0 mmol); HATU (4.2 g, 11 mmol); DIPEA (4.8 mL, 28 mmol); $CH_2Cl_2$ (20 mL); *purification:* LC (2x, 2.5% MeOH in $CH_2Cl_2$).
*Yield:* 3.1 g (70% pure), 92%, orange crystalline solid.
**$^1$H NMR** (400 MHz, Chloroform-*d*) δ: 9.32 (s, 1H), 7.50 – 7.37 (m, 2H), 6.95 – 6.83 (m, 2H), 4.48 (s, 1H), 3.80 (s, 3H), 3.51 – 3.22 (m, 2H), 2.63 – 2.43 (m, 1H), 2.04 – 1.81 (m, 2H), 1.72 – 1.64 (m, 1H), 1.51 (s, 9H) ppm.
**$^{13}$C NMR** (101 MHz, Chloroform-*d*) δ: 152.7, 121.4, 121.3, 116.4, 114.8, 114.1, 80.9, 55.8, 55.5, 47.2, 28.4 ppm.

#### (*S*)-2-((4-Methoxyphenyl)carbamoyl)pyrrolidin-1-ium chloride (27)

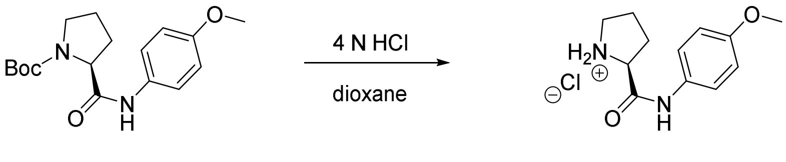


To a stirring solution of amide **26** (70%, 3.1 g, 9.7 mmol) in dioxane (8.0 mL) was added 4 N HCl in dioxane (3.0 mL). After 1 h additional 4 N HCl in dioxane (3.0 mL) and the reaction was stirred for 4 h at rt. The crude suspension was filtered and the precipitate washed with $CH_2Cl_2$ to afford the desired product which was used without further purification.
*Yield:* 2.3 g (70% pure), 83%, white solid.
**$^1$H NMR** (400 MHz, DMSO-*d*$_6$) δ: 10.83 (s, 1H), 10.17 (d, *J* = 45.0 Hz, 2H), 7.66 – 7.49 (m, 2H), 7.00 – 6.88 (m, 2H), 4.37 (s, 1H), 3.74 (s, 3H), 3.26 (d, *J* = 7.4 Hz, 3H), 2.42 (qt, *J* = 10.4, 2.5 Hz, 1H), 2.00 – 1.87 (m, 3H) ppm.
**$^{13}$C NMR** (101 MHz, DMSO-*d*$_6$) δ: 166.7, 156.2, 131.8, 124.9, 124.8, 121.5, 115.3, 114.5, 59.9, 55.7, 46.1, 30.2, 24.1 ppm.

#### (*S*)-*N*-(4-Methoxyphenyl)-1-((4-nitrophenyl)sulfonyl)pyrrolidine-2-carboxamide (28)

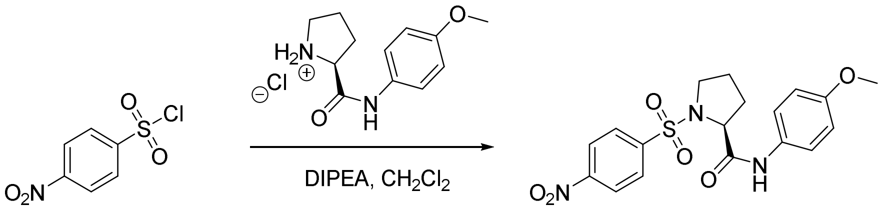


*General procedure A, reagents:* amine **27** (3.4 g, 13 mmol); 4-nitrophenylsulfonyl chloride (3.0 g, 13 mmol); DIPEA (7.0 mL, 40 mmol); $CH_2Cl_2$ (10 mL).
*Yield:* 4.3 g, 80%, orange solid.
**$^1$H NMR** (400 MHz, Chloroform-*d*) δ: 8.55 – 8.38 (m, 3H), 8.18 – 8.04 (m, 2H), 7.53 – 7.44 (m, 2H), 6.95 – 6.88 (m, 2H), 4.24 (dd, *J* = 8.6, 2.6 Hz, 1H), 3.72 (ddd, *J* = 10.4, 7.3, 3.4 Hz, 1H), 3.29 (td, *J* = 9.7, 6.7 Hz, 1H), 2.40 (ddd, *J* = 12.7, 6.4, 3.1 Hz, 1H), 1.94 (dddd, *J* = 15.9, 12.9, 7.2, 4.6 Hz, 1H), 1.84 – 1.54 (m, 2H) ppm.
**$^{13}$C NMR** (101 MHz, Chloroform-*d*) δ: 168.04, 156.85, 150.6, 141.57, 130.23, 129.1, 124.7, 121.9, 114.2, 63.1, 55.5, 50.2, 30.1, 24.5 ppm.

**(*S*)-1-((4-Aminophenyl)sulfonyl)-*N*-(4-methoxyphenyl)pyrrolidine-2-carboxamide (29)**

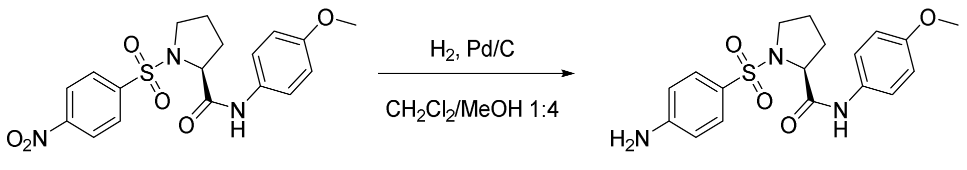


*General procedure B, reagents:* nitrosulfonamide **28** (9.2 g, 23 mmol); $H_2$ (1 atm); 10% Pd/C (920 mg, 10% w/w); $CH_2Cl_2$/MeOH (1:4, 150 mL).
*Yield:* 3.8 g, 45%, white solid.
**$^1$H NMR** (400 MHz, DMSO-$d_6$) δ: 9.69 (s, 1H), 7.62 – 7.40 (m, 4H), 6.95 – 6.78 (m, 2H), 6.71 – 6.57 (m, 2H), 6.08 (d, *J* = 4.9 Hz, 2H), 4.10 – 3.95 (m, 1H), 3.73 (s, 3H), 3.46 – 3.37 (m, 1H), 3.19 – 3.02 (m, 1H), 1.89 – 1.67 (m, 3H), 1.58 – 1.41 (m, 1H) ppm.

**(*S*)-1-((4-Acrylamidophenyl)sulfonyl)-*N*-(4-methoxyphenyl)pyrrolidine-2-carboxamide (7)**

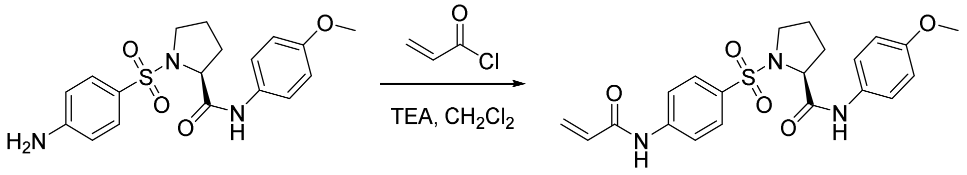


*General procedure C, reagents:* aniline **29** (60 mg, 0.15 mmol); acryloyl chloride (13 µL, 0.23 mmol, 2x); $Et_3N$ (67 µL, 0.47 mmol, 2x); $CH_2Cl_2$ (5.0 mL); *purification:* LC (40–100% EtOAc in hexane).
*Yield:* 31 mg, 44%, white crystalline solid.
**$^1$H NMR** (400 MHz, DMSO-$d_6$) δ: 10.59 (s, 1H), 9.81 (s, 1H), 7.95 – 7.88 (m, 2H), 7.88 – 7.77 (m, 2H), 7.56 – 7.47 (m, 2H), 6.93 – 6.84 (m, 2H), 6.47 (dd, *J* = 17.0, 10.0 Hz, 1H), 6.33 (dd, *J* = 16.9, 2.0 Hz, 1H), 5.85 (dd, *J* = 10.1, 2.0 Hz, 1H), 4.20 – 4.10 (m, 1H), 3.73 (s, 3H), 3.53 – 3.42 (m, 1H), 3.29 – 3.15 (m, 1H), 1.95 – 1.75 (m, 3H), 1.59 – 1.39 (m, 1H) ppm.
**$^{13}$C NMR** (101 MHz, Methanol-$d_4$) δ: 169.6, 163.7, 155.4, 143.2, 131.8, 131.4, 131.0, 128.6, 128.2, 121.2, 119.2, 113.8, 61.8, 55.2, 49.2, 31.1, 24.3 ppm.
**LRMS** (+ESI) *m/z*: $[M+H]^+$ 430.
**HRMS** (+ESI) *m/z*: $[M+H]^+$ calcd $C_{21}H_{24}N_3O_5S$ 430.1431, found 430.1425.
**HPLC** retention time 4.2 min, >99.0%.

**((4-Acrylamidophenyl)sulfonyl)-*D*-proline (30)**

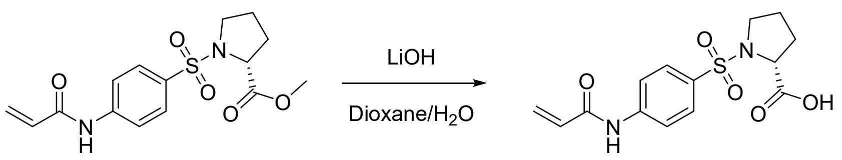


*General procedure D, reagents:* ester **5** (175 mg, 0.52 mmol); LiOH.$H_2O$ (65 mg, 1.55 mmol); dioxane (5.0 mL); $H_2O$ (2.5 mL).
*Yield:* 126 mg, 75%, colourless oil.
**LRMS** (+ESI) *m/z*: $[M+H]^+$ 325.

**(*R*)-1-((4-Acrylamidophenyl)sulfonyl)-*N*-(4-methoxyphenyl)pyrrolidine-2-carboxamide (8)**

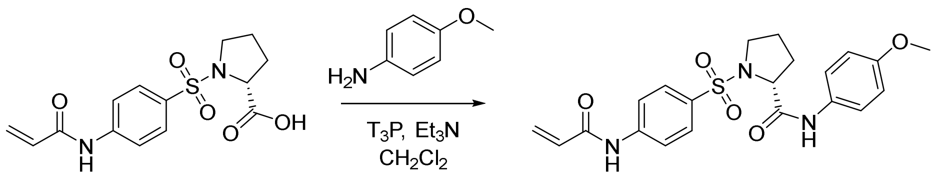


*General procedure E, reagents:* acid **30** (25 mg, 0.08 mmol); 4-methoxyaniline (14 mg, 0.12 mmol); $T_3P$ (50% in EtOAc, 98 µL, 0.15 mmol); $Et_3N$ (21 µL, 0.15 mmol); $CH_2Cl_2$ (1.0 mL); *purification:* LC (20–60% EtOAc in hexane).
*Yield:* 17 mg, 51%, white solid.
**$^1$H NMR** (400 MHz, DMSO-*d*$_6$) δ: 10.58 (s, 1H), 9.80 (s, 1H), 7.94 – 7.88 (m, 2H), 7.87 – 7.81 (m, 2H), 7.55 – 7.48 (m, 2H), 6.93 – 6.85(m, 2H), 6.46 (dd, J = 17.1, 9.9 Hz, 1H), 6.33 (dd, J = 17.1, 1.8 Hz, 1H), 5.84 (dd, J = 9.9, 1.8 Hz, 1H), 6.14 (dd, J = 7.3, 5.1 Hz, 1H), 3.51 – 3.42 (m, 1H), 3.25 – 3.17 (m, 1H), 1.92 – 1.77 (m, 3H), 1.58 – 1.44 (m, 1H) ppm.
**LRMS** (+ESI) *m/z*: $[M+H]^+$ 430.
**HRMS** (+ESI) *m/z*: $[M+H]^+$ calcd $C_{21}H_{24}N_3O_5S$ 430.1431, found 430.1454.
**HPLC** retention time 4.5 min, >99.0%.

***tert*-Butyl (4-(prop-2-yn-1-yloxy)phenyl)carbamate (31)**

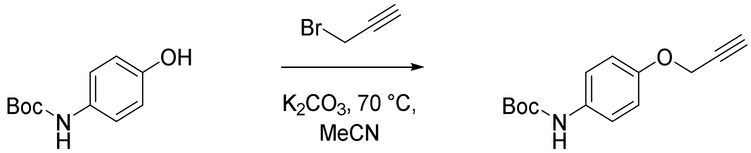


To a stirring solution of tert-butyl (4-hydroxyphenyl)carbamate (250 mg, 1.2 mmol) in MeCN (1.5 mL) in a microwave vial were added $K_2CO_3$ (250 mg, 1.8 mmol) and propargyl bromide (80% in toluene, 140 µL, 1.3 mmol). The vial was sealed and heated 70 °C, and the reaction was stirred for 14 h. The crude mixture was diluted with EtOAc (20 mL) and washed with H2O (15 mL). The aqueous phase was extracted with EtOAc (2x, 10 mL) and the combined organic phase was washed with brine (10 mL), dried ($MgSO_4$), filtered and concentrated *in vacuo* to afford the desired product which was used without further purification.
*Yield:* 300 mg, quantitative, orange oil.
**$^1$H NMR** (400 MHz, Chloroform-*d*) δ: 7.34 – 7.29 (m, 2H), 7.02 – 6.88 (m, 2H), 6.40 (s, 1H), 4.68 (d, *J* = 2.4 Hz, 2H), 2.53 (t, *J* = 2.4 Hz, 1H), 1.54 (s, 9H) ppm
**LRMS** (+ESI) *m/z*: $[M+H]^+$ 248.

**4-(Prop-2-yn-1-yloxy)benzenaminium chloride (32)**

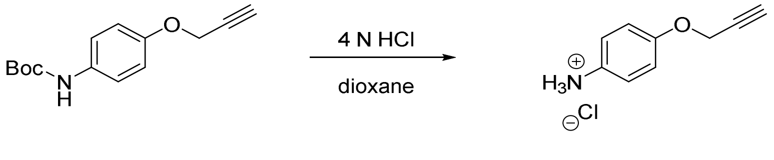


To a stirring solution of alkyne 31 (300 mg, 1.2 mmol) in dioxane (1.0 mL) cooled to 0 °C was added 4 N HCl in dioxane (1.5 mL). After 16 h the crude was concentrated *in vacuo* and azeotroped with MeOH (3x) to remove excess dioxane and afford the desired product which was used without further purification.

*Yield:* 180 mg, 80%, pale yellow solid.
**¹H NMR** (400 MHz, DMSO-$d_6$) δ: 10.22 (s, 3H), 7.40 – 7.29 (m, 2H), 7.14 – 7.00 (m, 2H), 4.83 (d, *J* = 2.3 Hz, 2H), 3.61 (t, *J* = 2.4 Hz, 1H). ppm

**(*S*)-1-((4-Acrylamidophenyl)sulfonyl)-*N*-(4-(prop-2-yn-1-yloxy)phenyl)pyrrolidine-2-carboxamide (9)**

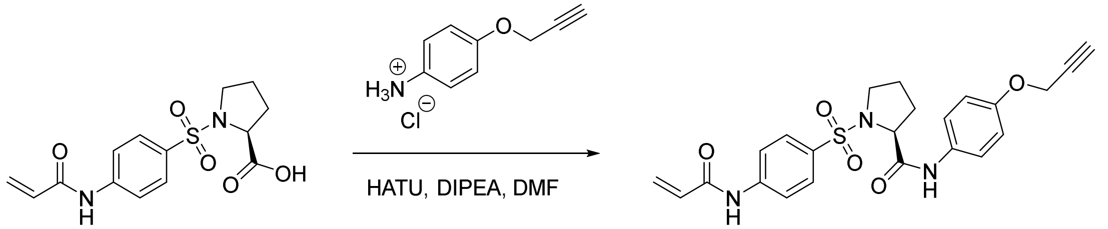


*General procedure E, reagents:* acid **25** (50 mg, 0.15 mmol); aniline **32** (42 mg, 0.23 mmol); HATU (88 mg, 0.23 mmol); DIPEA (140 µL, 0.75 mmol); DMF (800 µL); *purification:* LC (20–80% EtOAc in hexane).
*Yield:* 36 mg, 53%, yellow solid.
**¹H NMR** (400 MHz, Chloroform-*d*) δ: 9.22 (s, 1H), 8.78 (s, 1H), 7.91 – 7.84 (m, 2H), 7.84 – 7.75 (m, 2H), 7.56 – 7.45 (m, 2H), 7.01 – 6.86 (m, 2H), 6.49 (dd, *J* = 16.9, 1.4 Hz, 1H), 6.33 (dd, *J* = 16.9, 10.1 Hz, 1H), 5.81 (dd, *J* = 10.2, 1.4 Hz, 1H), 4.68 (d, *J* = 2.4 Hz, 2H), 4.14 (dd, *J* = 8.9, 3.3 Hz, 1H), 3.63 (ddd, *J* = 10.3, 7.0, 3.2 Hz, 1H), 3.24 (td, *J* = 9.8, 6.4 Hz, 1H), 2.54 (t, *J* = 2.4 Hz, 1H), 1.87 – 1.76 (m, 1H), 1.73 – 1.60 (m, 2H), 1.27 – 1.24 (m, 1H) ppm.
**¹³C NMR** (101 MHz, Chloroform-*d*) δ: 169.5, 164.4, 154.6, 143.4, 131.1, 130.6, 129.6, 129.1, 129.0, 122.0, 121.9, 119.8, 119.8, 115.3, 78.4, 75.6, 62.9, 56.1, 5.14, 30.1, 24.4 ppm.
**LRMS** (+ESI) *m/z*: $[M+H]^+$ 454.
**HRMS** (ESI) *m/z*: $[M+H]^+$ calcd for $C_{23}H_{24}N_3O_5S$: 454.1437; found: 454.1418.
**HPLC** retention time 4.3 min, 97.7%.

**Ethyl (*S*)-4-(1-((4-Acrylamidophenyl)sulfonyl)pyrrolidine-2-carboxamido)benzoate (33)**

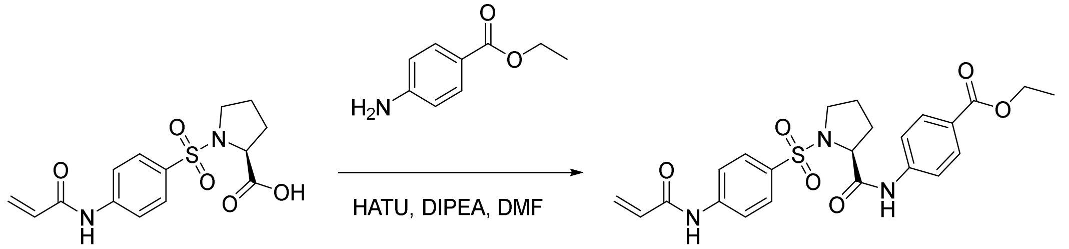


*General procedure E, reagents:* acid **25** (250 mg, 0.77 mmol); ethyl 4-aminobenzoate (130 mg, 0.77 mmol); HATU (350 mg, 0.92 mmol); DIPEA (400 µL, 2.3 mmol); $CH_2Cl_2$ (5.0 mL); *purification:* LC (20–60% EtOAc in hexane).
*Yield:* 250 mg (1:1 mixture with tetramethylurea), 54%, orange foam.
**¹H NMR** (400 MHz, Chloroform-*d*) δ: 9.01 (s, 1H), 8.13 (s, 1H), 8.08 – 8.03 (m, 2H), 7.93 – 7.83 (m, 4H), 7.73 – 7.68 (m, 2H), 6.53 (dd, *J* = 16.8, 1.2 Hz, 1H), 6.33 (dd, *J* = 16.8, 10.2 Hz, 1H), 5.87 (dd, *J* = 10.2, 1.2 Hz, 1H), 4.39 (q, *J* = 7.1 Hz, 2H), 4.18 (dd, *J* = 8.7, 2.7 Hz, 1H), 3.67 (ddd, *J* = 10.3, 7.2, 3.4 Hz, 1H), 3.27 (td, *J* = 9.7, 6.6 Hz, 1H), 2.42 – 2.32 (m, 1H), 1.83 (d, *J* = 18.5 Hz, 3H), 1.72 (ddt, *J* = 12.8, 6.6, 3.3 Hz, 1H), 1.67 – 1.57 (m, 1H), 1.42 (t, *J* = 7.1 Hz, 3H) ppm.

**(*S*)-4-(1-((4-Acrylamidophenyl)sulfonyl)pyrrolidine-2-carboxamido)benzoic acid (34)**

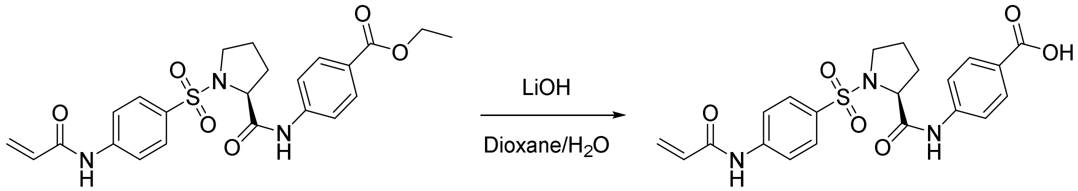


*General procedure D, reagents:* ester **33** (230 mg, 0.48 mmol, 1:1 mixture with tetramethylurea); LiOH.$H_2O$ (30 mg, 0.72 mmol); dioxane (1.0 mL); $H_2O$ (1.0 mL).
*Yield:* 200 mg, 92%, white solid.
**$^1$H NMR** (400 MHz, DMSO-*d*$_6$) δ: 12.77 (s, 1H), 10.60 (s, 1H), 10.28 (s, 1H), 7.98 – 7.89 (m, 4H), 7.89 – 7.83 (m, 2H), 7.78 – 7.72 (m, 2H), 6.47 (dd, *J* = 17.0, 10.0 Hz, 1H), 6.33 (dd, *J* = 17.0, 2.0 Hz, 1H), 5.85 (dd, *J* = 10.0, 2.0 Hz, 1H), 4.21 (t, *J* = 6.1 Hz, 1H), 3.52 – 3.43 (m, 1H), 3.28 – 3.17 (m, 0H), 1.89 (s, 3H), 1.65 – 1.46 (m, 1H) ppm.

**(*S*)-1-((4-Acrylamidophenyl)sulfonyl)-*N*-(4-((1-methyl-1H-pyrazol-3-yl)carbamoyl)phenyl)pyrrolidine-2-carboxamide (10, IMP-2552)**

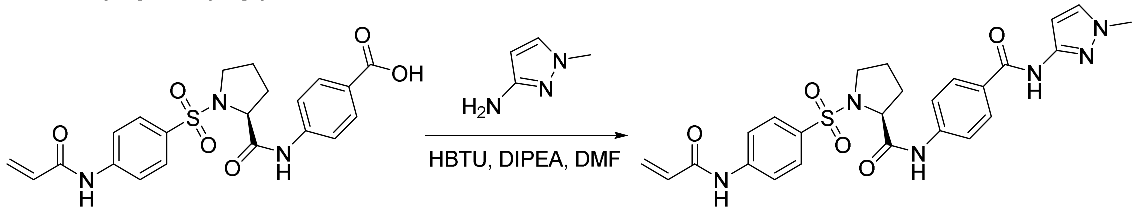


*General procedure E, reagents:* acid **34** (120 mg, 0.26 mmol); 1-methyl-1H-pyrazol-3-amine (26 mg, 0.26 mmol); HBTU (100 mg, 0.26 mmol); DIPEA (120 µL, 0.66 mmol); $CH_2Cl_2$ (5.0 mL); *purification:* LC (0–3% MeOH in $CH_2Cl_2$), then preparative HPLC (20–95% 0.1% formic acid MeCN in $H_2O$).
*Yield:* 15 mg, 11%, white solid.
**$^1$H NMR** (400 MHz, DMSO-*d*$_6$) δ: 10.72 (s, 1H), 10.67 (s, 1H), 10.26 (s, 1H), 8.04 – 7.97 (m, 2H), 7.98 – 7.92 (m, 2H), 7.91 – 7.79 (m, 2H), 7.77 – 7.68 (m, 2H), 7.61 (s, 1H), 6.59 (m, 1H), 6.49 (dd, *J* = 16.9, 10.1 Hz, 1H), 6.33 (d, *J* = 16.9 Hz, 1H), 5.85 (d, *J* = 10.1 Hz, 1H), 4.22 (t, *J* = 6.0 Hz, 1H), 3.79 (s, 3H), 3.49 (dt, *J* = 10.7, 6.0 Hz, 1H), 3.24 (dt, *J* = 10.0, 6.2 Hz, 1H), 1.99 – 1.81 (m, 3H), 1.56 (m, 1H) ppm.
**$^{13}$C NMR** (101 MHz, DMSO-*d*$_6$) δ: 171.0, 164.2, 164.2, 147.6, 143.8, 142.1, 131.9, 131.4, 129.3, 129.2, 129.1, 129.1, 128.6, 119.7, 119.2, 97.9, 62.3, 49.6, 38.8, 31.6, 24.8 ppm.
**LRMS** (+ESI) *m/z*: $[M+H]^+$ 523.
**HRMS** (+ESI) *m/z*: $[M+H]^+$ calcd for $C_{25}H_{27}N_6O_5S$ 523.1758, found 523.1778.
**HPLC** retention time 3.6 min, >99.0%.

***tert*-Butyl (4-((1-methyl-1H-pyrazol-3-yl)carbamoyl)phenyl)carbamate (35)**

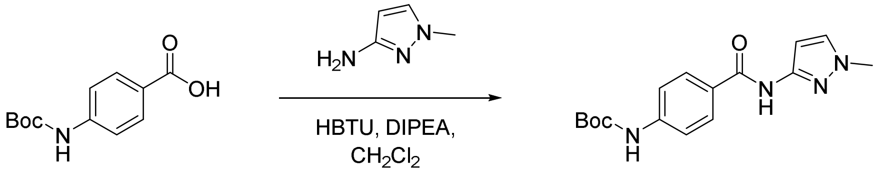


*General procedure E, reagents:* 4-Boc-aminobenzoic acid (551 mg, 2.3 mmol); 1-methyl-1*H*-pyrazol-3-amine (205 mg, 2.1 mmol); HBTU (961 mg, 2.5 mmol); DIPEA (920 µL, 5.3 mmol); DMF (4 mL); *purification:* LC (10–90% EtOAc in hexane).

*Yield:* 610 mg, 91%, white solid.
**$^1$H NMR** (400 MHz, Methanol-$d_4$) δ: 7.90 – 7.86 (m, 2H), 7.59 – 7.54 (m, 2H), 7.52 (d, *J* = 2.3 Hz, 1H), 6.61 (d, *J* = 2.3 Hz, 1H), 3.85 (s, 3H), 1.55 (s, 9H) ppm.
**LRMS** (+ESI) *m/z*: $[M+H]^+$ 317.

**4-Amino-*N*-(1-methyl-1H-pyrazol-3-yl)benzamide hydrogen chloride (36)**

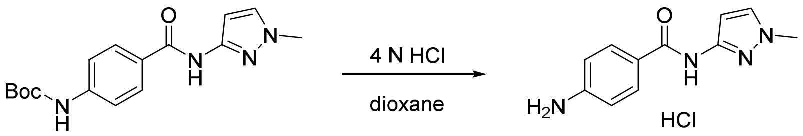


A solution of carbamate **35** (300 mg, 1.0 mmol) in 4 N HCl in dioxane (2.5 mL) was stirred at rt for 23 h. To the crude suspension was added $CH_2Cl_2$ and filtered *in vacuo*. The precipitate washed with $CH_2Cl_2$ (20 mL) and hexane (10 mL) to afford the desired product which was used without further purification.
*Yield:* 240 mg, 99%, white solid.
**$^1$H NMR** (400 MHz, DMSO-$d_6$) δ: 10.88 (s, 1H), 8.03 (d, *J* = 8.2 Hz, 1H), 7.38 (d, *J* = 8.2 Hz, 1H), 6.58 (d, *J* = 2.3 Hz, 1H), 4.16 (s, 2H), 3.79 (s, 3H) ppm.
**LRMS** (+ESI) *m/z*: $[M+H]^+$ 217.

**(*R*)-1-((4-Acrylamidophenyl)sulfonyl)-*N*-(4-((1-methyl-1H-pyrazol-3-yl)carbamoyl)phenyl)pyrrolidine-2-carboxamide (11)**

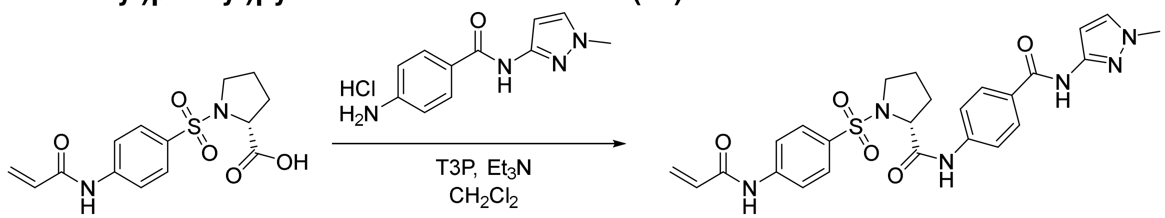


*General procedure E, reagents:* acid **35** (50 mg, 0.15 mmol); aniline **X** (40 mg, 0.19 mmol); $T_3P$ (50% in EtOAc, 196 µL, 0.31 mmol); $Et_3N$ (86 µL, 0.62 mmol); $CH_2Cl_2$ (2.0 mL); *purification:* LC (0–10% MeOH in $CH_2Cl_2$).
*Yield:* 19 mg, 24%, off-white solid.
**$^1$H NMR** (400 MHz, DMSO-$d_6$) δ: 10.71 (s, 1H), 10.59 (s, 1H), 10.21 (s, 1H), 8.03 – 7.96 (m, 2H), 7.95 – 7.89 (m, 2H), 7.88 – 7.82 (m, 2H), 7.74 – 7.69 (m, 2H), 7.60 (d, J = 2.1 Hz, 1H), 6.58 (d, J = 2.1 Hz, 1H), 6.46 (dd, J = 16.9, 10.0 Hz, 1H), 6.33 (dd, J = 16.9, 1.8 Hz, 1H), 5.84 (dd, J = 10.0, 1.8 Hz, 1H), 4.23 – 4.17 (m, 1H), 3.78 (s, 3H), 3.53 – 3.44 (m, 1H), 3.27 – 3.18 (m, 1H), 1.94 – 1.82 (m, 3H), 1.60 – 1.49 (m, 1H) ppm.
**LRMS** (+ESI) *m/z*: $[M+H]^+$ 523.
**HRMS** (+ESI) *m/z*: $[M+H]^+$ calcd for $C_{25}H_{27}N_6O_5S$ 523.1758, found 523.1768.
**HPLC** retention time 4.09 min, 94.4%.

**4-Nitro-1-(prop-2-yn-1-yl)-1H-pyrazole (37)**

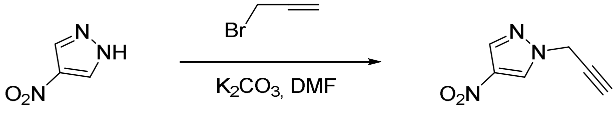

To a transparent stirring solution of 4-nitropyrazole (2.8 g, 18 mmol) in DMF (2.0 mL), were added $K_2CO_3$ (6.1 g, 28 mmol) and propargyl bromide (80% in toluene, 1.4 mL, 28 mmol). The reaction mixture immediately turned brown, and was stirred at rt for 1 h, before quenching with $H_2O$ (50 mL). The aqueous phase was extracted with EtOAc (10 mL, 3x) and the combined organic phase was washed with brine (10 mL), dried ($MgSO_4$), filtered and concentrated *in vacuo*. The crude mixture was purified by liquid chromatography (0–20% EtOAc in hexane) to afford the desired compound as a pale-yellow oil (2.3 g, 88%).

**$^1$H NMR** (400 MHz, Chloroform-*d*) δ: 8.44 (s, 1H), 8.12 (s, 1H), 5.01 (d, *J* = 2.6 Hz, 2H), 2.70 (t, *J* = 2.6 Hz, 1H) ppm.

#### 1-(Prop-2-yn-1-yl)-1H-pyrazol-4-amine (38)

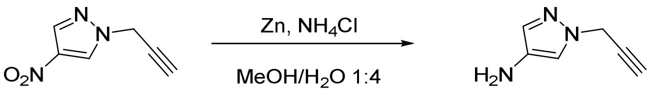


To a clear stirring solution of pyrazole **37** (500 mg, 3.3 mmol) in MeOH (10 mL), were added $NH_4Cl$ (1.8 g, 33 mmol) and zinc (1.1 g, 17 mmol). The reaction mixture slowly turned to dark orange, and after 30 min $H_2O$ (1.0 mL) was added. After 4 h, $NH_4Cl$ (1.8 g, 33 mmol) in $H_2O$ (1.0 mL) and zinc (1.1 g, 17 mmol) were added to the solution. After 16 h, the reaction mixture was filtered through celite and washed with MeOH, then concentrated *in vacuo*. The crude mixture was purified by liquid chromatography (50–100% EtOAc in hexane) to afford the desired compound as a brown oil (60 mg, 15%) which was used immediately in the next step.

**$^1$H NMR** (400 MHz, DMSO-*d*$_6$) δ: 7.07 (s, 1H), 6.94 (s, 1H), 4.82 (d, *J* = 2.5 Hz, 2H), 3.89 (broad s, 2H), 3.41 (t, *J* = 2.6 Hz, 1H) ppm.

#### (*S*)-1-((4-Acrylamidophenyl)sulfonyl)-*N*-(4-((1-(prop-2-yn-1-yl)-1H-pyrazol-4-yl)carbamoyl)phenyl)pyrrolidine-2-carboxamide (12, IMP-2566)

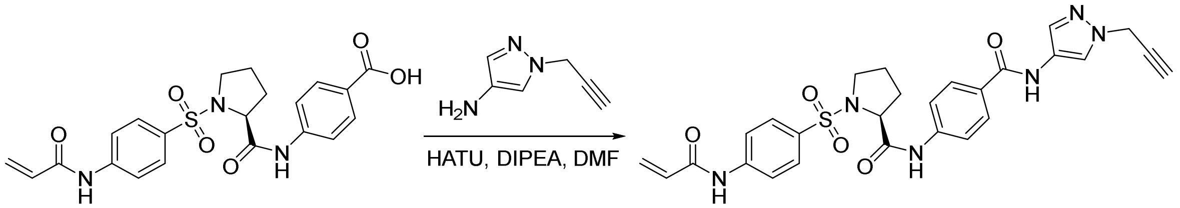


*General procedure E, reagents:* acid **34** (95 mg, 0.21 mmol); aminopyrazole **X** (52 mg, 0.21 mmol); HATU (81 mg, 0.21 mmol); DIPEA (150 µL, 0.85 mmol); DMF (1.0 mL); *purification:* preparative HPLC (30–95% 0.1% formic acid MeCN in $H_2O$).
*Yield:* 60 mg, 51%, pale pink solid.
**$^1$H NMR** (400 MHz, DMSO-*d*$_6$) δ: 10.71 (s, 1H), 10.44 (s, 1H), 10.31 (s, 1H), 8.16 (s, 1H), 8.00 – 7.91 (m, 4H), 7.90 – 7.83 (m, 2H), 7.81 – 7.74 (m, 2H), 7.64 (s, 1H), 6.49 (dd, *J* = 16.9, 10.1 Hz, 1H), 6.33 (dd, *J* = 17.0, 1.9 Hz, 1H), 5.85 (dd, *J* = 9.9, 2.0 Hz, 1H), 5.04 (d, *J* = 2.6 Hz, 2H), 4.23 (t, *J* = 6.0 Hz, 1H), 3.56 – 3.47 (m, 2H), 3.24 (dt, *J* = 10.8, 6.6 Hz, 1H), 1.96 – 1.81 (m, 3H), 1.55 (p, *J* = 7.4, 6.4 Hz, 1H) ppm.

**$^{13}$C NMR** (101 MHz, DMSO-*d*$_6$) δ: 170.96, 164.20, 163.38, 143.73, 142.02, 131.86, 131.46, 131.34, 129.20, 129.06, 128.73, 128.58, 122.75, 121.01, 119.64, 119.28, 79.15, 76.71, 62.33, 49.63, 41.45, 31.53, 24.81 ppm.
**LRMS** (+ESI) *m/z*: $[M+H]^+$ 547.
**HRMS** (+ESI) *m/z*: $[M+H]^+$ calcd for $C_{27}H_{27}N_6O_5S$ 547.1764, found 547.1757.
**HPLC** retention time 3.6 min, >99.0%.

**(*S*)-1-((4-Acrylamidophenyl)sulfonyl)-*N*-(3-methoxyphenyl)pyrrolidine-2-carboxamide (13, IMP-2505)**

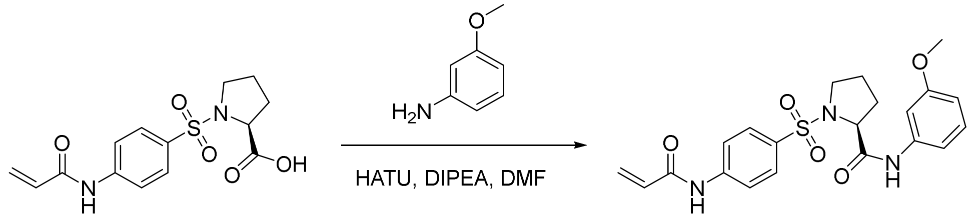


*General procedure E, reagents:* acid **25** (30 mg, 0.09 mmol); 3-methoxyaniline (17 mg, 0.13 mmol); HATU (51 mg, 0.13 mmol); DIPEA (63 µL, 0.36 mmol); DMF (1.0 mL); *purification:* reverse phase LC (5–100% 0.1% formic acid MeCN in $H_2O$).
*Yield:* 26 mg, 67%, off-white solid.
**$^1$H NMR** (400 MHz, DMSO-*d*$_6$) δ: 10.61 (s, 1H), 9.95 (s, 1H), 7.95 – 7.88 (m, 2H), 7.87 – 7.83 (m, 2H), 7.33 (t, *J* = 2.2 Hz, 1H), 7.23 (t, *J* = 8.1 Hz, 1H), 7.17 (ddd, *J* = 8.1, 1.9, 1.1 Hz, 1H), 6.66 (ddd, *J* = 8.2, 2.6, 1.1 Hz, 1H), 6.47 (dd, *J* = 17.0, 10.0 Hz, 1H), 6.33 (dd, *J* = 16.9, 2.0 Hz, 1H), 5.88 – 5.81 (m, 1H), 4.18 (t, *J* = 6.1 Hz, 1H), 3.74 (s, 3H), 3.50 – 3.43 (m, 1H), 3.23 (dt, *J* = 10.1, 6.0 Hz, 1H), 1.87 (ddd, *J* = 5.8, 4.2, 1.9 Hz, 2H), 1.58 – 1.47 (m, 1H), 1.19 (d, *J* = 6.6 Hz, 1H) ppm.
**$^{13}$C NMR** (101 MHz, DMSO-*d*$_6$) δ: 170.5, 164.1, 143.6, 140.3, 131.8, 129.9, 129.0, 128.5, 119.6, 112.1, 109.6, 105.5, 62.3, 55.4, 49.6, 31.5, 24.7 ppm.
**LRMS** (+ESI) *m/z*: $[M+H]^+$ 431.
**HRMS** (+ESI) *m/z*: $[M+H]^+$ calcd for $C_{21}H_{24}N_3O_5S$ 430.1431, found 430.1431.
**HPLC** retention time 3.9 min, >99.0%.

**1-((2-methoxy-4-nitrophenyl)sulfonyl)pyrrolidine (39)**

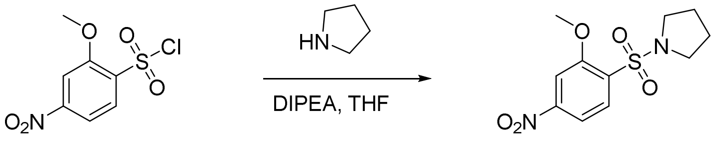


*General procedure A, reagents:* pyrrolidine (130 µL, 1.6 mmol); 2-methoxy-4-nitrophenylsulfonyl chloride (350 mg, 1.6 mmol); DIPEA (550 µL, 3.2 mmol); THF (4.0 mL).
*Yield:* 390 mg, 85%, orange solid.
**$^1$H NMR** (400 MHz, Chloroform-*d*) δ: 8.15 (d, *J* = 8.5 Hz, 1H), 7.92 (dd, *J* = 8.5, 2.1 Hz, 1H), 7.87 (d, *J* = 2.1 Hz, 1H), 4.08 (s, 3H), 3.51 – 3.36 (m, 4H), 1.98 – 1.81 (m, 4H) ppm.
**$^{13}$C NMR** (101 MHz, Chloroform-*d*) δ: 157.3, 151.2, 133.3, 132.8, 115.2, 107.3, 56.8, 47.9, 25.9 ppm.

**3-methoxy-4-(pyrrolidin-1-ylsulfonyl)aniline (40)**

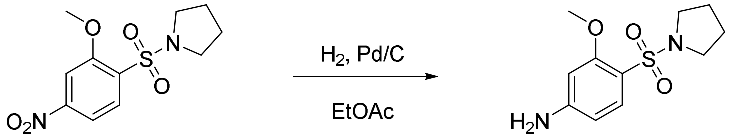

*General procedure B, reagents:* nitrosulfonamide **39** (380 mg, 1.3 mmol); $H_2$ (1 atm); 10% Pd/C (38 mg, 10% w/w); EtOAc (20 mL).
*Yield:* 340 mg, quantitative, brown solid.
**$^1$H NMR** (400 MHz, Chloroform-*d*) δ: 7.36 (d, *J* = 8.6 Hz, 1H), 6.25 (d, *J* = 2.0 Hz, 1H), 6.15 (dd, *J* = 8.6, 2.0 Hz, 1H), 5.99 (s, 2H), 3.16 – 3.10 (m, 4H), 1.72 – 1.63 (m, 4H) ppm.
**$^{13}$C NMR** (101 MHz, Chloroform-*d*) δ: 163.4, 160.0, 138.2, 116.6, 109.9, 101.6, 60.4, 52.6, 30.4 ppm.

**N-(3-methoxy-4-(pyrrolidin-1-ylsulfonyl)phenyl)acrylamide (14, IMP1721)**

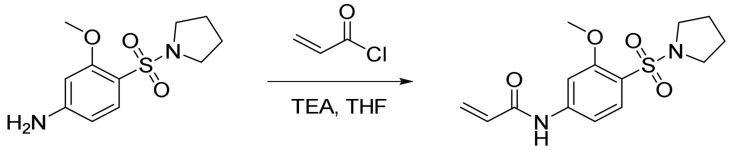

*General procedure C, reagents:* aniline **40** (100 mg, 0.39 mmol); acryloyl chloride (50 µL, 0.59 mmol); $Et_3N$ (90 µL, 0.65 mmol); THF (2.0 mL); *purification:* LC (20–80% EtOAc in hexane).
*Yield:* 81 mg, 67%, yellow vetrous solid.
**$^1$H NMR** (400 MHz, DMSO-*d*$_6$) δ: 10.53 (s, 1H), 7.71 (d, *J* = 8.6 Hz, 1H), 7.67 (d, *J* = 1.9 Hz, 1H), 7.31 (dd, *J* = 8.6, 1.9 Hz, 1H), 6.45 (dd, *J* = 17.0, 10.0 Hz, 1H), 6.32 (dd, *J* = 17.0, 2.0 Hz, 1H), 5.84 (dd, *J* = 10.0, 2.0 Hz, 1H), 3.87 (s, 3H), 3.27 – 3.15 (m, 4H), 1.80 – 1.67 (m, 4H) ppm.
**$^{13}$C NMR** (101 MHz, DMSO-*d*$_6$) δ: 164.2, 157.5, 145.0, 132.7, 131.9, 128.6, 120.8, 110.8, 103.4, 56.2, 48.0, 25.67 ppm.
**HRMS** (ESI) *m/z*: $[M+H]^+$ calcd for $C_{14}H_{19}N_2O_4S$: 311.1066; found: 311.1052.

### NMR Figures

#### *N*-(4-(Pyrrolidin-1-ylsulfonyl)phenyl)acrylamide (1, IMP-1704)

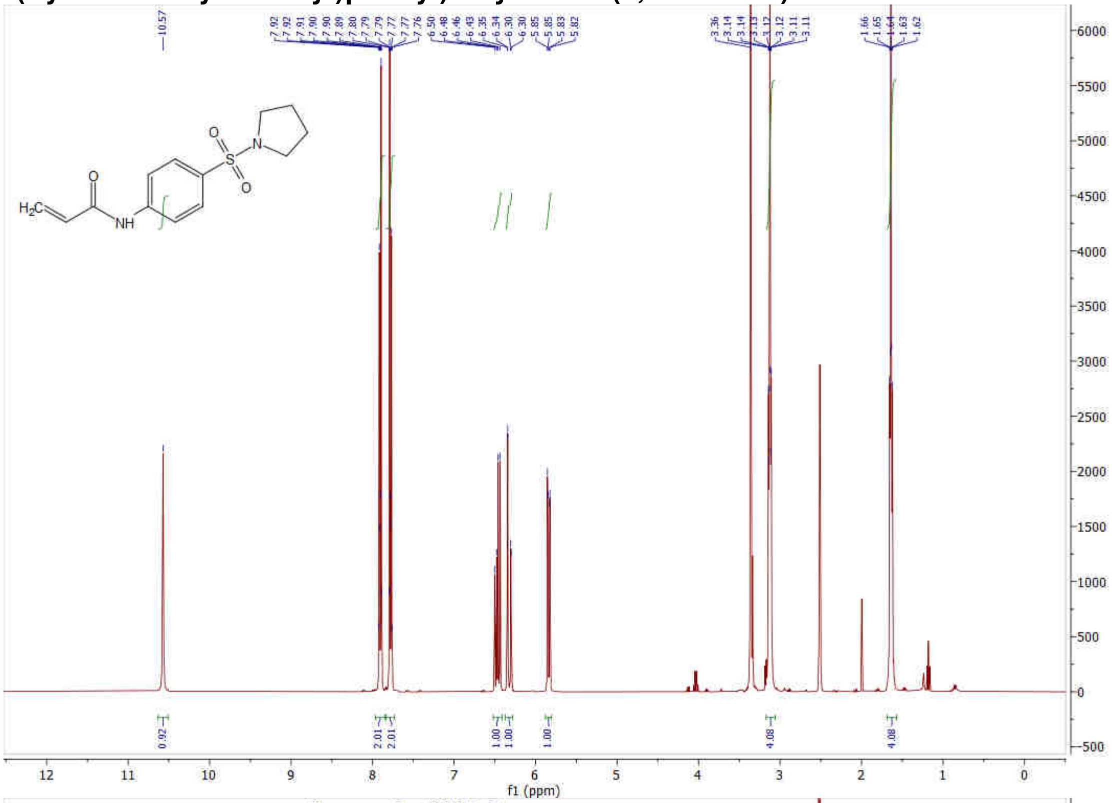


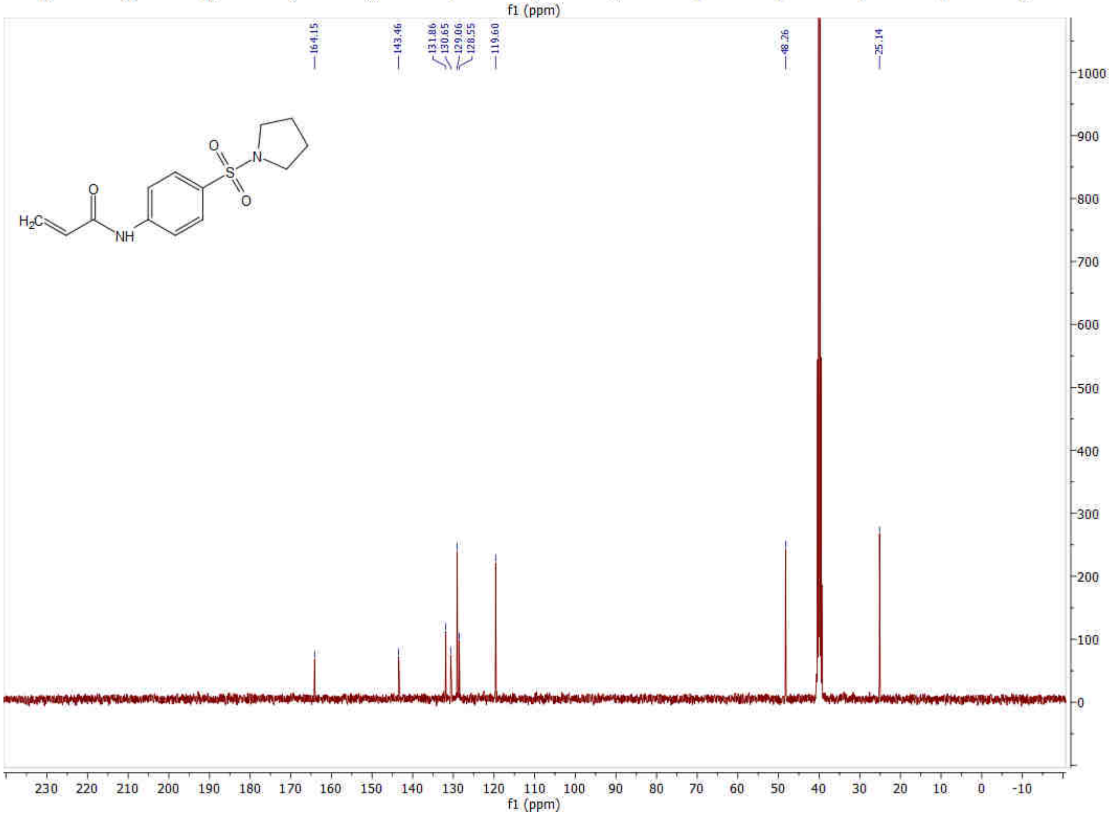

**N-(4-(Pyrrolidine-1-carbonyl)phenyl)acrylamide (2)**

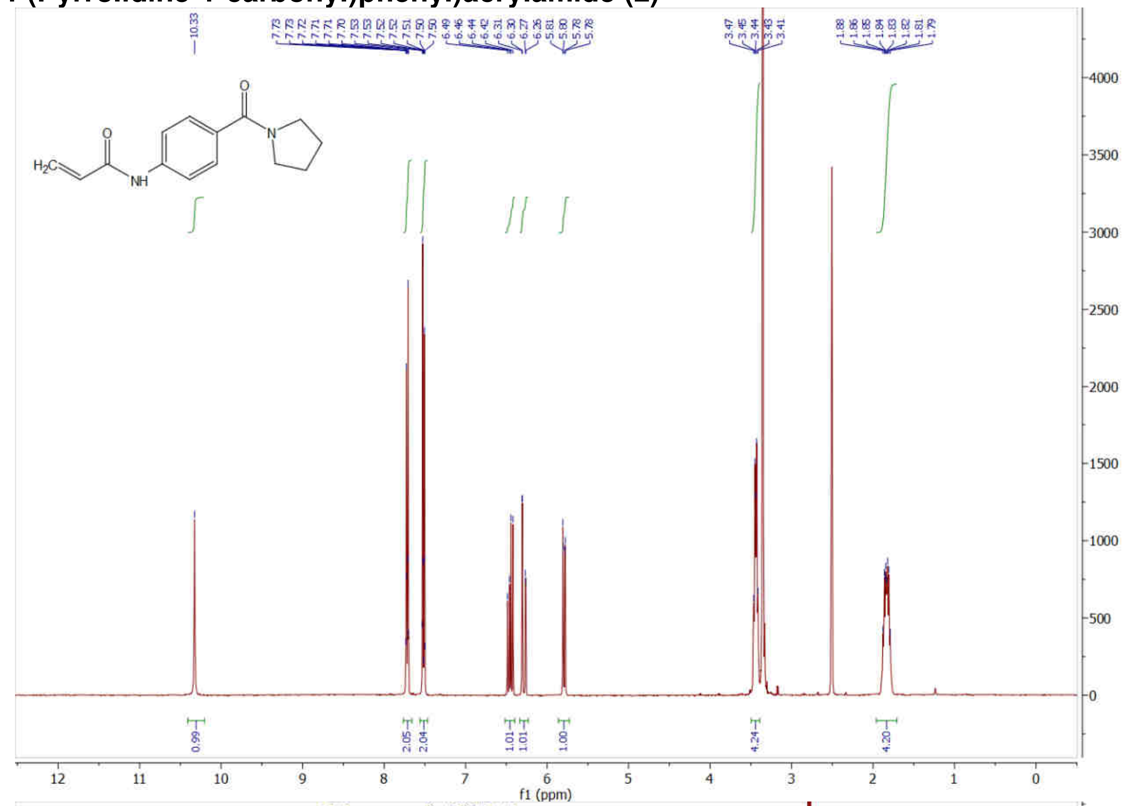


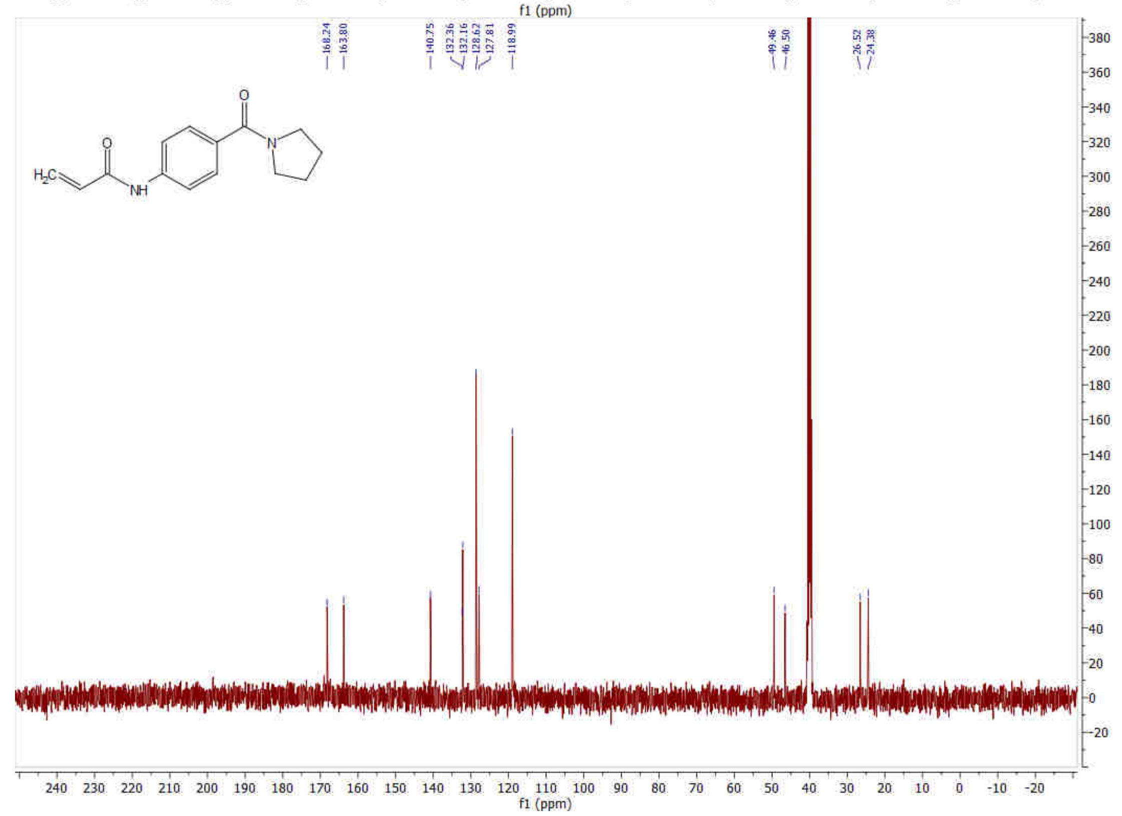

***N*-(3-(Pyrrolidin-1-ylsulfonyl)phenyl)acrylamide (3)**

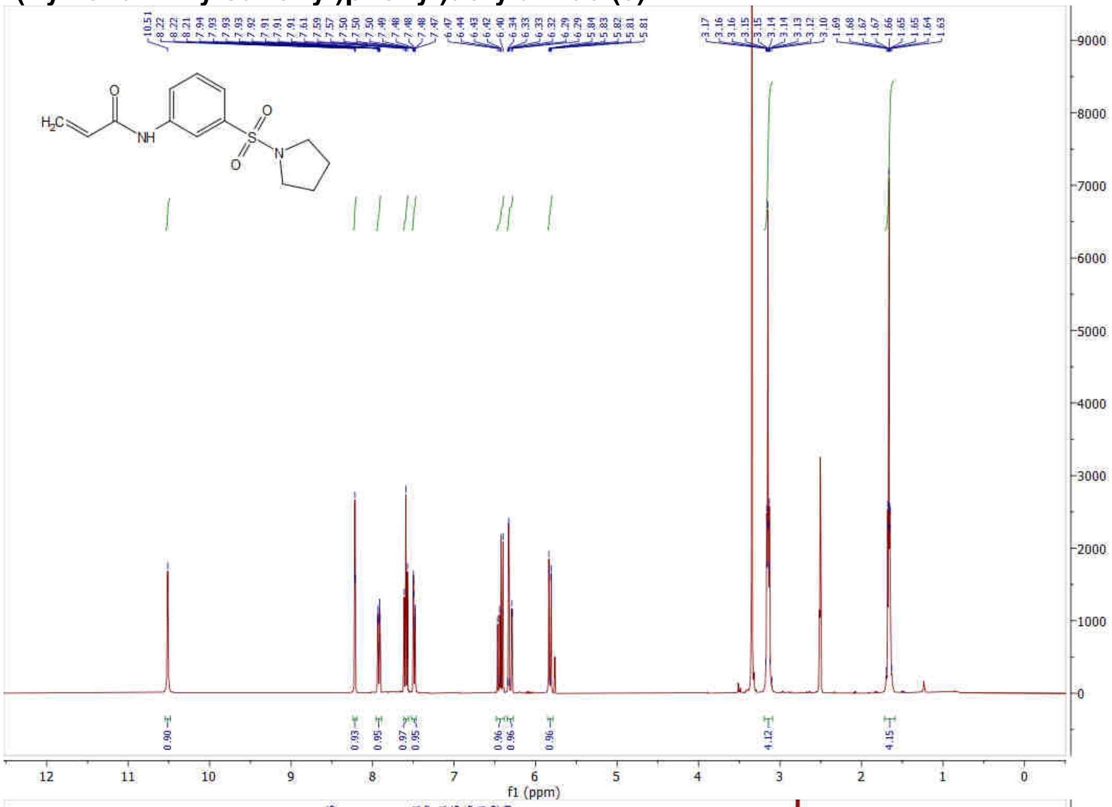


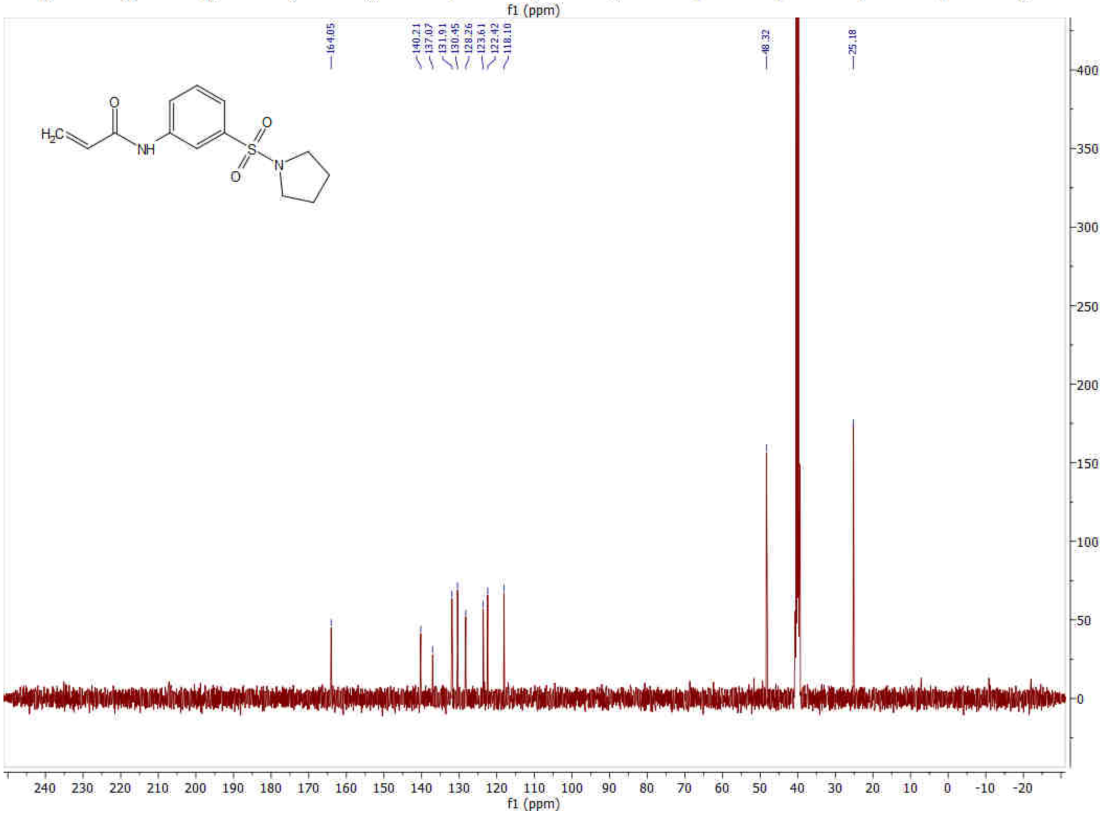

**Methyl ((4-acrylamidophenyl)sulfonyl)-*L*-prolinate (4, IMP1712)**

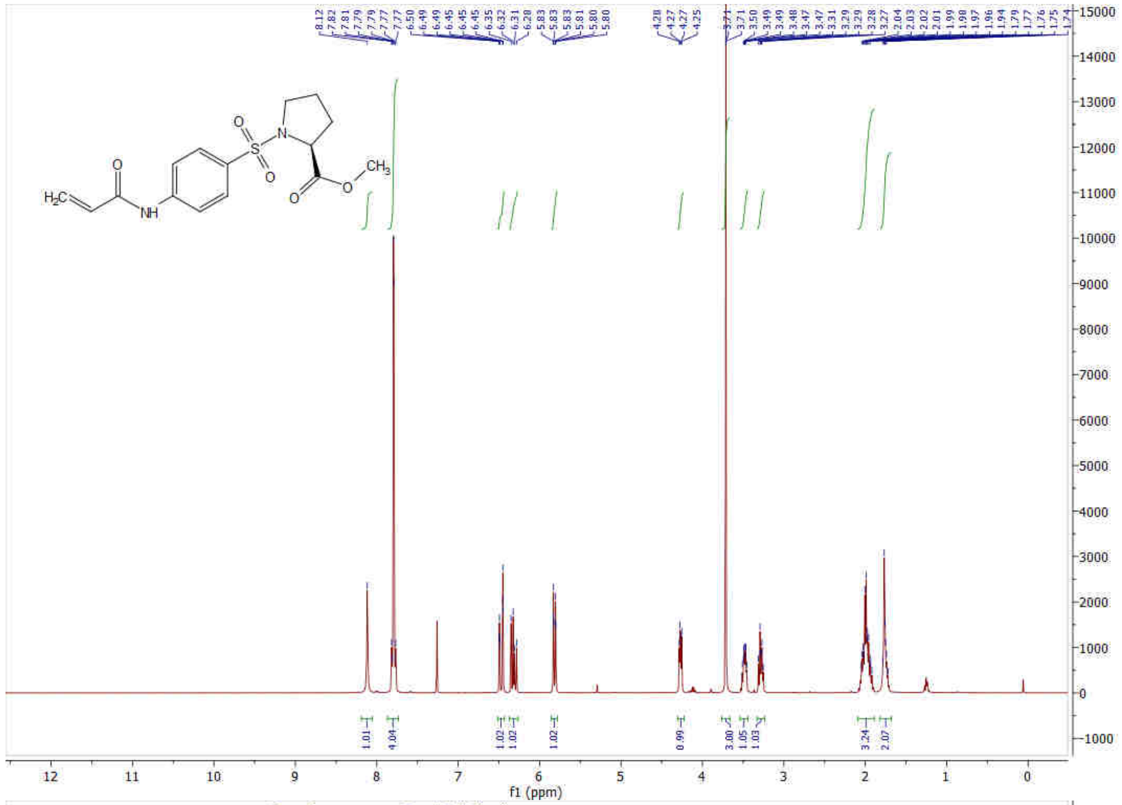

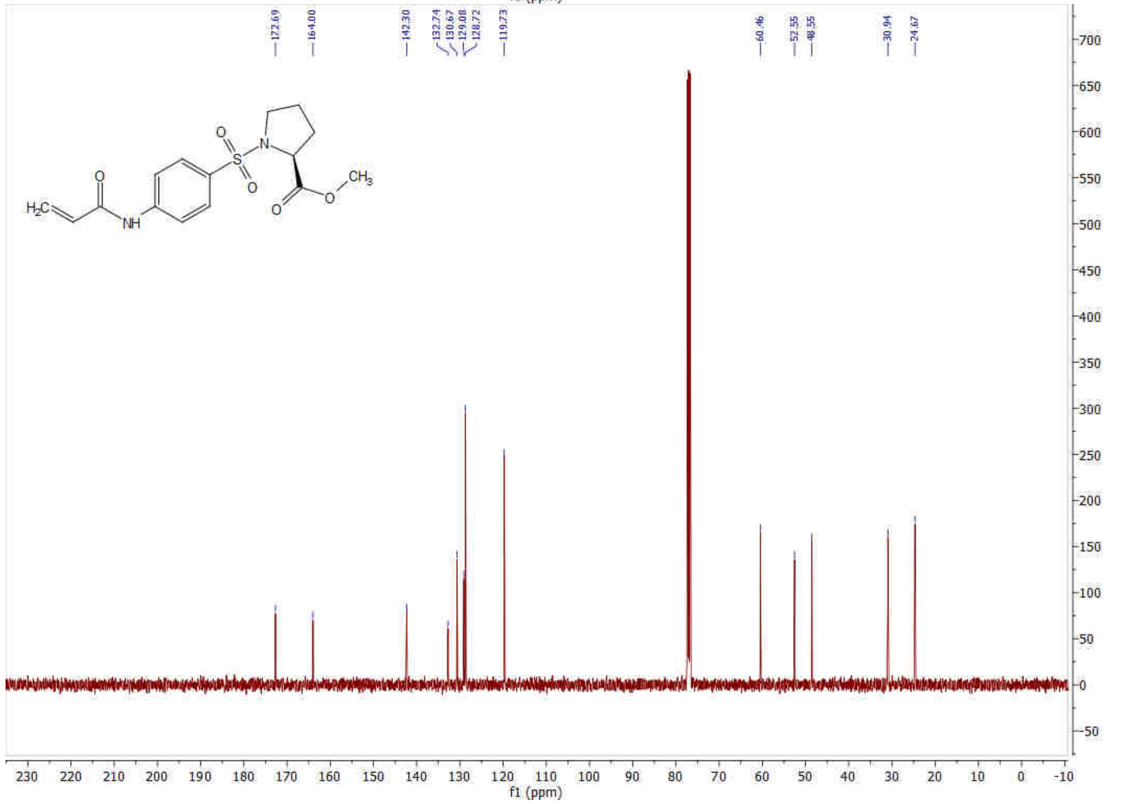

### Methyl ((4-acrylamidophenyl)sulfonyl)-*D*-prolinate (5)

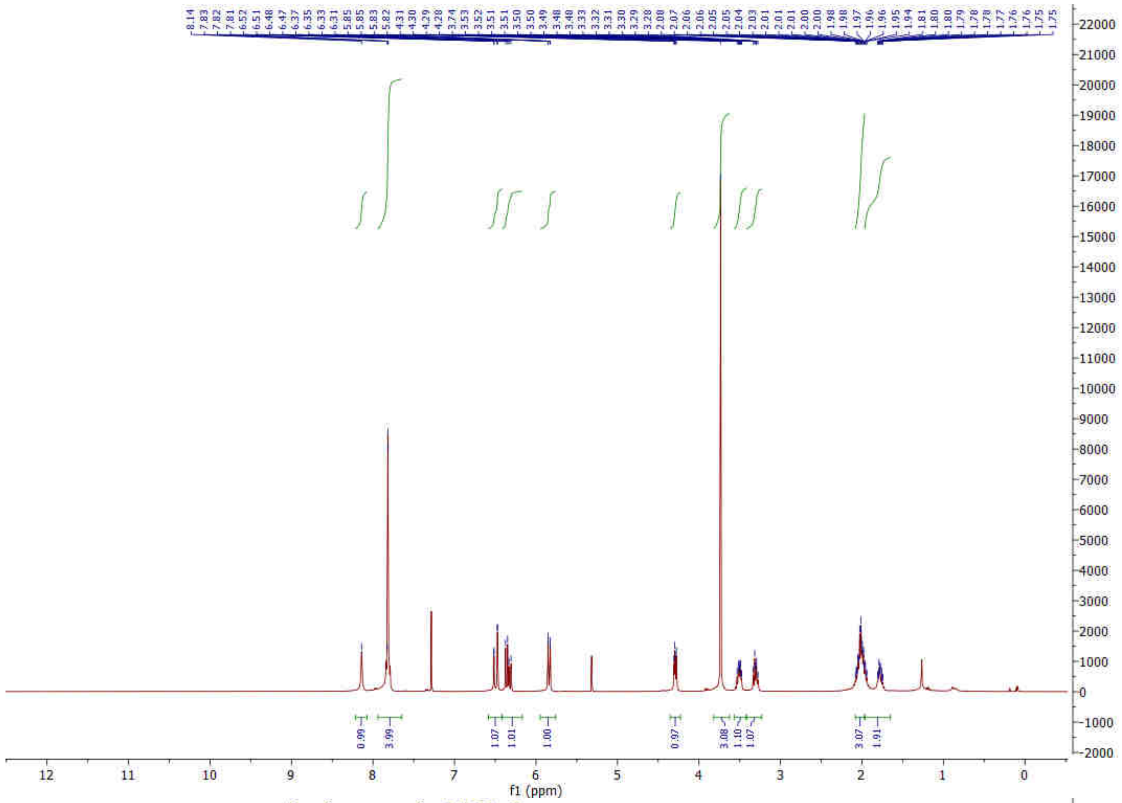


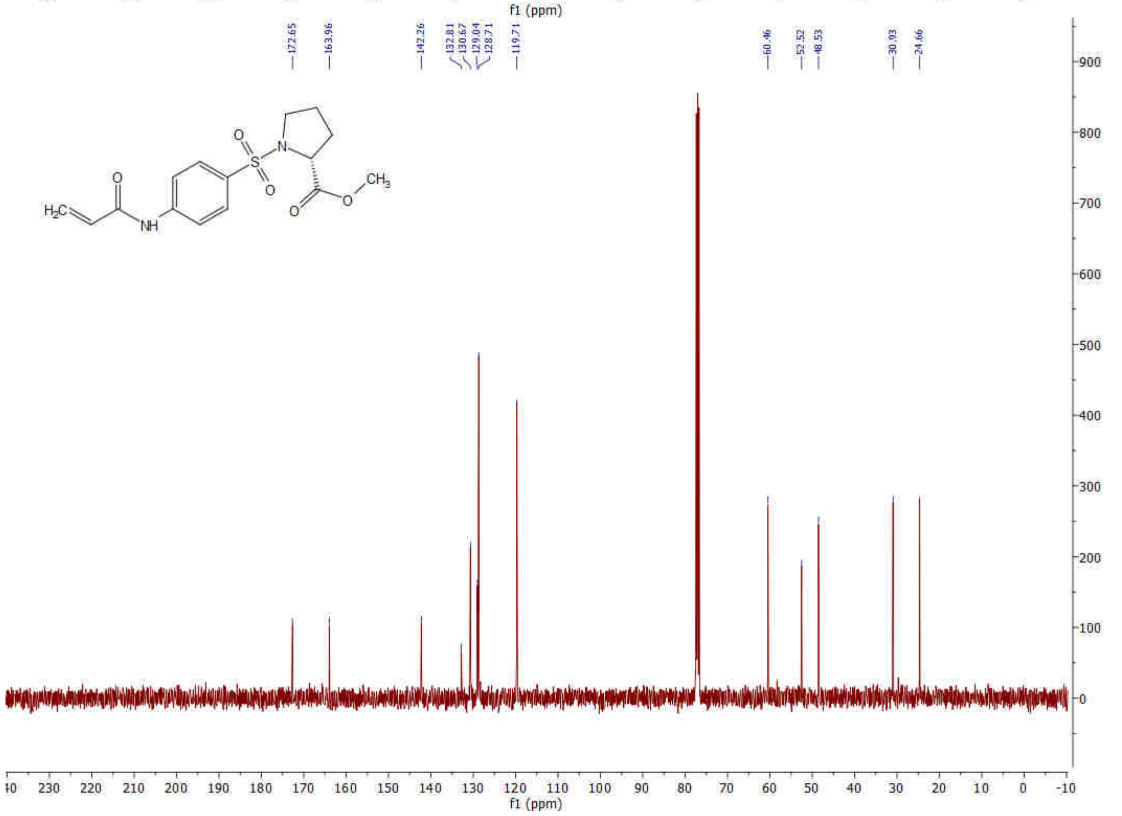

**(*S*)-1-((4-Acrylamidophenyl)sulfonyl)-*N*-(prop-2-yn-1-yl)pyrrolidine-2-carboxamide (6)**

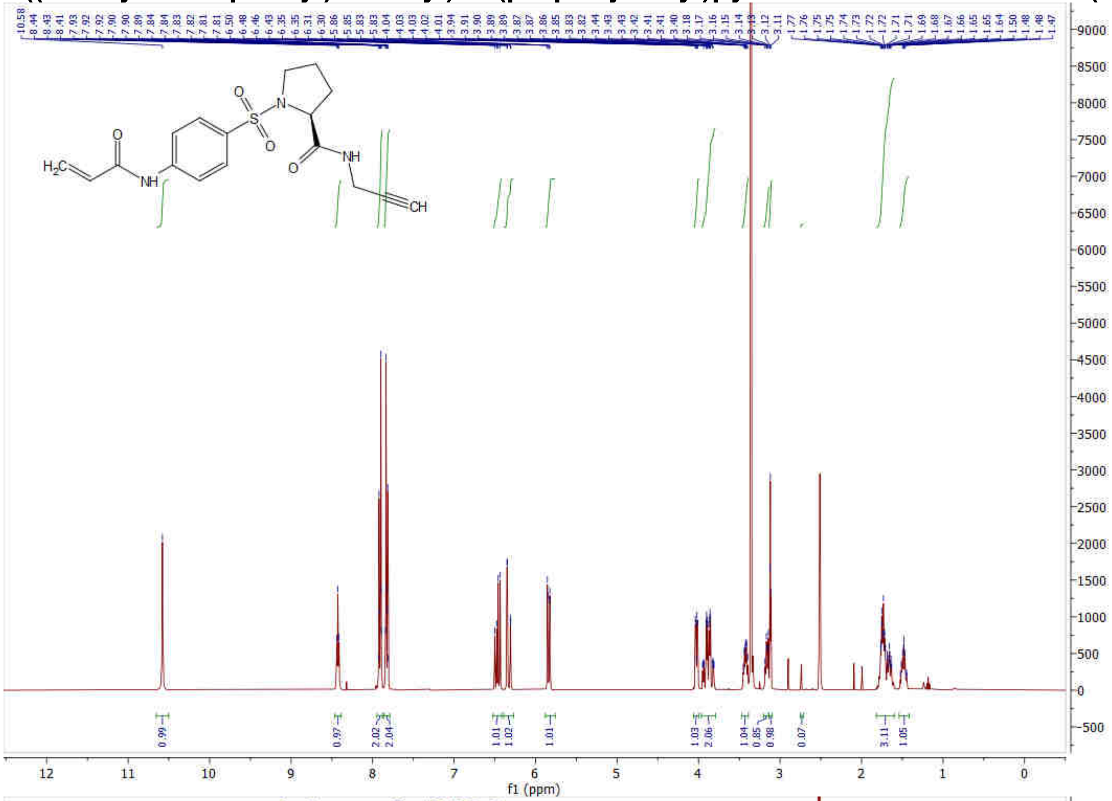

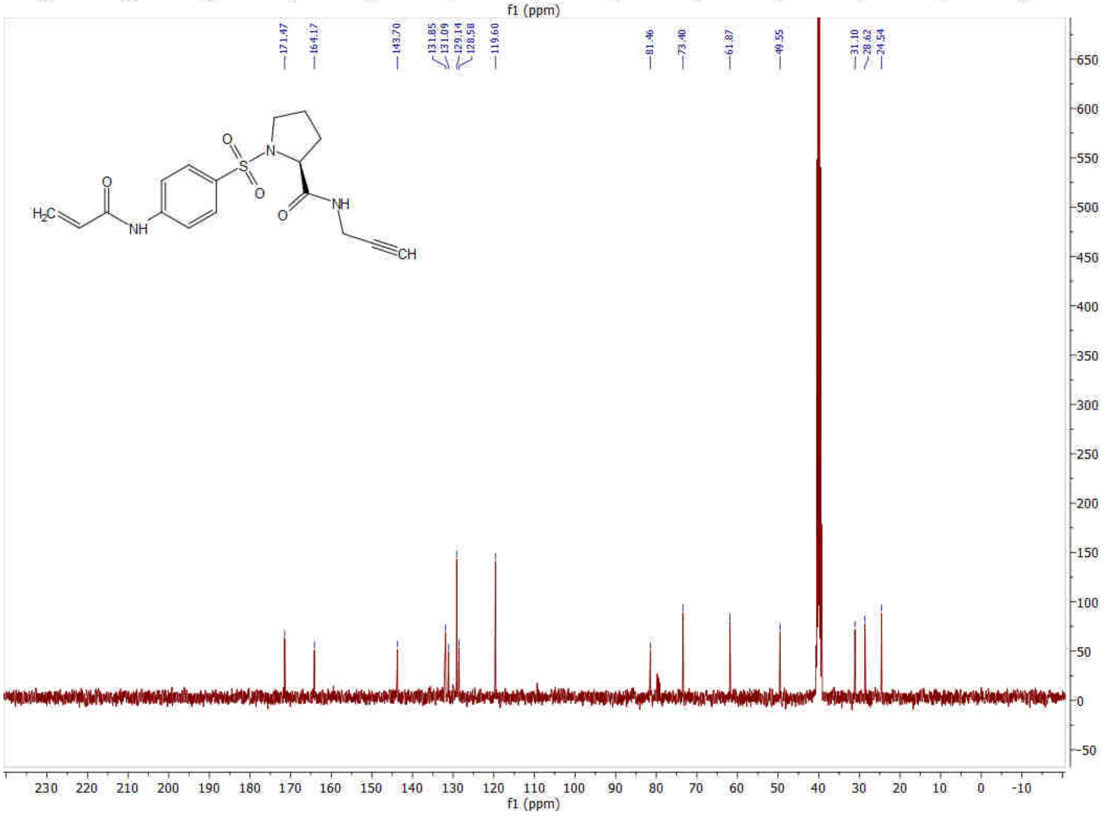

**(*S*)-1-((4-Acrylamidophenyl)sulfonyl)-*N*-(4-methoxyphenyl)pyrrolidine-2-carboxamide (7)**

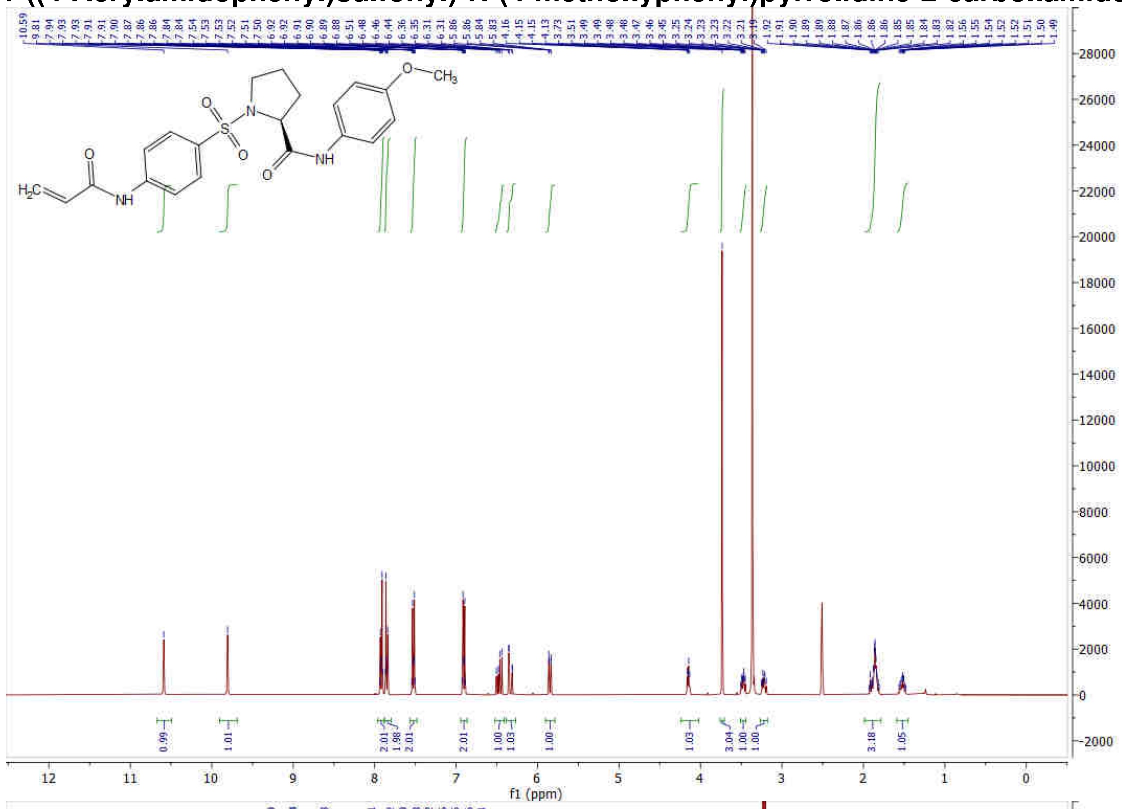


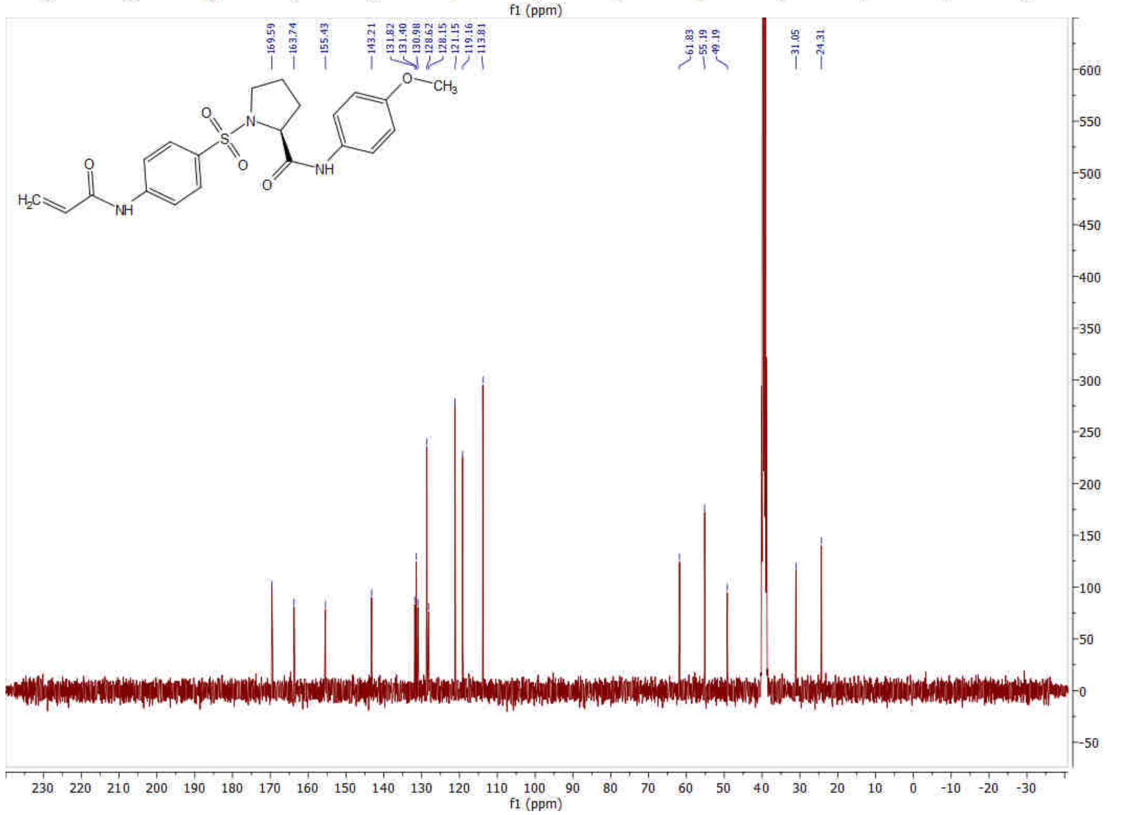

**(*R*)-1-((4-Acrylamidophenyl)sulfonyl)-*N*-(4-methoxyphenyl)pyrrolidine-2-carboxamide (8)**

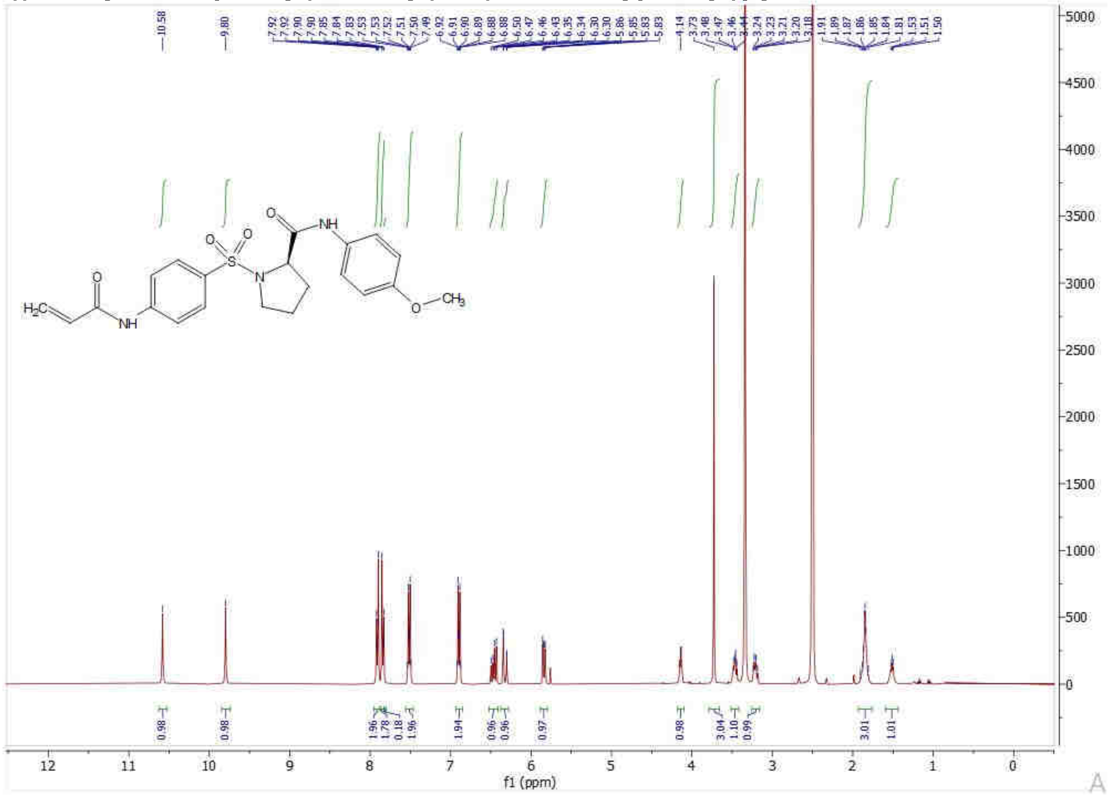

**(*S*)-1-((4-Acrylamidophenyl)sulfonyl)-*N*-(4-(prop-2-yn-1-yloxy)phenyl)pyrrolidine-2-carboxamide (9)**

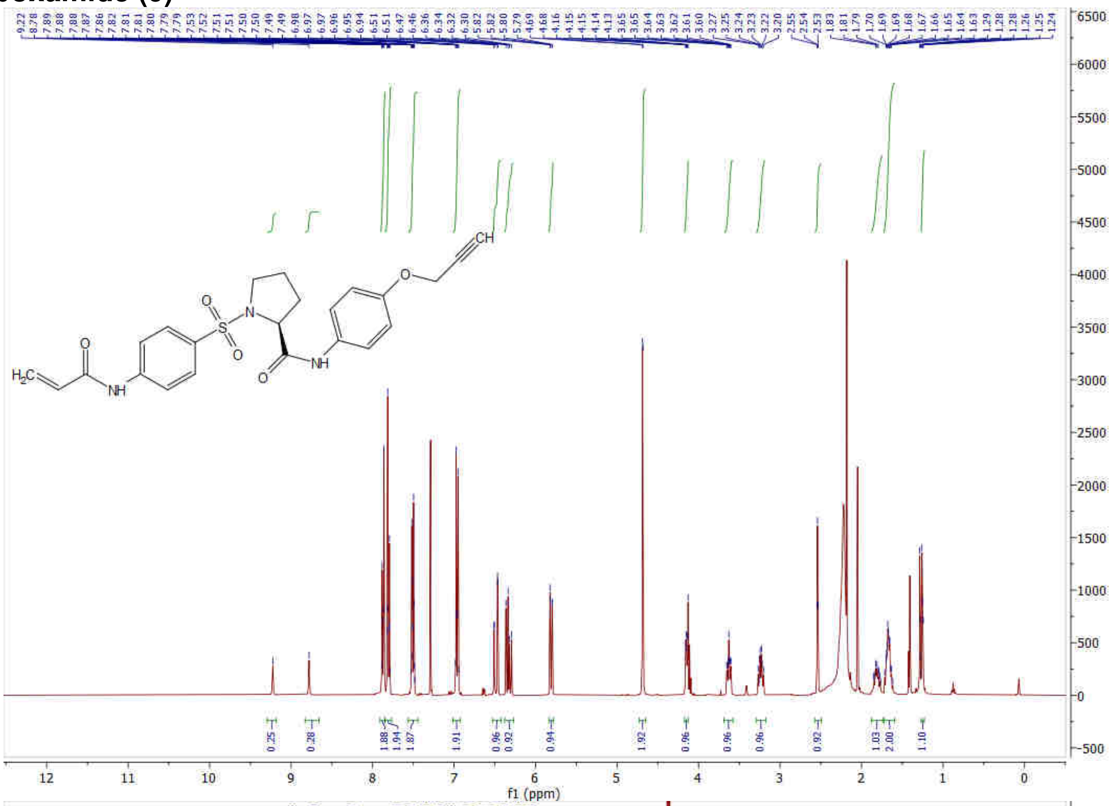


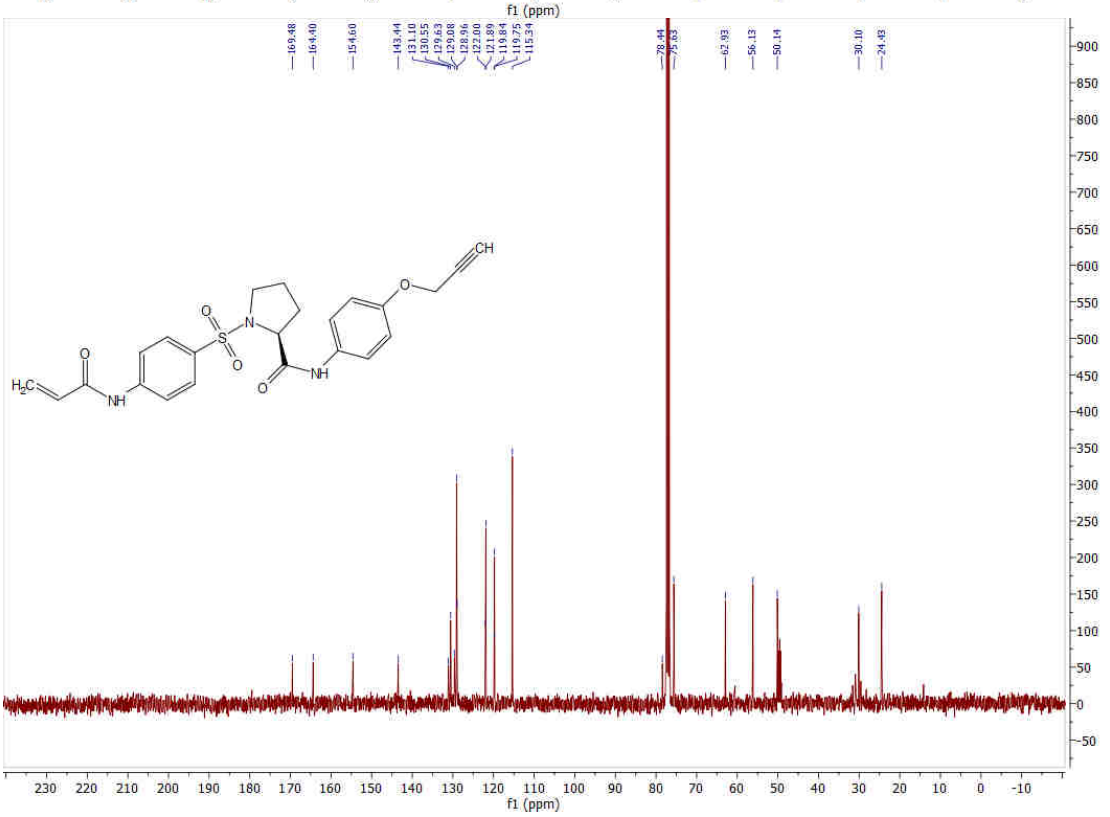

**(*S*)-1-((4-Acrylamidophenyl)sulfonyl)-*N*-(4-((1-methyl-1H-pyrazol-3-yl)carbamoyl)phenyl)pyrrolidine-2-carboxamide (10, IMP-2552)**

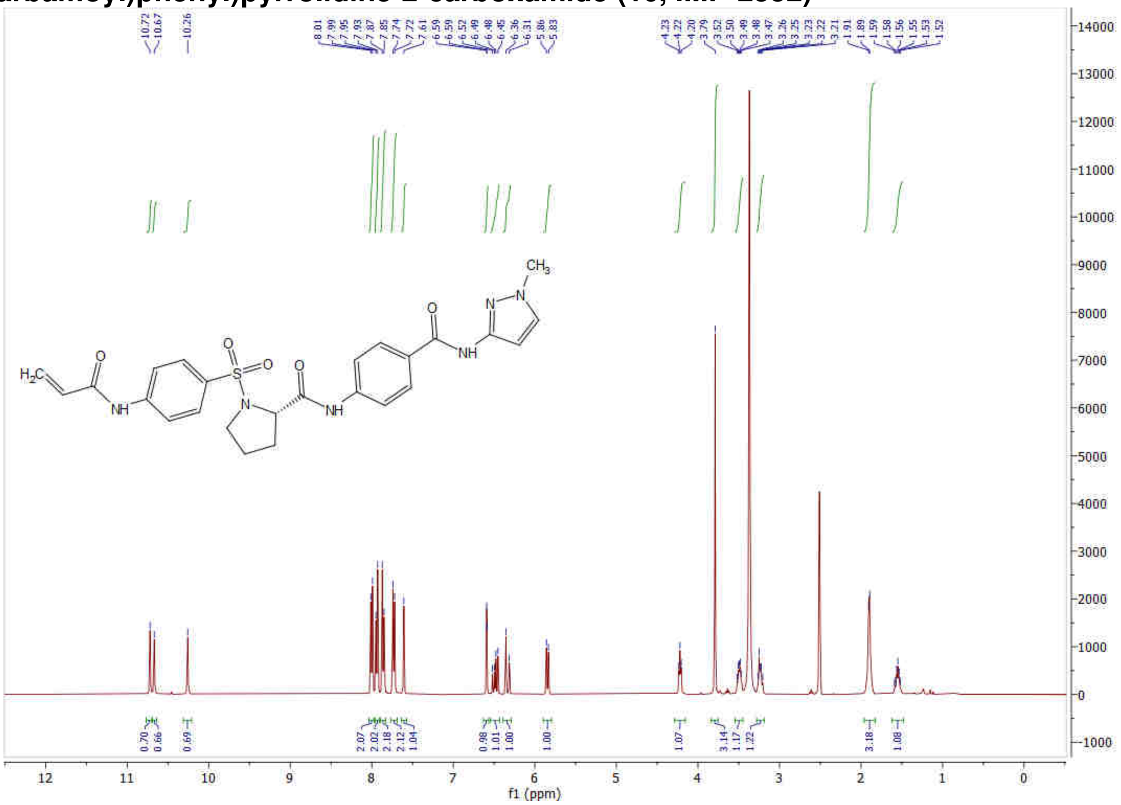


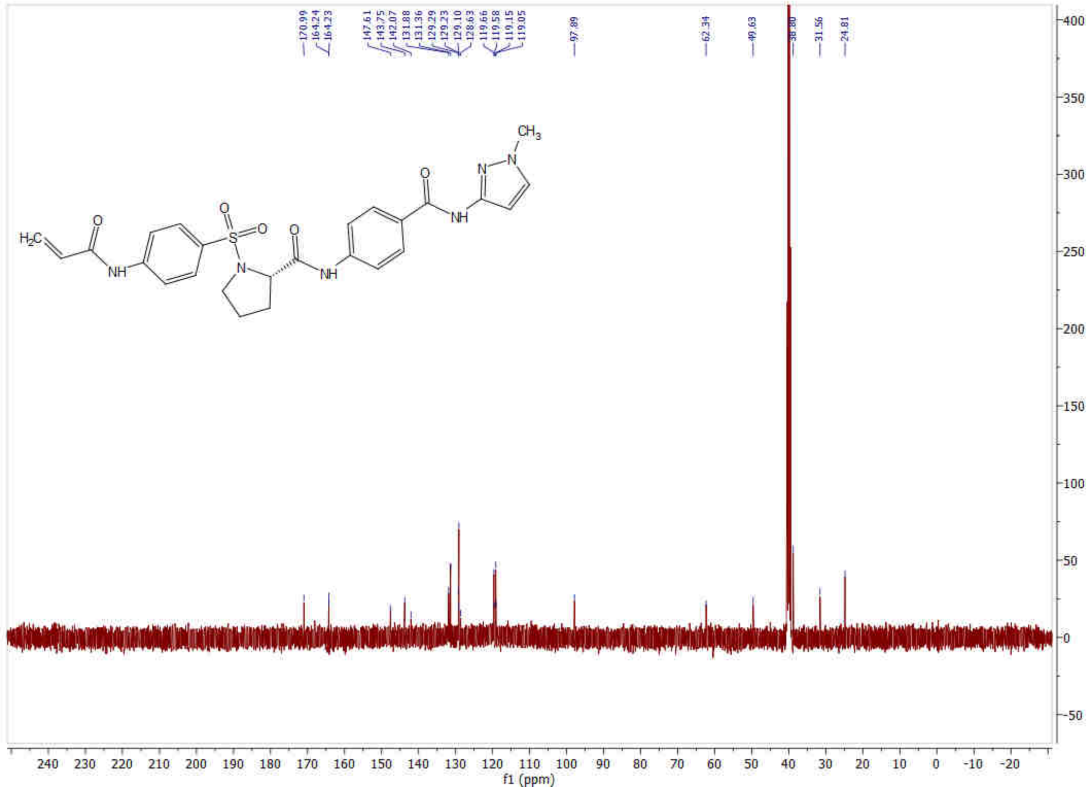

**(*R*)-1-((4-Acrylamidophenyl)sulfonyl)-*N*-(4-((1-methyl-1H-pyrazol-3-yl)carbamoyl)phenyl)pyrrolidine-2-carboxamide (11)**

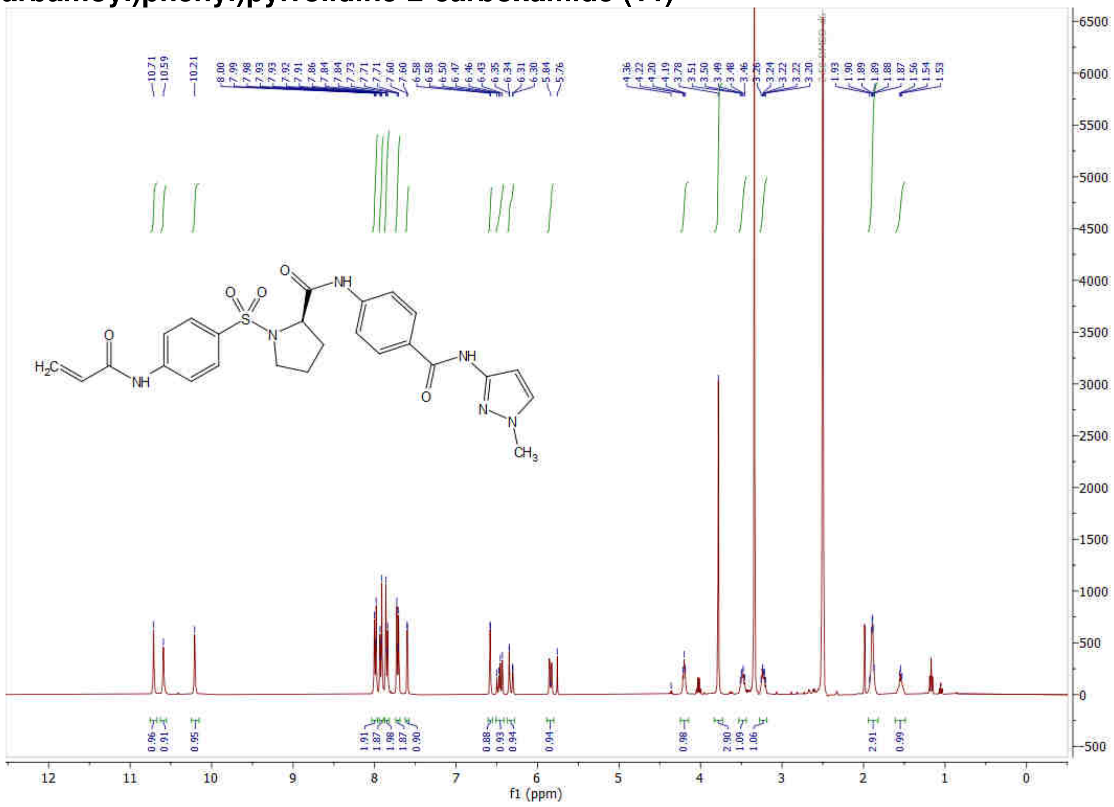

**(*S*)-1-((4-Acrylamidophenyl)sulfonyl)-*N*-(4-((1-(prop-2-yn-1-yl)-1H-pyrazol-4-yl)carbamoyl)phenyl)pyrrolidine-2-carboxamide (12, IMP-2566)**

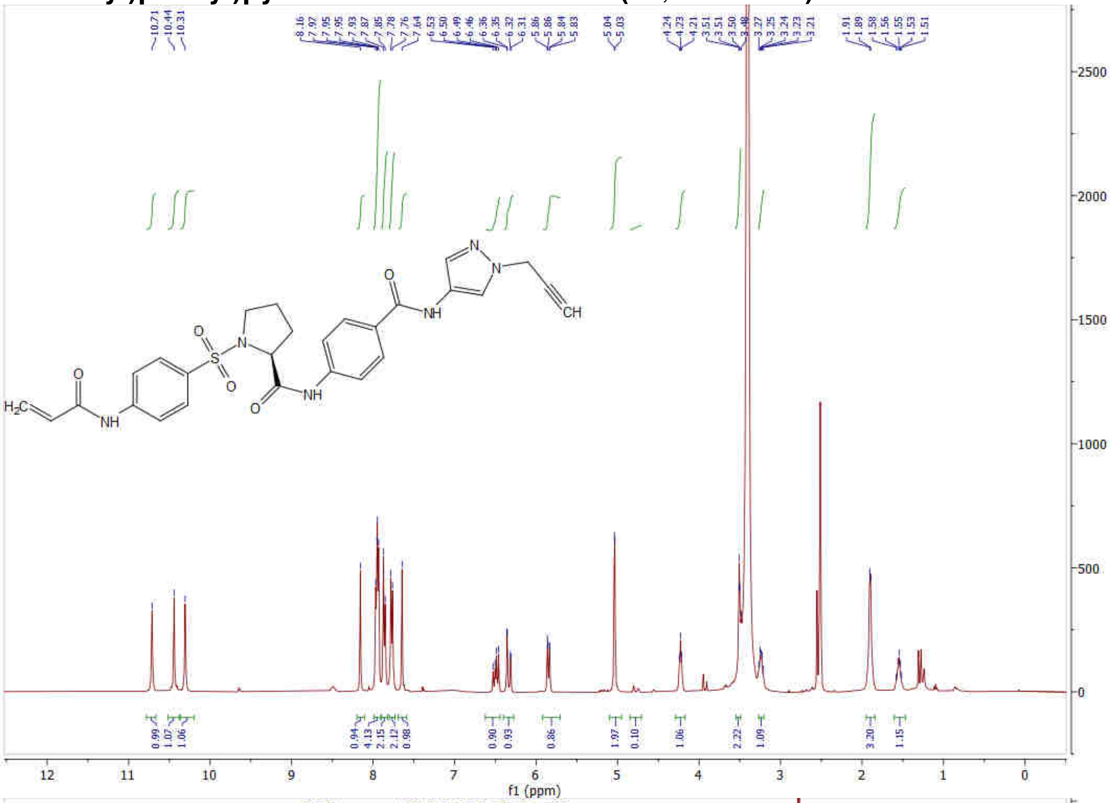

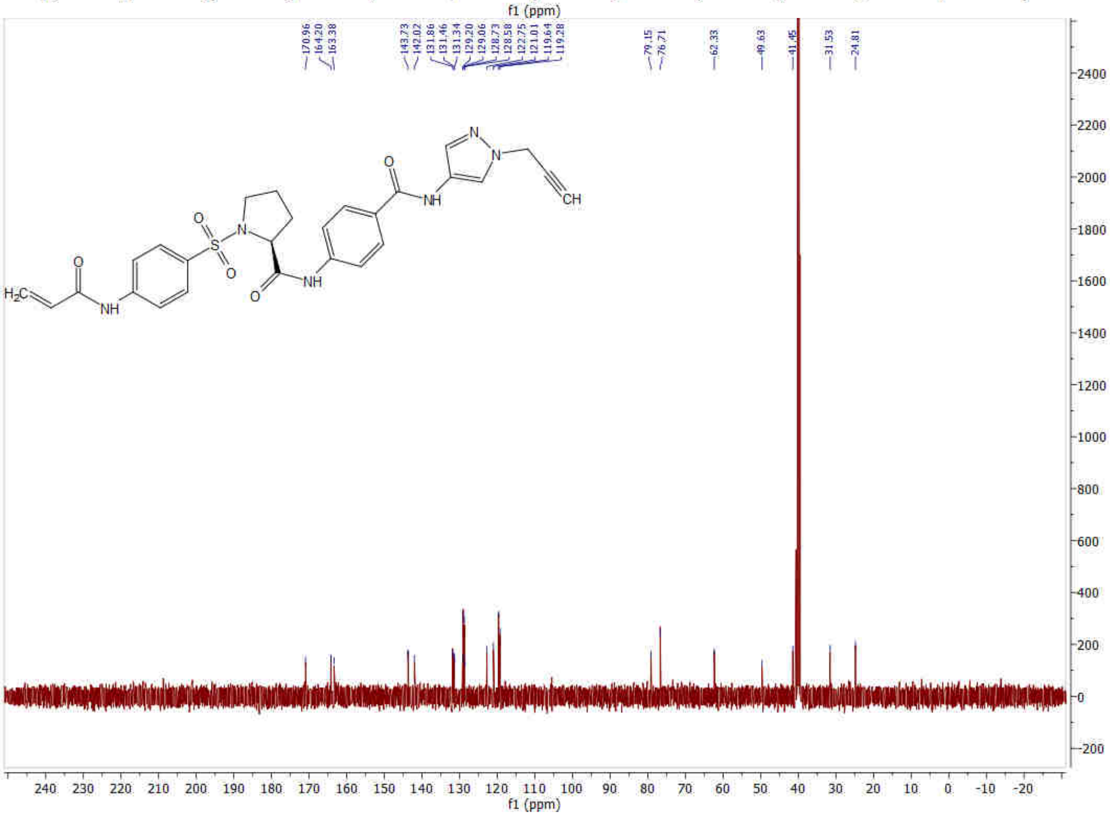

**(*S*)-1-((4-Acrylamidophenyl)sulfonyl)-*N*-(3-methoxyphenyl)pyrrolidine-2-carboxamide (13, IMP-2505)**

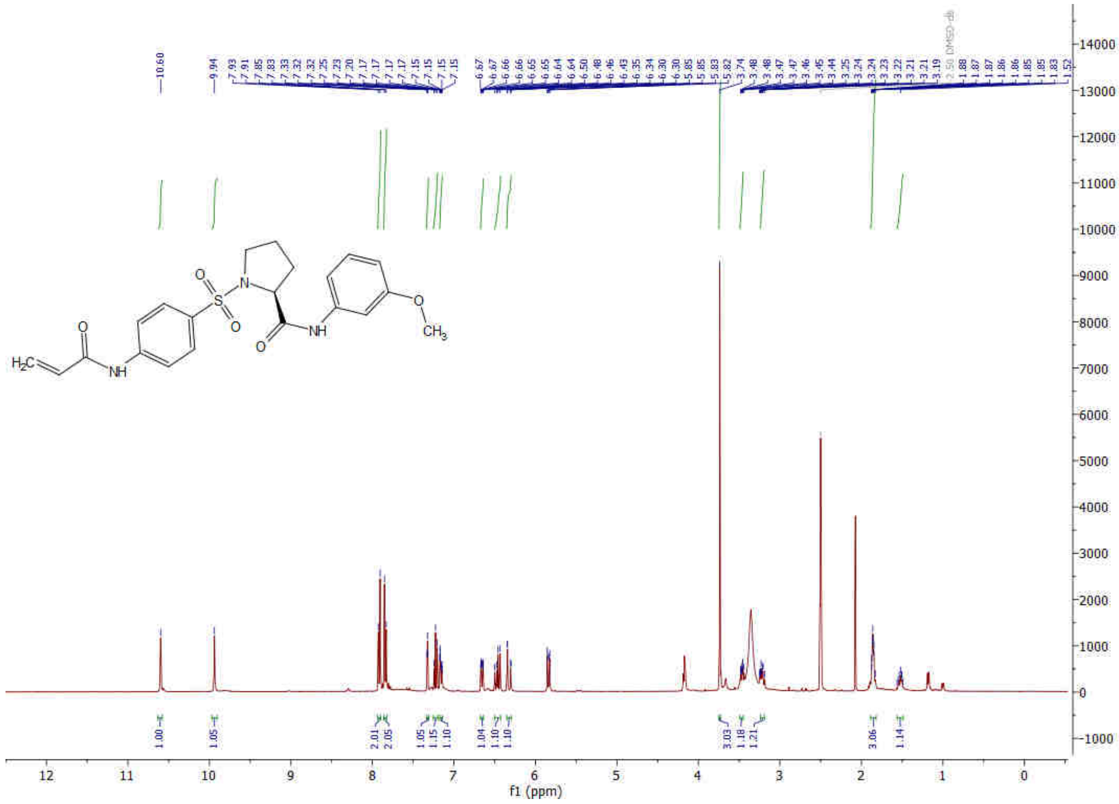

### HPLC Traces

#### *N*-(4-(Pyrrolidin-1-ylsulfonyl)phenyl)acrylamide (1, IMP-1704)

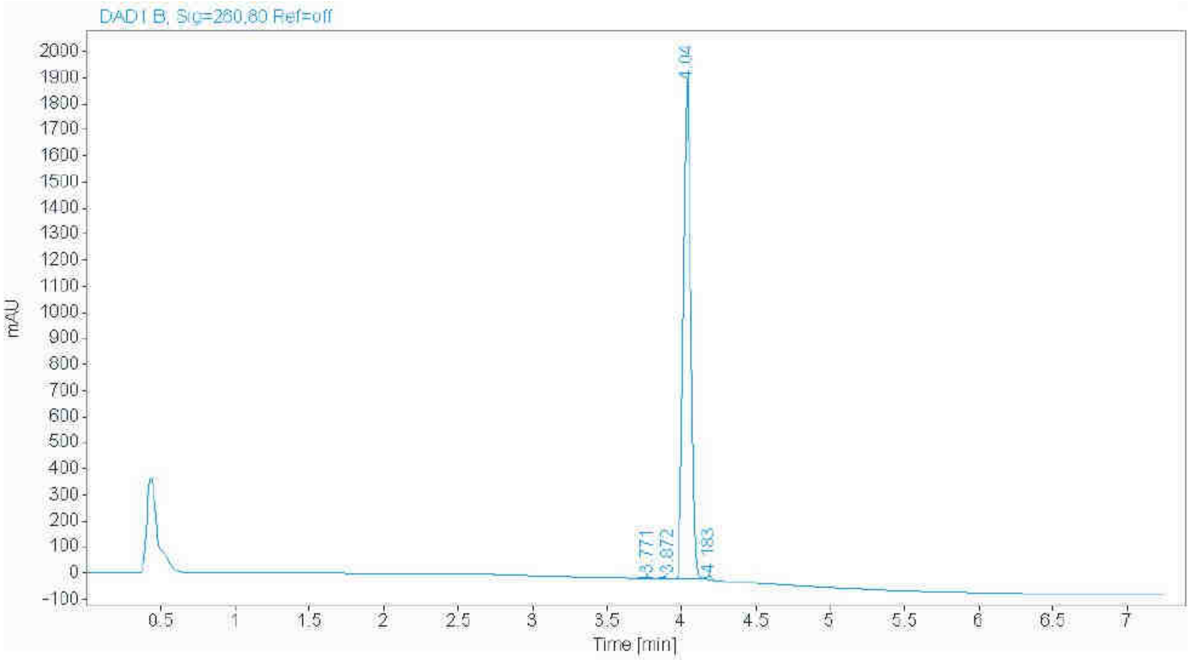


| RT [min] | Width [min] | Area | Height | Area% |
|---|---|---|---|---|
| 3.771 | 0.0572 | 12.8330 | 3.4617 | 0.1859 |
| 3.872 | 0.0503 | 9.9700 | 3.2077 | 0.1444 |
| 4.040 | 0.0577 | 6848.6528 | 1913.8254 | 99.2228 |
| 4.183 | 0.0429 | 30.8401 | 11.6413 | 0.4468 |
| | Sum | 6902.2959 | | |

#### *N*-(4-(Pyrrolidine-1-carbonyl)phenyl)acrylamide (2)

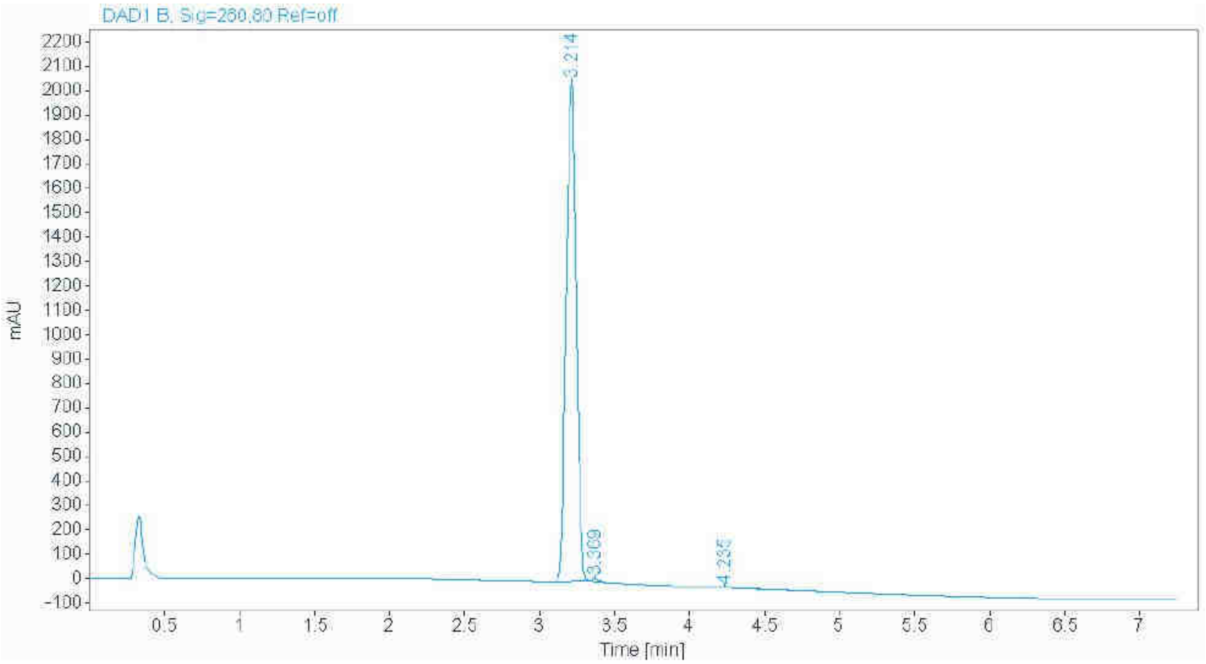


| RT [min] | Width [min] | Area | Height | Area% |
|---|---|---|---|---|
| 3.214 | 0.0750 | 9658.4990 | 2049.9414 | 99.4610 |
| 3.369 | 0.0475 | 29.8351 | 9.8111 | 0.3072 |
| 4.235 | 0.1254 | 22.5057 | 2.3158 | 0.2318 |
| | Sum | 9710.8398 | | |

***N*-(3-(Pyrrolidin-1-ylsulfonyl)phenyl)acrylamide (3)**

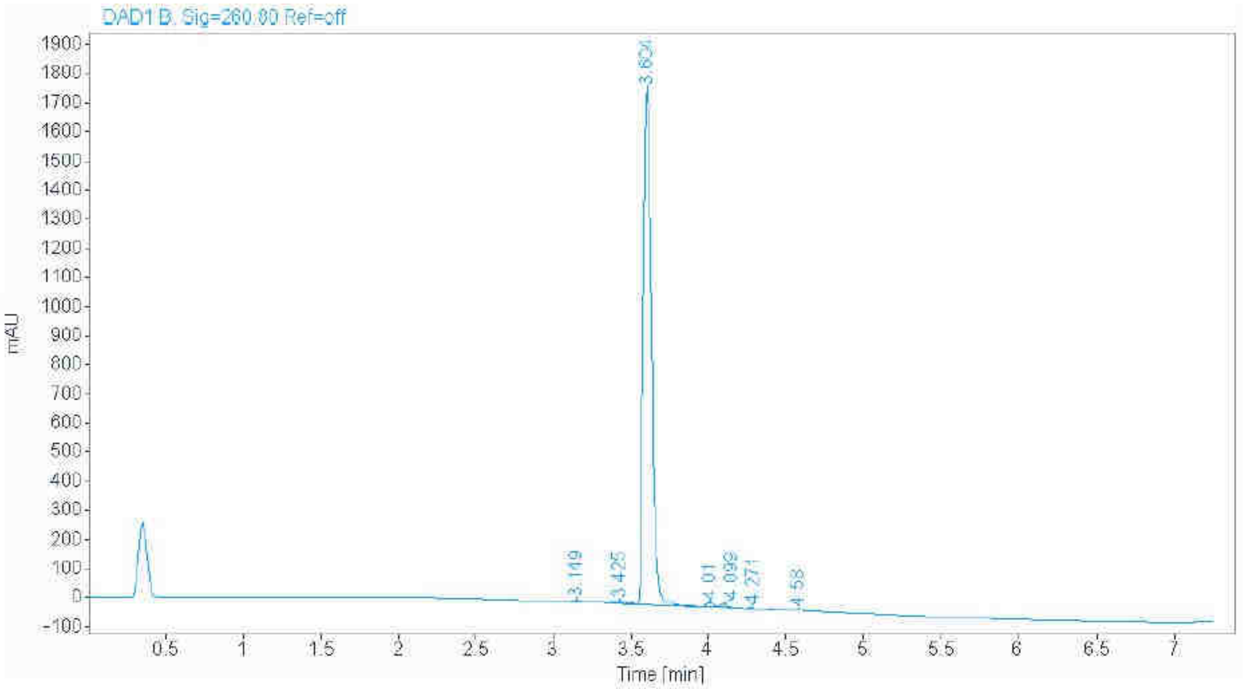


| RT [min] | Width [min] | Area | Height | Area% |
|---|---|---|---|---|
| 3.149 | 0.0594 | 10.3622 | 2.5528 | 0.1588 |
| 3.425 | 0.0845 | 16.5179 | 3.1858 | 0.2531 |
| 3.604 | 0.0539 | 6406.0527 | 1782.8141 | 98.1778 |
| 4.010 | 0.0449 | 32.4232 | 11.5093 | 0.4969 |
| 4.099 | 0.0485 | 31.4536 | 10.6448 | 0.4821 |
| 4.271 | 0.1560 | 15.8032 | 1.3554 | 0.2422 |
| 4.580 | 0.0538 | 12.3349 | 3.4389 | 0.1890 |
| | Sum | 6524.9478 | | |

**Methyl ((4-acrylamidophenyl)sulfonyl)-*L*-prolinate (4, IMP1712)**

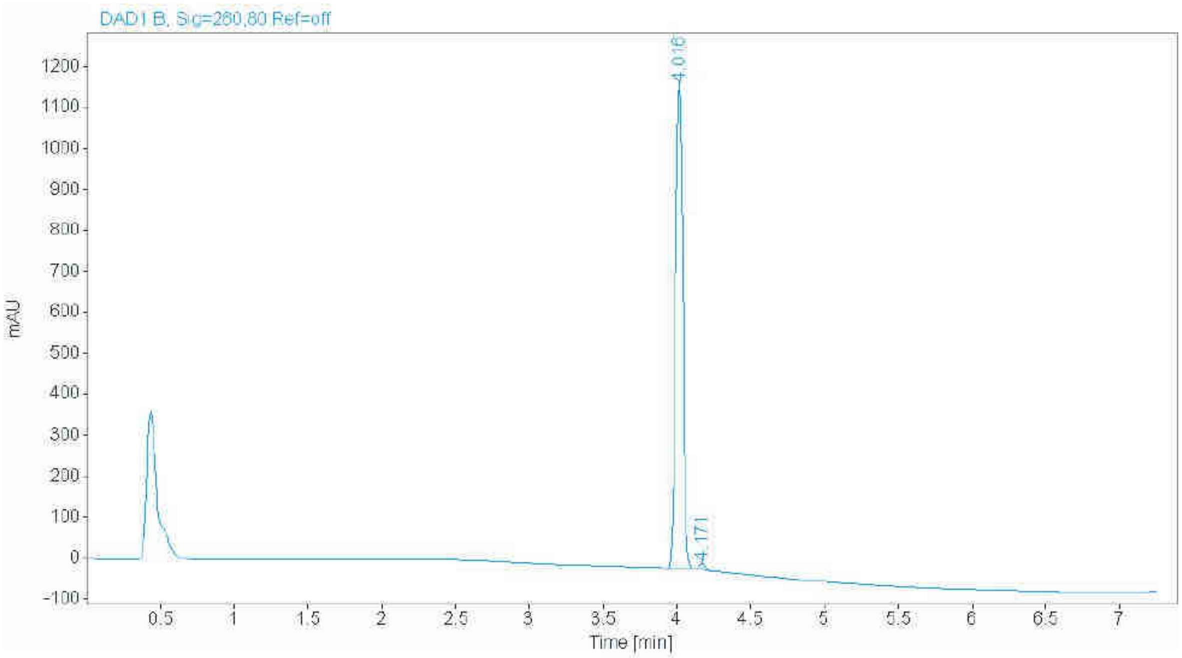


| RT [min] | Width [min] | Area | Height | Area% |
|---|---|---|---|---|
| 4.016 | 0.0529 | 3951.4736 | 1185.1375 | 98.8760 |
| 4.171 | 0.0471 | 44.9209 | 14.9492 | 1.1240 |
| | Sum | 3996.3945 | | |

**Methyl ((4-acrylamidophenyl)sulfonyl)-*D*-prolinate (5)**

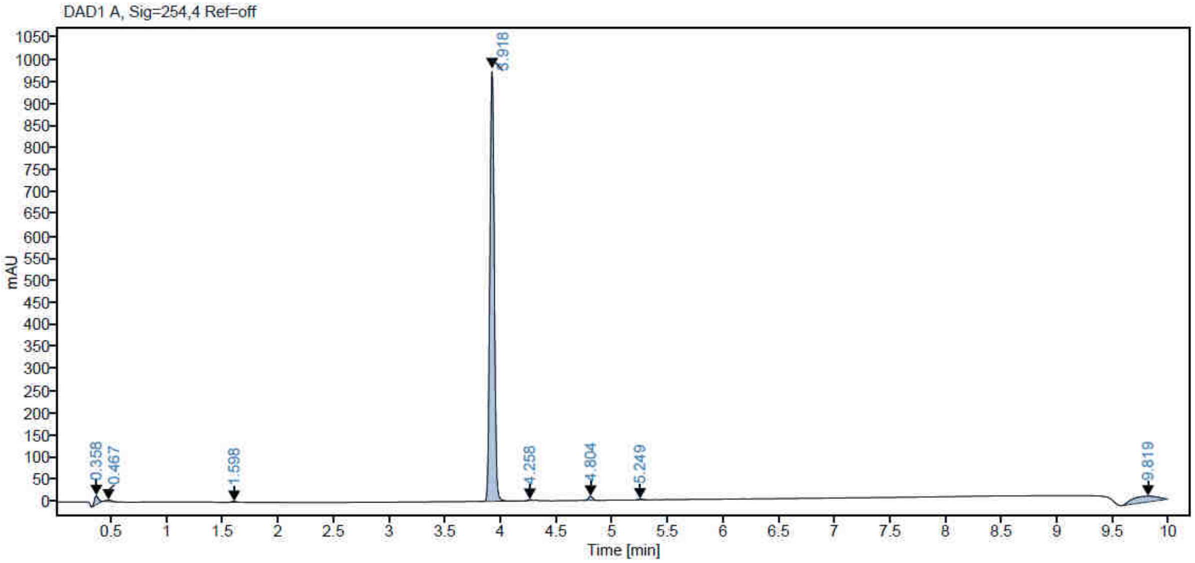


| RT [min] | Width [min] | Area | Height | Area% |
|---|---|---|---|---|
| 1.598 | 0.0946 | 15.5108 | 2.3634 | 0.557 |
| 3.918 | 0.0446 | 2731.1531 | 978.4450 | 98.121 |
| 4.258 | 0.0375 | 5.4588 | 2.3184 | 0.196 |
| 4.804 | 0.0447 | 25.6539 | 9.1610 | 0.922 |
| 5.249 | 0.0467 | 5.6877 | 1.9134 | 0.204 |
| **Sum** | | 2783.4643 | | |

**(*S*)-1-((4-Acrylamidophenyl)sulfonyl)-*N*-(prop-2-yn-1-yl)pyrrolidine-2-carboxamide (6)**

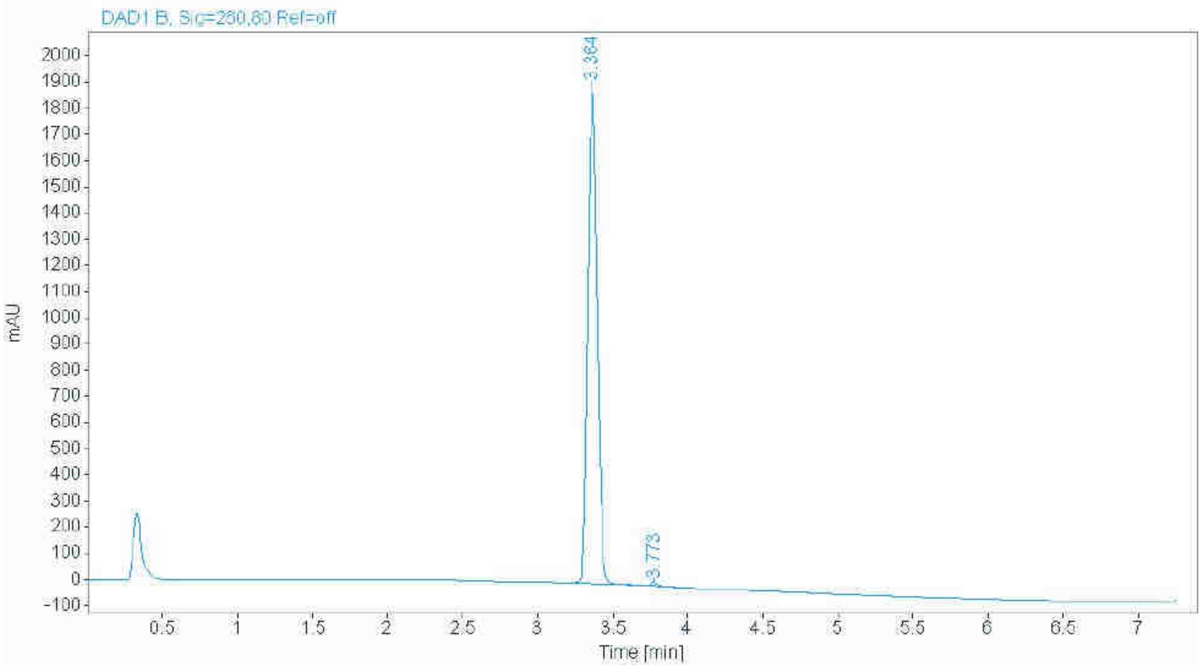


| RT [min] | Width [min] | Area | Height | Area% |
|---|---|---|---|---|
| 3.364 | 0.0691 | 8398.2217 | 1918.0364 | 99.3993 |
| 3.773 | 0.0527 | 50.7556 | 14.5393 | 0.6007 |
| | Sum | 8448.9773 | | |

**(*S*)-1-((4-Acrylamidophenyl)sulfonyl)-*N*-(4-methoxyphenyl)pyrrolidine-2-carboxamide (7)**

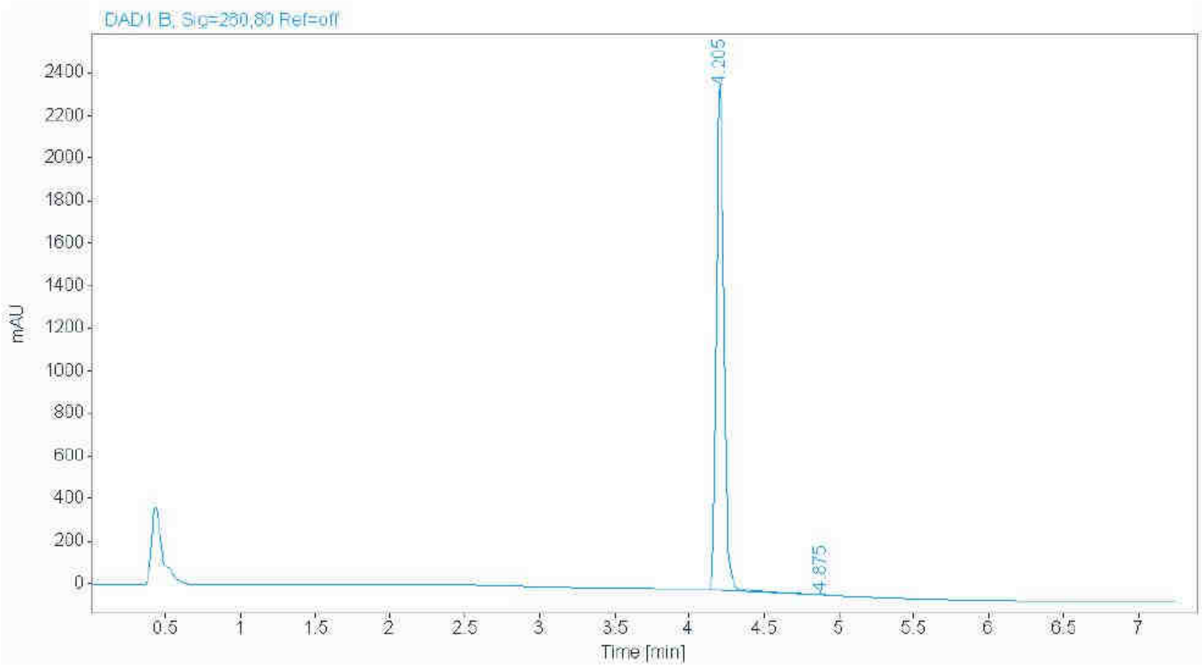


| RT [min] | Width [min] | Area | Height | Area% |
|---|---|---|---|---|
| 4.205 | 0.0583 | 8623.8184 | 2378.1069 | 99.9026 |
| 4.875 | 0.0460 | 8.4121 | 2.8891 | 0.0974 |
| | Sum | 8632.2304 | | |

**(*R*)-1-((4-Acrylamidophenyl)sulfonyl)-*N*-(4-methoxyphenyl)pyrrolidine-2-carboxamide (8)**

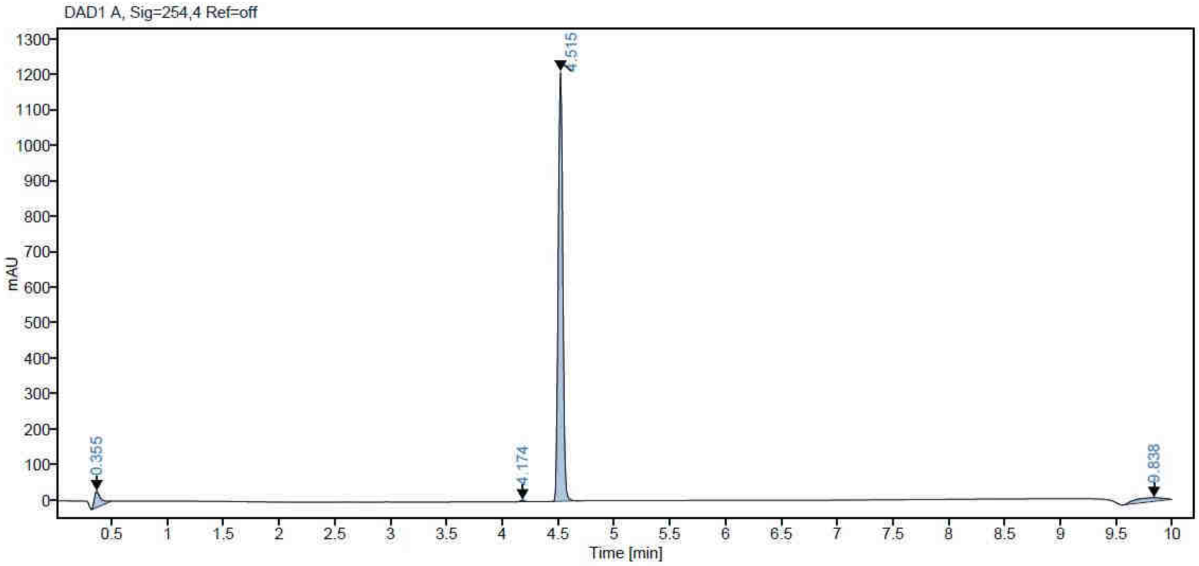


| RT [min] | Width [min] | Area | Height | Area% |
|---|---|---|---|---|
| 4.174 | 0.0439 | 12.3452 | 4.5113 | 0.358 |
| 4.515 | 0.0451 | 3433.5769 | 1210.5248 | 99.642 |
| **Sum** | | 3445.922 | | |

**(*S*)-1-((4-Acrylamidophenyl)sulfonyl)-*N*-(4-(prop-2-yn-1-yloxy)phenyl)pyrrolidine-2-carboxamide (9)**

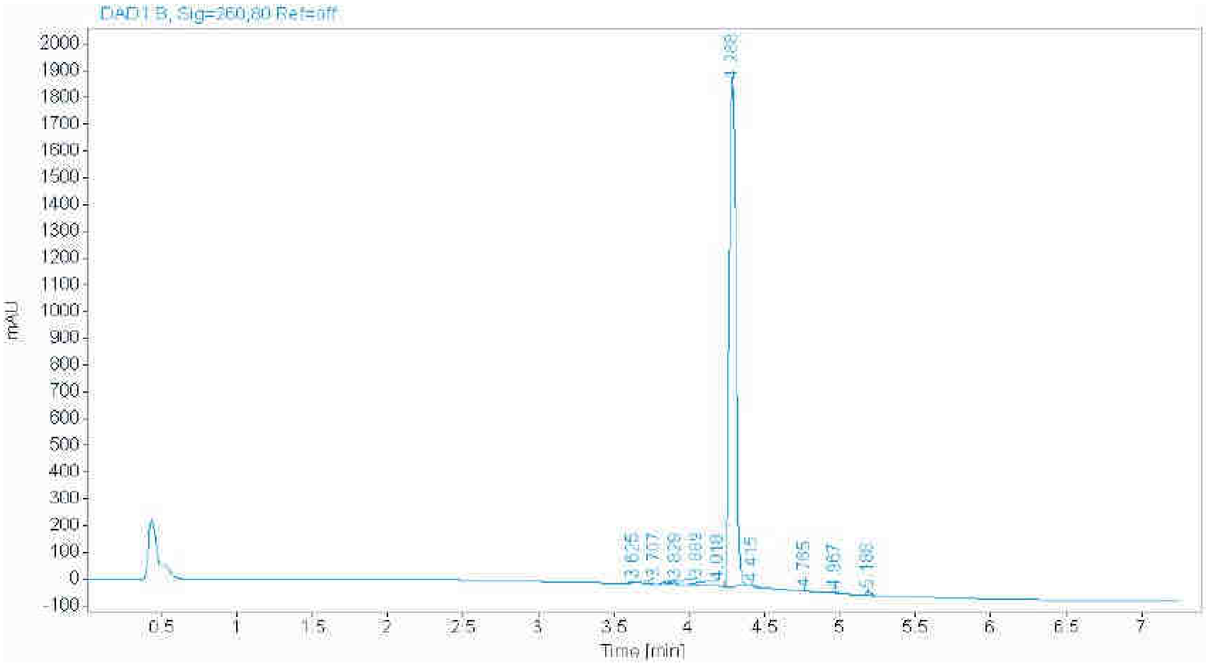


| RT [min] | Width [min] | Area | Height | Area% |
|---|---|---|---|---|
| 3.625 | 0.0428 | 7.8136 | 2.9598 | 0.1355 |
| 3.707 | 0.0489 | 28.6767 | 9.5870 | 0.4975 |
| 3.829 | 0.0350 | 7.1071 | 3.3191 | 0.1233 |
| 3.889 | 0.0414 | 18.1426 | 7.1913 | 0.3147 |
| 4.018 | 0.0471 | 21.1688 | 7.0282 | 0.3672 |
| 4.288 | 0.0487 | 5632.6465 | 1892.5148 | 97.7093 |
| 4.415 | 0.0444 | 6.8031 | 2.4480 | 0.1180 |
| 4.765 | 0.0421 | 9.6745 | 3.7456 | 0.1678 |
| 4.967 | 0.0520 | 8.4660 | 2.5991 | 0.1469 |
| 5.188 | 0.0444 | 24.1971 | 8.7102 | 0.4197 |
| | Sum | 5764.6960 | | |

**(*S*)-1-((4-Acrylamidophenyl)sulfonyl)-*N*-(4-((1-methyl-1H-pyrazol-3-yl)carbamoyl)phenyl)pyrrolidine-2-carboxamide (10, IMP-2552)**

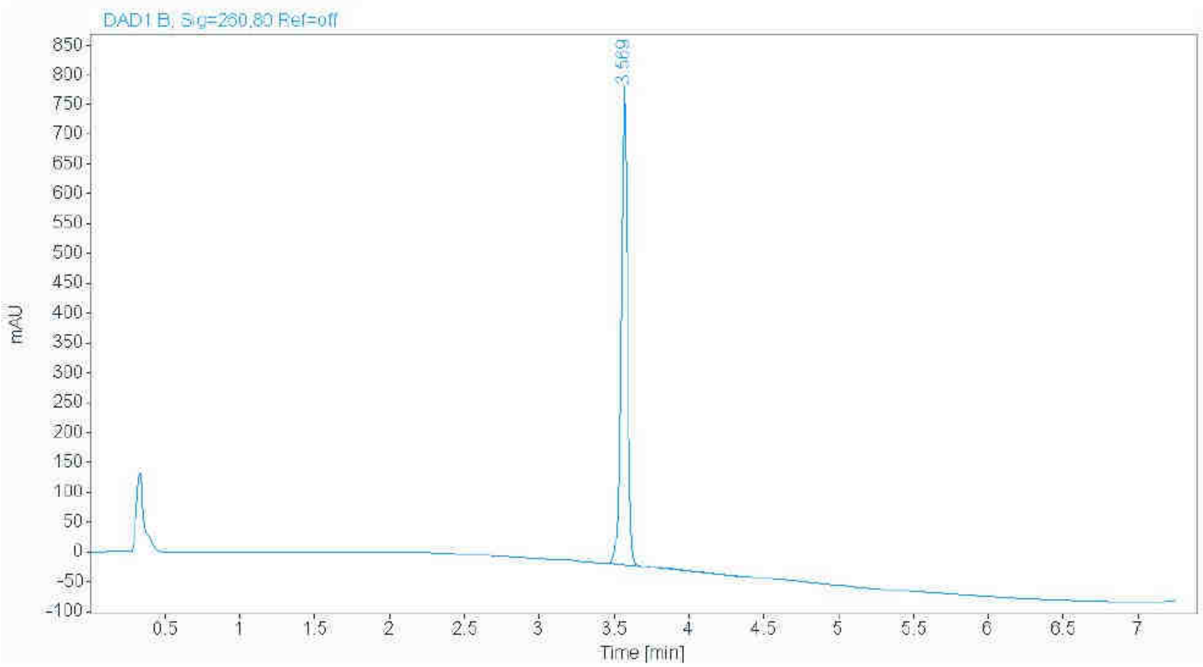


| RT [min] | Width [min] | Area | Height | Area% |
|---|---|---|---|---|
| 3.569 | 0.0465 | 2379.6851 | 804.5959 | 100.0000 |
| | Sum | 2379.6851 | | |

**(*R*)-1-((4-Acrylamidophenyl)sulfonyl)-*N*-(4-((1-methyl-1H-pyrazol-3-yl)carbamoyl)phenyl)pyrrolidine-2-carboxamide (11)**

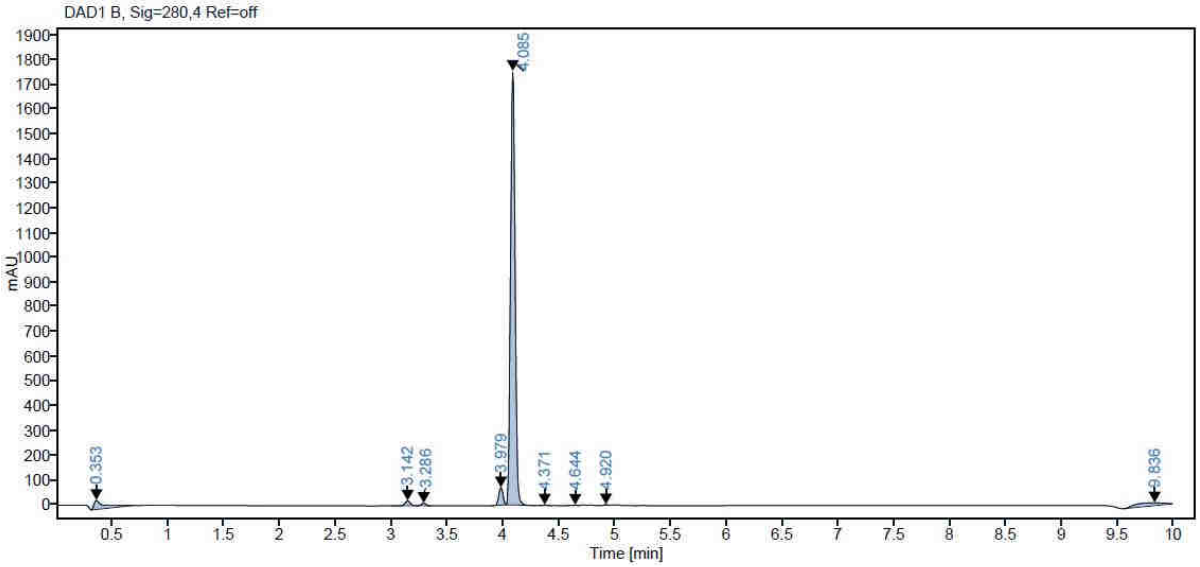


| RT [min] | Width [min] | Area | Height | Area% |
|---|---|---|---|---|
| 3.142 | 0.0529 | 64.7917 | 19.4264 | 1.217 |
| 3.286 | 0.0453 | 28.8753 | 10.1085 | 0.542 |
| 3.979 | 0.0426 | 185.5625 | 70.7643 | 3.484 |
| 4.085 | 0.0454 | 5026.905 | 1753.59 | 94.383 |
| 4.371 | 0.0421 | 6.7368 | 2.6095 | 0.126 |
| 4.644 | 0.0583 | 6.5666 | 1.6549 | 0.123 |
| 4.92 | 0.0423 | 6.6206 | 2.5492 | 0.124 |
| **Sum** | | 5326.0583 | | |

**(*S*)-1-((4-Acrylamidophenyl)sulfonyl)-*N*-(4-((1-(prop-2-yn-1-yl)-1H-pyrazol-4-yl)carbamoyl)phenyl)pyrrolidine-2-carboxamide (12, IMP-2566)**

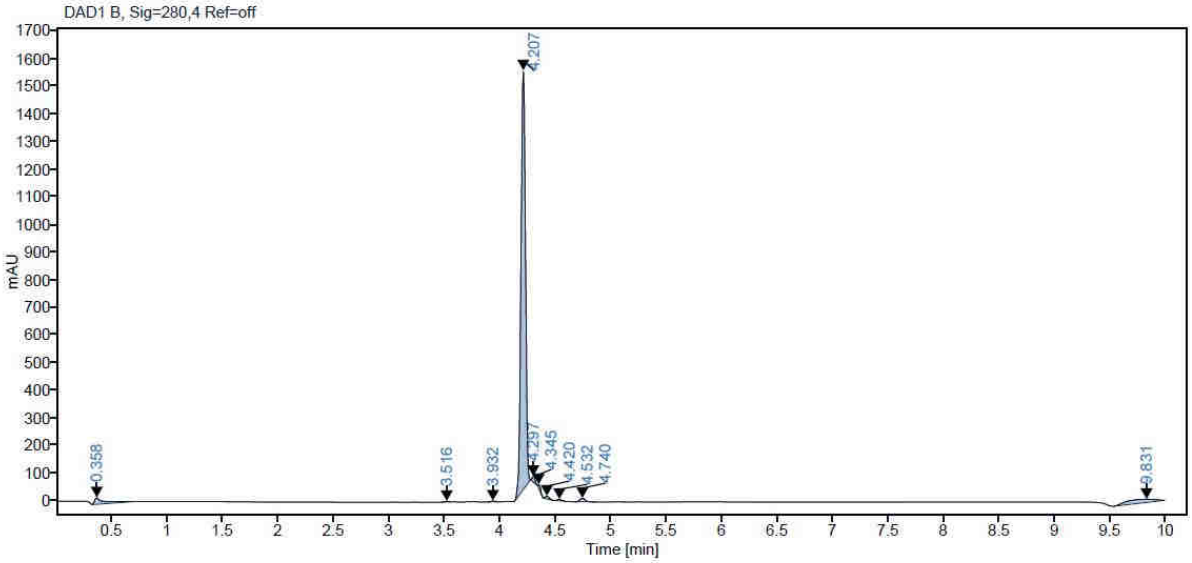


| RT [min] | Width [min] | Area | Height | Area% |
|---|---|---|---|---|
| 3.516 | 0.0401 | 6.1244 | 2.5453 | 0.140 |
| 3.932 | 0.0421 | 9.0759 | 3.5205 | 0.208 |
| 4.207 | 0.0444 | 4206.396 | 1514.299 | 96.242 |
| 4.297 | 0.0293 | 37.6807 | 20.6459 | 0.862 |
| 4.345 | 0.0351 | 28.3177 | 13.4499 | 0.648 |
| 4.42 | 0.0375 | 24.54 | 10.4237 | 0.561 |
| 4.532 | 0.0447 | 14.7185 | 5.2542 | 0.337 |
| 4.74 | 0.0529 | 43.7828 | 13.137 | 1.002 |
| **Sum** | | 4370.636 | | |

**(*S*)-1-((4-Acrylamidophenyl)sulfonyl)-*N*-(3-methoxyphenyl)pyrrolidine-2-carboxamide (13, IMP-2505)**

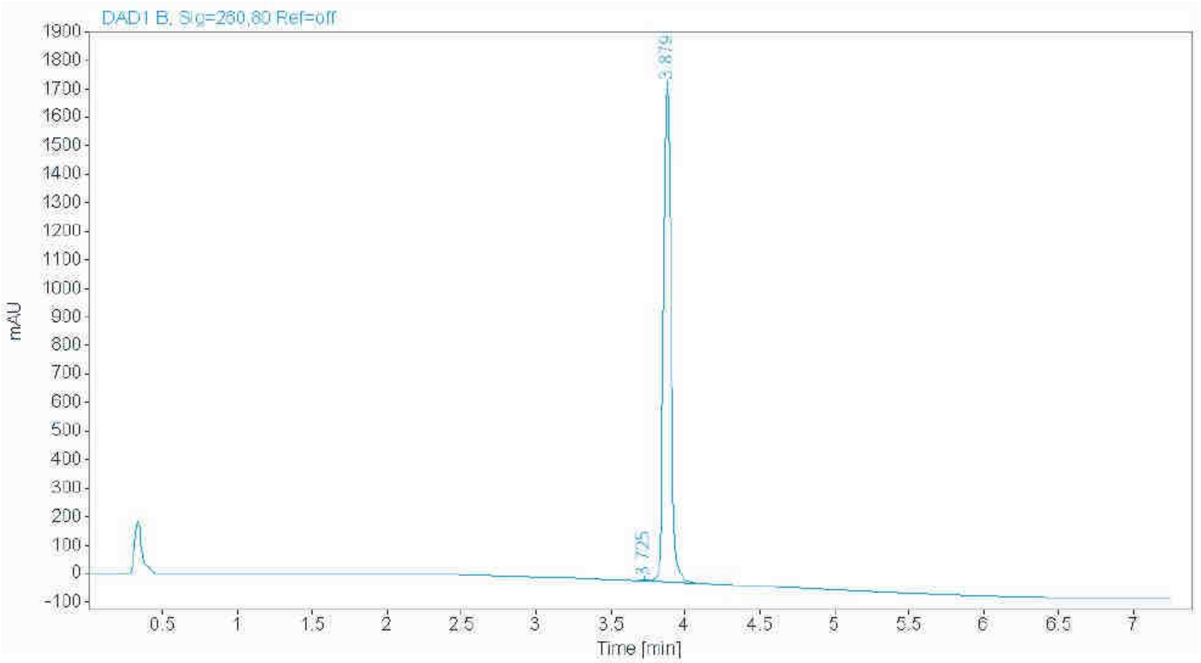


| RT [min] | Width [min] | Area | Height | Area% |
|---|---|---|---|---|
| 3.725 | 0.0420 | 9.9573 | 3.8667 | 0.1740 |
| 3.879 | 0.0521 | 5711.0508 | 1746.7648 | 99.8260 |
| | Sum | 5721.0081 | | |
